# Development of potent dual BET/HDAC inhibitors *via* pharmacophore merging and structure-guided optimization

**DOI:** 10.1101/2023.07.18.547334

**Authors:** Nicolas Bauer, Dimitrios-Ilias Balourdas, Joel R. Schneider, Xin Zhang, Lena M. Berger, Benedict-Tilman Berger, Nick A. Klopp, Jens T. Siveke, Stefan Knapp, Andreas C. Joerger

**Affiliations:** Institute of Pharmaceutical Chemistry, Goethe University, Max-von-Laue-Str. 9, 60438 Frankfurt am Main, Germany; Structural Genomics Consortium (SGC), Buchmann Institute for Life Sciences, Max-von-Laue-Str. 15, 60438 Frankfurt am Main, Germany; Bridge Institute of Experimental Tumor Therapy, West German Cancer Center, University Hospital Essen, University of Duisburg-Essen, Essen, Germany; Division of Solid Tumor Translational Oncology, German Cancer Consortium (DKTK Partner Site Essen) and German Cancer Research Center, DKFZ, Heidelberg, Germany; German translational cancer network (DKTK) site Frankfurt/Mainz

## Abstract

Bromodomain and extra-terminal motif (BET) proteins and histone deacetylases (HDACs) are prime targets in cancer therapy. Recent research has particularly focused on the development of dual BET/HDAC inhibitors for hard-to-treat tumors such as pancreatic cancer. Here, we have developed a new series of potent dual BET/HDAC inhibitors by choosing starting scaffolds that enabled us to optimally merge the two functionalities into a single compound. Systematic structure-guided modification of both warheads then led to optimized binders that were superior in potency to both parent compounds, with the best molecules of this series binding to both BRD4 bromodomains as well as HDAC1/2 with EC_50_ values in the 100-nanomolar range in cellular NanoBRET target engagement assays. Importantly, this on-target activity also translated into promising efficacy in pancreatic cancer and NUT midline carcinoma cells. Our lead molecules effectively blocked histone H3 deacetylation in pancreatic cancer cells and upregulated the tumor suppressor *HEXIM1* and proapoptotic *p57*, both markers of BET inhibition. In addition, they have the potential to downregulate oncogenic drivers of NUT midline carcinoma, as demonstrated for *MYC* and *TP63* mRNA levels. Overall, this study expands the portfolio of available dual BET/class I HDAC inhibitors for future translational studies in different cancer models.

## INTRODUCTION

Epigenetic alterations modifying the chromatin structure and resulting in aberrant gene transcription play a crucial role in cancer development [1]. Due to the reversibility of epigenetic modifications, chromatin-interacting proteins have emerged as attractive targets for the treatment of cancer [2]. Many epigenetic drugs have been approved or are currently in clinical trials, including histone deacetylase (HDAC) and bromodomain and extra-terminal (BET) inhibitors [2–7]. A combination of BET and HDAC inhibitors, for example, has been suggested for the treatment of pancreatic ductal adenocarcinoma (PDAC) [8], which is one of the most lethal human cancers that is resistant to virtually all therapeutic approaches [9]. Combination therapy is a common strategy to prevent compensatory mechanisms such as drug resistance [10]. The use of multiple drugs, however, has certain limitations due to possible drug-drug interactions and different biodistribution or pharmacokinetic profiles. By employing multi-target drugs, these issues can be avoided while additionally leading to simpler regulatory processes and possibly better patient compliance.

Several dual BET/HDAC inhibitors have been developed based on BET inhibitors (+)-JQ1 [11–13], RVX-208 [14], ABBV-744 [15], I-BET295 [16], I-BET762 [17], and other inhibitor scaffolds [18–23]. As an HDAC inhibiting moiety, most dual BET/HDAC inhibitors are based on suberoyl anilide hydroxamic acid (SAHA), a pan-HDAC inhibitor with a relatively short metabolic half-life of 2 h [24]. These dual BET/HDAC inhibitors showed promising antitumor effects in different types of cancer cell lines, including leukemia [14, 18, 20], colorectal carcinoma [21], NUT midline carcinoma (NMC) [12, 16], and PDAC cells [11, 12]. We have, for example, developed TW9, an adduct of the BET inhibitor (+)-JQ1 and class I HDAC inhibitor CI-994, which was more potent in inhibiting proliferation of PDAC cells than its parental molecules (+)-JQ1 or CI-994 alone, or combined treatment of both inhibitors [12]. Gene expression profiling showed that the antitumor effects of TW9 correlate with a dysregulation of a FOSL1-directed transcriptional program [12] and upregulation of proapoptotic genes BIM, NOXA, PUMA, and BMF [25].

The human genome encodes for 18 different HDACs, which are grouped into four different classes based on sequence homology to yeast HDACs and domain organization. Class I, II, and IV HDACs are zinc-dependent, whereas class III HDACs, the so-called sirtuins, require NAD^+^ as a cofactor [26, 27]. The class I HDAC family comprises HDAC1-3 and HDAC8. HDAC1-3 are located primarily in the nucleus where they are involved in histone deacetylation and are recruited to large multi-protein complexes involved in nucleosome remodeling [26, 27]. HDAC1 and 2, for example, are recruited to the nucleosome remodeling and deacetylase complex (NuRD), the transcriptional corepressor Sin3A, corepressor of REST (CoREST), and the mitotic deacetylase complex (MiDAC) [28–31], while HDAC3 associates with the SMRT/N-CoR corepressor complex [32, 33], highlighting the crucial role of these HDAC family members in transcriptional regulation and gene silencing.

As pointed out above, most available dual BET/HDAC inhibitors contain a pan-HDAC hydroxamic acid warhead. The few published dual inhibitors with a class I specific HDAC warhead are relatively large molecules with a molecular weight of over 600 Da [12, 15], limiting application *in vivo* due to poor pharmacokinetic properties. In order to overcome limitations of simple adduct formation of two available inhibitors targeting either HDAC or BET proteins, we aimed to develop an integrated dual BET/HDAC inhibitor. We merged BRD4 and class I HDAC pharmacophores, resulting in dual inhibitors of reduced molecular weight with a core scaffold of only 350 Da harboring both inhibitor activities. The starting molecule was designed by merging the scaffolds of bromodomain inhibitor MS436 and HDAC inhibitor CI-944 with a class I HDAC selective benzamide moiety [34]. Subsequent synthetic efforts and structure-guided design resulted in dual inhibitors with optimized warheads, binding to both bromodomains of BRD4 as well as HDAC1/2 with affinities in the 100 nM range. Preliminary studies in cancer cells revealed promising biological activities in PDAC and NMC cells.

## RESULTS AND DISCUSSION

### Strategy for dual BET/HDAC inhibitor development

The initial dual BET/HDAC inhibitor, NB161 (**1**), was developed by merging BET inhibitor MS436 with class I HDAC inhibitor CI-994. BD1-selective inhibitor MS436 was chosen as a starting scaffold because it appeared ideal for incorporating the zinc-binding moiety of HDAC inhibitor CI-994 due to structural overlap (**Figure 1A**). NB161 was synthesized *via* diazotization and subsequent azo coupling of aniline **5** and 5-amino-2-methylphenol with isoamyl nitrite. Aniline **5** was prepared by amide coupling and subsequent deprotection of *Boc*-protected *o*-phenylenediamine **2** and *Fmoc*-protected *p*-aminobenzoic acid **3** (**Scheme 1**). Binding of NB161 to BRD4 was assessed by thermal shift assays using differential scanning fluorimetry (DSF), which measures the increase in the melting temperature (*T*_m_) of the protein upon ligand binding, which for a given protein correlates with the affinity of the ligand. For the first bromodomain of BRD4 (BD1), NB161 induced a *T*_m_ shift, Δ*T*_m_, of 5.3 K, which was significantly higher than that for the parent molecule MS436 (Δ*T*_m_ = 4.0 K) (**Table 1**). The *T*_m_ shift for the second bromodomain (BD2) was almost identical for both inhibitors (Δ*T*_m_ ≍ 3 K), indicating that binding was not compromised upon modification and even slightly improved for BD1. We then determined a high-resolution crystal structure of BRD4 BD1 in complex with NB161 to elucidate its binding mode and provide a structural framework for inhibitor optimization (**Figure 1B**). The binding mode in the acetyl-lysine pocket of BD1 overlapped with that of MS436, including the hydrogen bond with the side chain of the conserved Asn140. The diazo moiety of NB161 interacted with a structural water molecule, and the phenyl linker was sandwiched between Leu92 and Trp81 of the WPF-shelf region. The HDAC-binding moiety made no specific interactions with the protein and protruded into the solvent.

**Figure 1.**
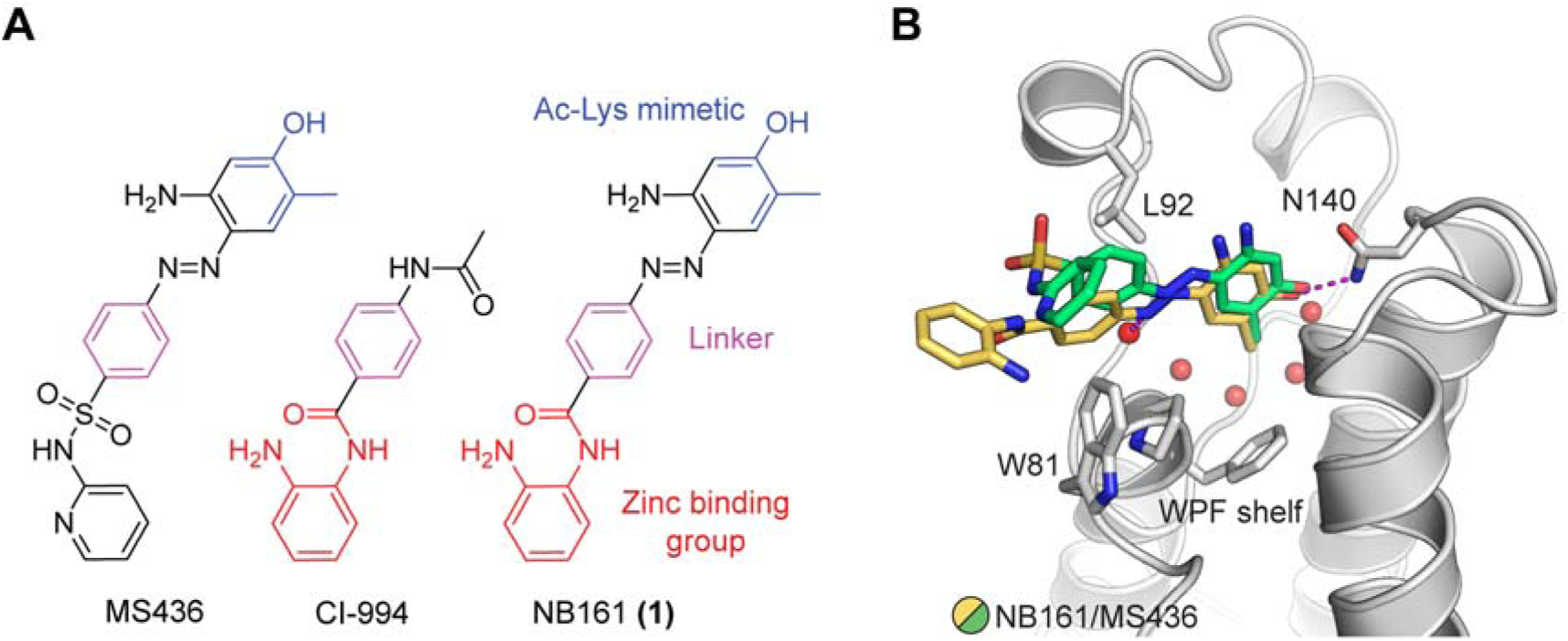
Dual inhibitor design strategy. (A) Merging of pharmacophores. (B) Binding mode of the first-generation dual inhibitor NB161 (yellow stick model) in BRD4 BD1 (grey ribbon diagram) superimposed onto that of the parent BET inhibitor MS436 (green stick model, PDB entry 4nud). For clarity, only the protein for the NB161 complex is shown. Key interacting side chains are highlighted as stick models, and selected structural water molecules in the binding pocket are shown as red spheres.

**Scheme 1.**
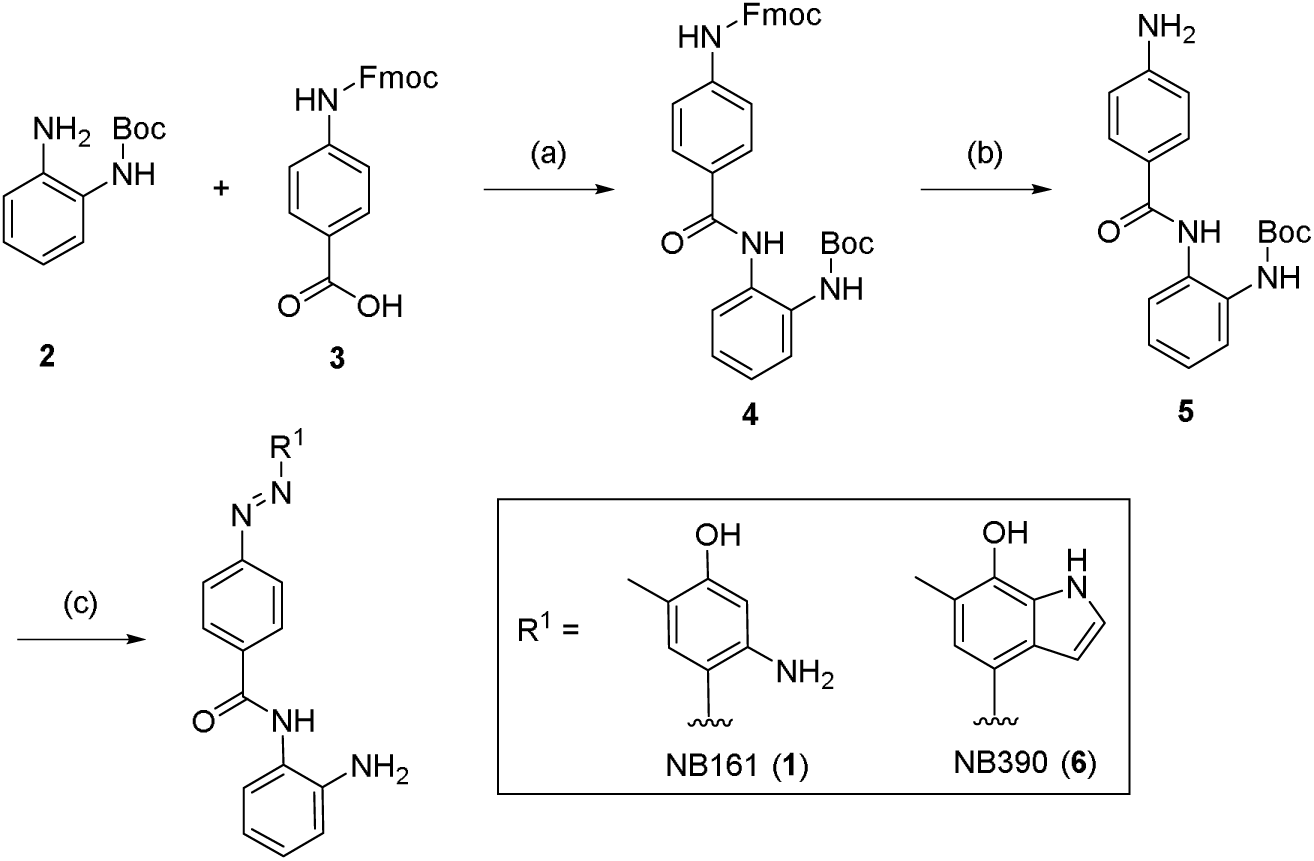
Synthesis of inhibitors NB161 and NB390. Reagents and conditions: (a) PyAOP, DIPEA, DMF, rt, 16 h; (b) morpholine/ACN, rt, 2 h; (c) (1) (i) conc. HCl, isoamyl nitrite, MeOH/ACN, - 10 °C, 1 h; (ii) 5-amino-2-methylphenol or 6-methyl-1*H*-indol-7-ol, K_2_CO_3_, MeOH/H_2_O/ACN, −10 C to rt, 2 h; (2) TFA/DCM, rt, 1 h. The 7-hydroxyindol moiety was synthesized following published protocols [35, 36].

**Table 1.** Optimization of the Asn140 binding moiety and replacement of the diazo moiety.

| Compounds | DSF $\Delta T_m$ (K) | | NanoBRET EC<sub>50</sub> (μM) | | | |
| --- | --- | --- | --- | --- | --- | --- |
|  | BRD4-BD1 | BRD4-BD2 | BRD4-BD1 | BRD4-BD2 | HDAC1 | HDAC2 |
| MS436 | 4.0 ± 0.4 | 2.9 ± 0.1 | 2.85 ± 0.4 | 62 ± 15 | 18.8 ± 1.0 | > 50 |
| NB161 (**1**) | 5.2 ± 0.3 | 3.0 ± 0.2 | n.d. | n.d. | 10.8 ± 5.1 | > 50 |
| NB390 (**6**) | 7.5 ± 0.2 | 5.2 ± 0.2 | 0.19 ± 0.03 | 0.05 ± 0.01 | 2.1 ± 0.7 | 1.3 ± 0.9 |
| NB437 (**22**) | 4.0 ± 0.2 | 4.8 ± 0.2 | 0.18 ± 0.03 | 0.20 ± 0.03 | 23 ± 11 | 25 ± 12 |
| NB480 (**25**) | 1.9 ± 0.2 | 1.9 ± 0.3 | 2.55 ± 0.08 | 2.94 ± 0.18 | 2.3 ± 0.6 | 3.7 ± 2.4 |
| NB462 (**31a**) | 3.9 ± 0.4 | 4.5 ± 0.1 | 0.25 ± 0.02 | 0.19 ± 0.02 | 2.6 ± 1.1 | 1.6 ± 0.7 |
| (+)-JQ1 | 6.7 ± 0.1 | 5.8 ± 0.6 | 0.06 ± 0.01 | 0.11 ± 0.01 | n.d. | n.d. |
| CI-944 | n.d. | n.d. | n.d. | n.d. | 5.0 ± 0.8 | 2.0 ± 0.7 |

### Second-generation dual inhibitors: structure-guided optimization of the BET-binding moiety and linker variation

Next, we optimized the BET-binding moiety based on the crystal structure of BRD4 BD1 in complex with NB161. To improve binding, we first replaced the phenol moiety of NB161 with a hydroxyindole (**Scheme 1**), yielding compound NB390 (**6**). Pleasingly, NB390 showed a significantly higher thermal shift for BD1 and BD2 in DSF assays (Δ*T*_m_ = 7.5 K and 5.2 K, respectively) than NB161 (**Table 1**), which was comparable to the potency of (+)-JQ1, validating our design strategy. The crystal structure of the complex with BD1 showed the hydroxyindole moiety engaging in two hydrogen bonds with Asn140 in the acetyl-lysine binding pocket and forming additional hydrophobic interactions with Leu94 and the gatekeeper residue Ile146, as anticipated (**Figure 2B**). In the next optimization round, we modified the linker region to eliminate the chemically liable azo-moiety (**Figure 2A**). This strategy also allowed us to replace the 7-hydroxyindole moiety with a more stable pyrrolopyridone moiety, thereby facilitating compound synthesis while retaining all key interactions with the BD1/2 binding pocket.

**Figure 2.**
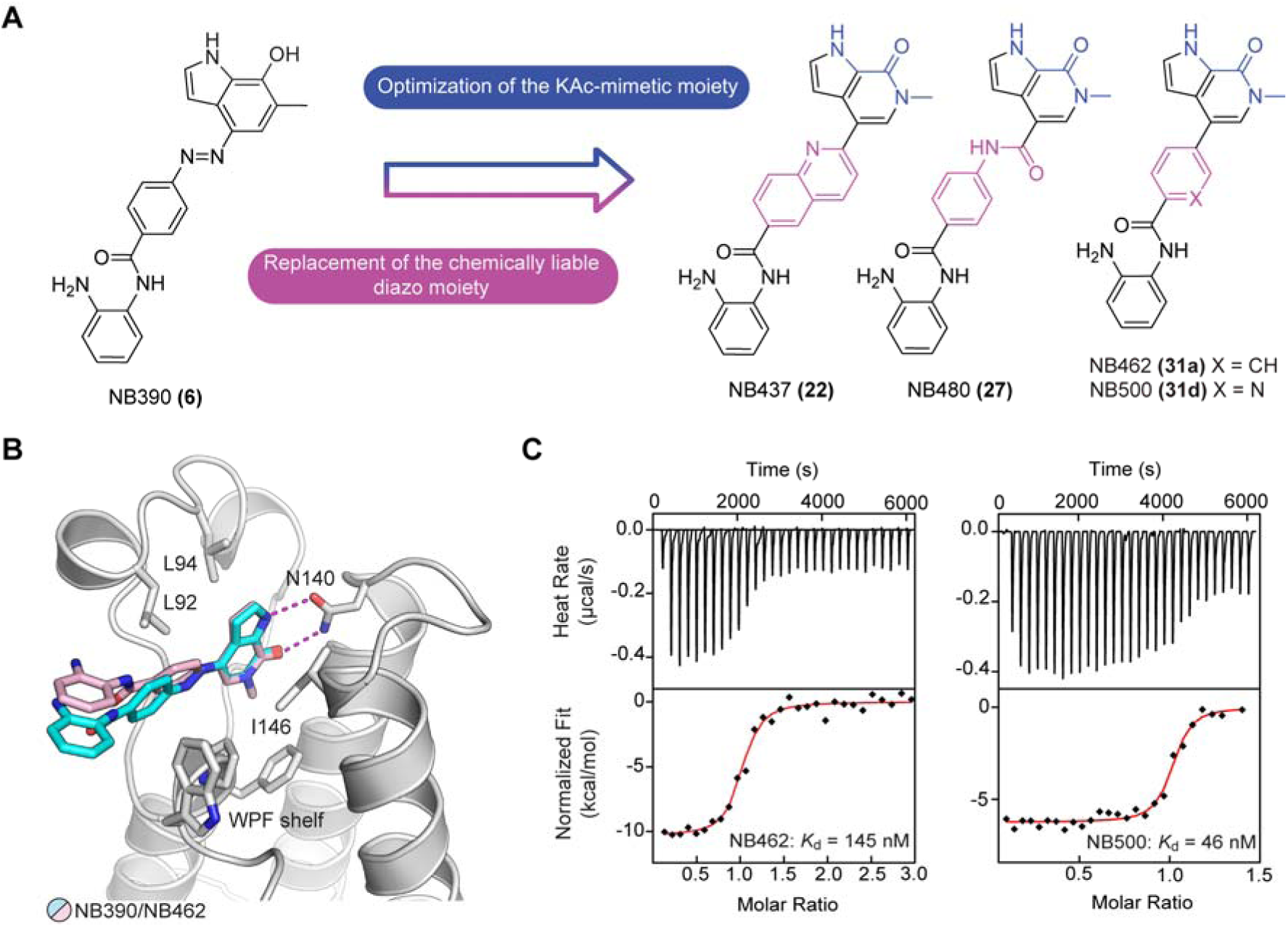
Development of second-generation dual inhibitors by structure-guided optimization of the acetyl lysine-mimetic moiety and the linker region. (A) SAR-guided optimization of the acetyl lysine-mimetic moiety and replacement of the diazo moiety. (B) Overlay of the crystal structures of BRD4 BD1 in complex with NB390 (cyan stick model) and NB462 (pink stick model) showing that the newly introduce phenyl linker in NB462 superimposes well with the position of the chemically liable diazo moiety in NB390. The protein is shown as a gray cartoon representation with selected interacting side chains highlighted as stick models. For clarity, only the BD1 domain of the NB462 complex is shown. (C) Representative ITC data of NB462 and NB500 binding to BRD4 BD1.

The synthesis of borylated intermediate **15** was adapted from published protocols [37, 38] (**Scheme 2**). For the reductive cyclization of **10** with iron powder, acetic acid was used as a cosolvent to improve solubility and reduce the amount of solvent needed for the reaction. The final borylation of **14** was carried out on a 10 g-scale, and the product could be triturated from hexane/diethyl ether, eliminating the need for chromatographic purification. The chlorinated quinoline intermediate **19** was synthesized *via* oxidation of quinoline **16** to the *N*-oxide **17** with *m*CPBA, followed by reaction with mesyl chloride/water and subsequent chlorination with thionyl chloride to yield intermediates **18** and **19 (Scheme 3A**). Suzuki-coupling of **19** and boronate **15** provided compound **20**, which was then hydrolyzed and coupled to *Boc*-protected *o*-phenylenediamine **2** to yield inhibitor NB437 (**22**), *via* intermediate **21**. Transmetallation of **14** with isopropyl magnesium chloride followed by reaction with dry ice provided carboxylic acid **23** (**Scheme 3B**). After basic detosylation, **24** was first converted into the acyl chloride and then reacted with methyl 4-aminobenzoate to yield **25**. Ester hydrolysis and amide coupling with *o*-phenylenediamine provided inhibitor NB480 (**27**). Suzuki coupling of the respective halobenzene **28a-f** with boronate **15** provided methyl esters **29a-f** (**Scheme 3C**). Saponification and amide coupling then readily provided inhibitors **31a-f**.

**Scheme 2.**
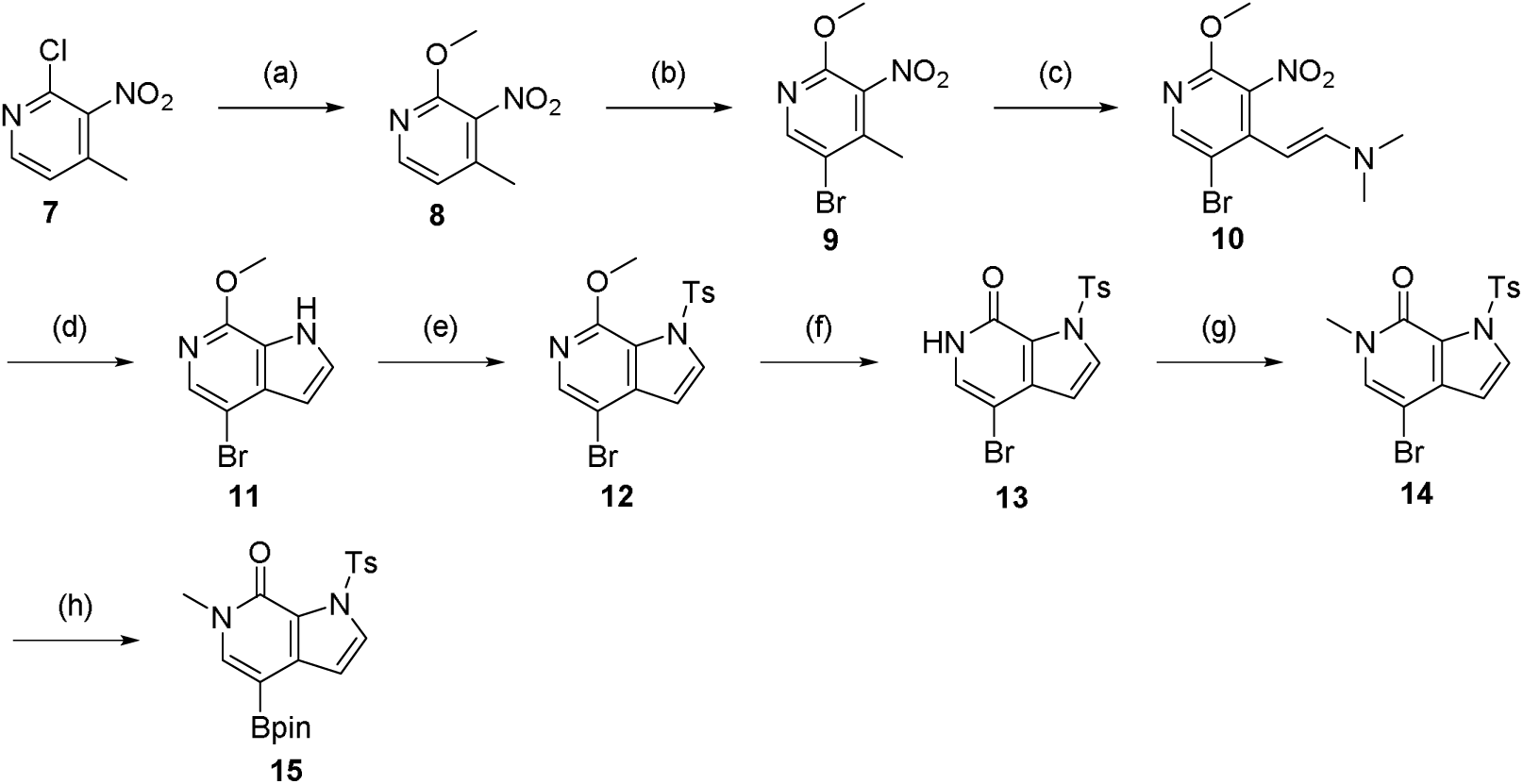
Synthesis of intermediate **15**. Reagents and conditions: (a) NaOMe, MeOH, reflux, 16 h; (b) Br_2_, NaOAc, AcOH, 80 °C, 16 h; (c) DMF-DMA, DMF, 90 °C, 16 h; (d) Fe, AcOH/MeOH/H_2_O, reflux, 2 h; (e) NaH, TsCl, THF, 0 °C to rt, 2 h; (f) HCl in dioxane, 50 °C, 2 h; (g) NaH, MeI, DMF, 0 °C to rt, 2 h; (h) B_2_pin_2_, KOAc, Pd XPhos G2, XPhos, dioxane, 80 °C, 2 h.

**Scheme 3.**
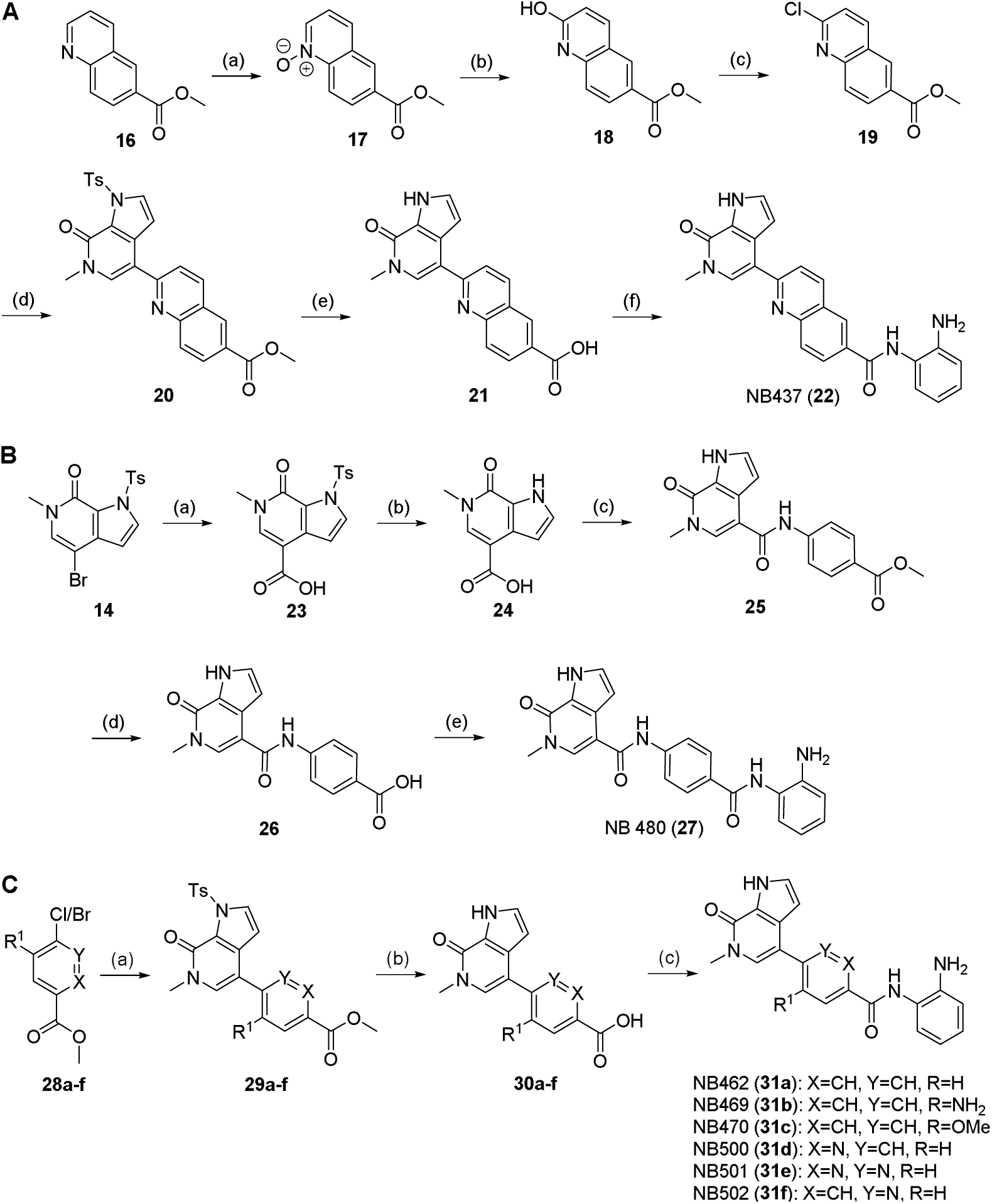
Synthesis of second-generation dual inhibitors. (A) Synthesis of inhibitor NB437. Reagents and conditions: (a) *m*CPBA, DCM, 0 C to rt, 3 h; (b) MsCl, H_2_O/ACN, rt, 45 min; (c) SOCl_2_, DMF, DCM, 0°C to rt, 16 h; (d) **15**, K_3_PO_4_, Pd XPhos G2, XPhos, dioxane/H_2_O, 70 °C, 1 h; (e) LiOH · H_2_O, dioxane/H_2_O, 80°C, 2 h; (f) (1) **2**, PyAOP, DIPEA, DMF, rt, 16 h; (2) TFA/DCM, rt, h. (B) Synthesis of inhibitor NB480. Reagents and conditions: (a) (1) *i*PrMgCl · LiCl, THF, −40 C, 2 h; (2) CO_2 (s)_, 0.5 h; (b) LiOH · H_2_O, dioxane/H_2_O, 90 C, 1 h; (c) (1) SOCl_2_, dioxane, 80 C, 16 h; (2) methyl 4-aminobenzoate, DIPEA, DMA, rt, 1 h; (d) LiOH · H_2_O, THF/MeOH/H_2_O, 60 °C, 1 h; (e) *o*-phenylene diamine, PyAOP, DIPEA, DMF, rt, 16 h. (C) Synthesis of inhibitors NB462, NB469, NB470, and NB500-502. Reagents and conditions: (a) **15**, K_3_PO_4_, Pd XPhos G2, XPhos, dioxane/H_2_O, 70 C, 1 h; (b) LiOH · H_2_O, dioxane/H_2_O, 80 C, 2 h; (c) *o*-phenylene diamine, PyAOP, DIPEA, DMF, rt, 16 h

For all three synthesized inhibitor variants, linker modification resulted in a reduced potency in DSF thermal shift assays (**Table 1**). Replacing the diazo group with an amide linker (NB480), for example, drastically reduced potency, and only a very modest Δ*T*_m_ of 1.9 K was observed for both BD1 and BD2. Substituting the phenyldiazene moiety with a quinoline (NB437) or simply deleting the diazo moiety (NB462, **31a**) was much better tolerated, though, resulting in two more or less equipotent BRD4 binders, with Δ*T*_m_ values between 4 and 5 K for BD1 and BD2, respectively. ITC measurements showed that NB462 binds to BD1 with a *K*_d_ of 145 nM (**Figure 2C**). High-resolution (1.2 Å) crystal structures of BD1 complexed with NB437 and NB462 revealed that the binding mode of the Asn140-targeting moiety was virtually unperturbed by the linker modification and that the loss in potency compared with the parent molecule for the shorter compound NB462 can be attributed to suboptimal hydrophobic packing against the WPF-shelf region (**Figure 2B** and **Supporting Information Figure S1**).

The cellular binding of inhibitors NB437 and NB462 was measured *via* a NanoBRET target engagement assay against the first and second bromodomain of BRD4 as well as against HDAC1/2 (**Table 1**). Both inhibitors bound to the BRD4 bromodomains with an EC_50_ in the range of 100-400 nM. When testing HDAC1/2 binding potency in cells by NanoBRET assay, NB462 was about an order of magnitude more potent than NB437 (EC_50_ of 2.6 vs 23 µM for HDAC1 and 1.6 vs 25 µM for HDAC2). We therefore decided to explore further SAR on the shorter, more soluble variant NB462, testing the effect of additional substituents or heteroatoms in the central aromatic ring of the linker (**Scheme 3C**). Replacing the phenyl group by a pyridine moiety in NB500 (**31d**) resulted in a three-fold increased binding affinity to BRD4 BD1 in ITC experiments, with a *K*_d_ value of 46 nM (**Figure 2C** and **Supporting Information Table S1**). The binding affinities of NB462 and NB500 to BD1 and BD2 in the cellular NanoBRET assays were virtually identical, though, and HDAC1/2 inhibition was reduced slightly (**Table 2**). Adding exocyclic substituents, however, drastically impaired both BET and HDAC inhibition in cells (**Table 2)**. Also, the position of the endocyclic nitrogen appears to be critical because nicotinamide-containing NB501 (**31e**) and pyridazine-containing NB502 (**31f**) showed an about two-fold reduced binding affinity to BRD4 compared with NB500.

**Table 2.** Modification of the central ring.

| Compounds | X | Y | R<sup>1</sup> | DSF $\Delta T_m$ (K) | | NanoBRET EC<sub>50</sub> (μM) | | | |
| --- | --- | --- | --- | --- | --- | --- | --- | --- | --- |
|  |  |  |  | BRD4-BD1 | BRD4-BD2 | BRD4-BD1 | BRD4-BD2 | HDAC1 | HDAC2 |
| NB462 (**31a**) | CH | CH | H | 3.9 ± 0.4 | 4.5 ± 0.1 | 0.25 ± 0.02 | 0.19 ± 0.02 | 2.6 ± 1.1 | 1.55 ± 0.73 |
| NB469 (**31b**) | CH | CH | NH<sub>2</sub> | 0.9 ± 0.2 | 1.7 ± 0.3 | 3.7 ± 0.5 | 2.05 ± 0.37 | 34.4 ± 5.8 | 39.7 ± 9.0 |
| NB470 (**31c**) | CH | CH | OMe | 3.8 ± 0.1 | 4.3 ± 0.3 | 0.39 ± 0.06 | 0.23 ± 0.07 | > 50 | > 50 |
| NB500 (**31d**) | N | CH | H | 4.2 ± 0.4 | 4.7 ± 0.4 | 0.24 ± 0.02 | 0.16 ± 0.04 | 4.8 ± 2.3 | 2.48 ± 0.66 |
| NB501 (**31e**) | CH | N | H | 2.9 ± 0.3 | 3.6 ± 0.2 | 0.39 ± 0.02 | 0.29 ± 0.12 | 16.8 ± 6.7 | 4.56 ± 0.57 |
| NB502 (**31f**) | N | N | H | 2.6 ± 0.0 | 3.4 ± 0.2 | 0.48 ± 0.14 | 0.32 ± 0.13 | 6.1 ± 1.0 | 3.09 ± 0.13 |
| (+)-JQ1 | - | - | - | 6.7 ± 0.1 | 5.8 ± 0.6 | 0.06 ± 0.01 | 0.11 ± 0.01 | n.d. | n.d. |
| CI-944 | - | - | - | n.d. | n.d. | n.d. | n.d. | 5.0 ± 0.8 | 2.0 ± 0.7 |

### Optimization of the HDAC-binding moiety

For final optimization of our dual inhibitors, we aimed to improve HDAC1/2 binding by targeting the hydrophobic, so-called foot pocket next to the catalytic zinc ion [39] (**Figure 3A**). Appropriately substituted anilines **35a-c** were accessible through Suzuki coupling of *Boc*-protected 4-bromo-2-nitroaniline **33** with different aryl boronic acids and subsequent reduction with iron powder (**Scheme 4A**). Amide coupling with carboxylic acids **30a** and **d**, followed by *Boc*-deprotection then provided the derivatized inhibitors NB503 (**38**) and NB512-514 (**39a-c**) (**Scheme 4B**). Amide coupling of carboxylic acid **30d** with 4-fluorobenzene-1,2-diamine yielded inhibitor NB507 (**40**) (**Scheme 4C**). By introducing a phenyl or 2-thienyl moiety at position 5, the affinity for HDAC1 and HDAC2 could be significantly improved, resulting in nanomolar EC_50_ values (**Table 3**, **Figure 3D**), with phenyl-substituted NB512 showing the strongest binding to HDAC1 (EC_50_ = 110 nM) and thienyl-substituted NB514 the strongest binding to HDAC2 (EC_50_ = 60 nM). The foot pocket in human HDAC3 is smaller than in HDAC1/2 due to variation of one of the residues lining the pocket (Tyr107 in HDAC3, whereas HDAC1/2 have a serine at the equivalent position). Accordingly, targeting the foot pocket in HDAC3 had only a relatively small effect on inhibitor binding, with HDAC3 EC_50_ values of the third-generation inhibitors remaining in the low- to mid-micromolar range (**Table 3, Supporting Information Figure S2**).

**Figure 3.**
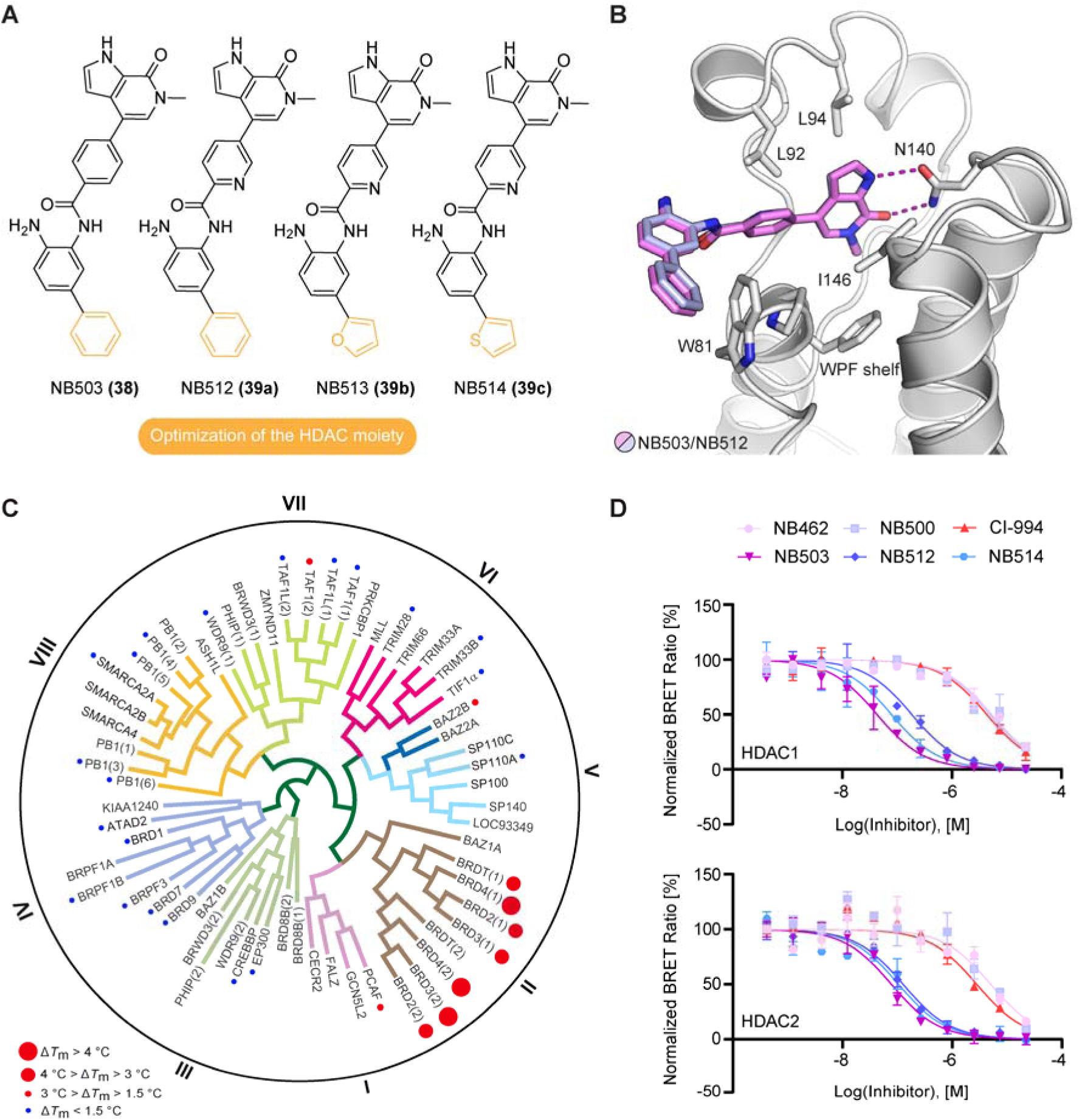
Development of third-generation dual inhibitors with optimized warhead for HDAC1/2 inhibition. (A) Chemical structures of dual inhibitors with modified HDAC binding moiety. (B) Superposition of the crystals structures of NB503 (magenta stick model) and NB512 (pale blue stick model) bound to BRD4 BD1 (gray ribbon diagram). For clarity, only the protein domain for the NB503 complex is shown. Selected side chains are highlighted as gray stick models. In both complexes, the newly introduced HDAC moiety of the inhibitor protrudes into the solvent next to the WPF shelf, weakly interacting with Trp81. (C) DSF bromodomain selectivity panel for inhibitor NB503 measured at a compound concentration of 10 µM showing high selectivity for the BET family domains. (D) Representative NanoBRET data and fits for inhibitor binding to HDAC1 and HDAC2 measured in intact cells.

**Table 3.**
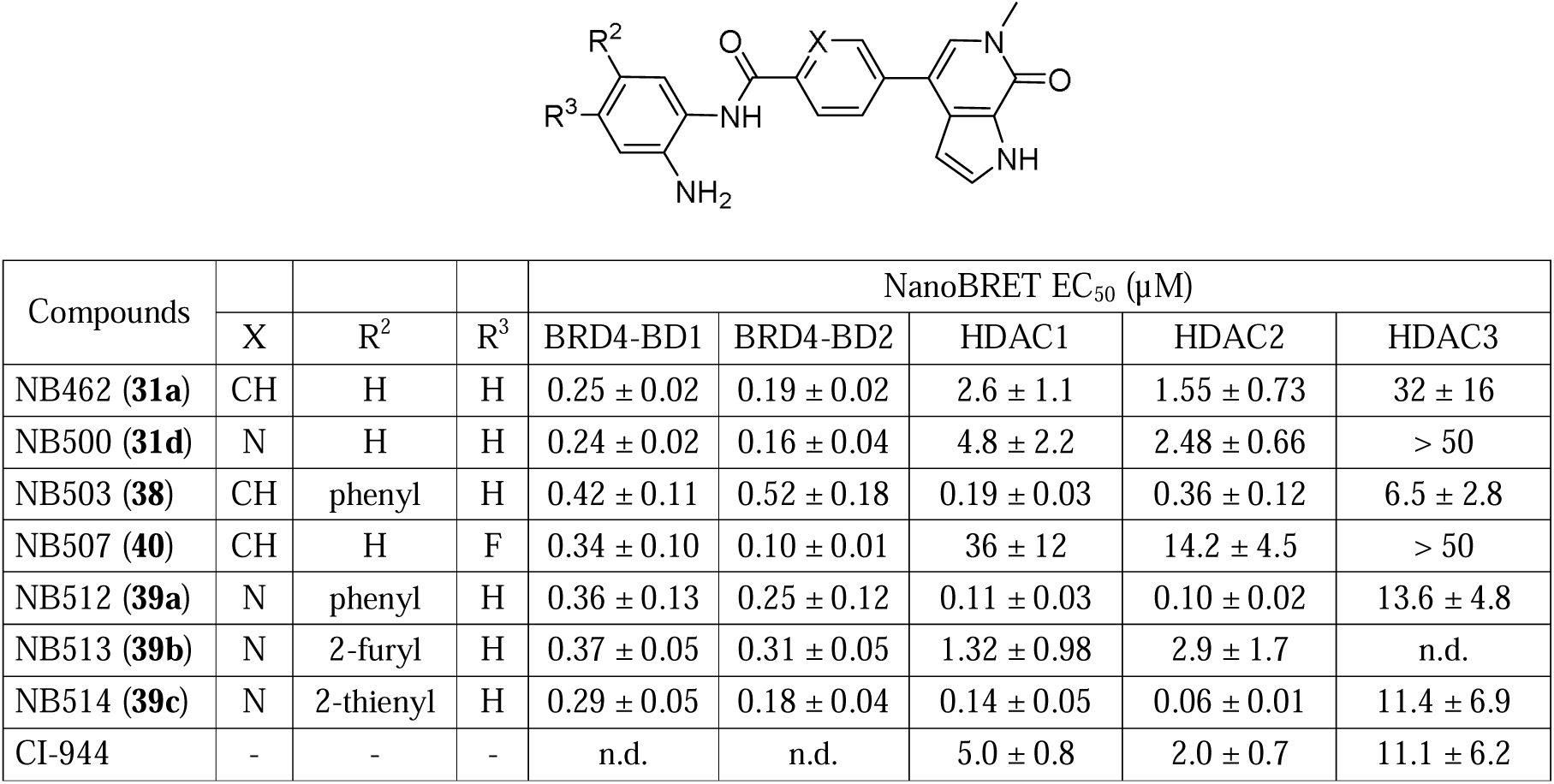
Optimization of the HDAC warhead.

**Scheme 4.**
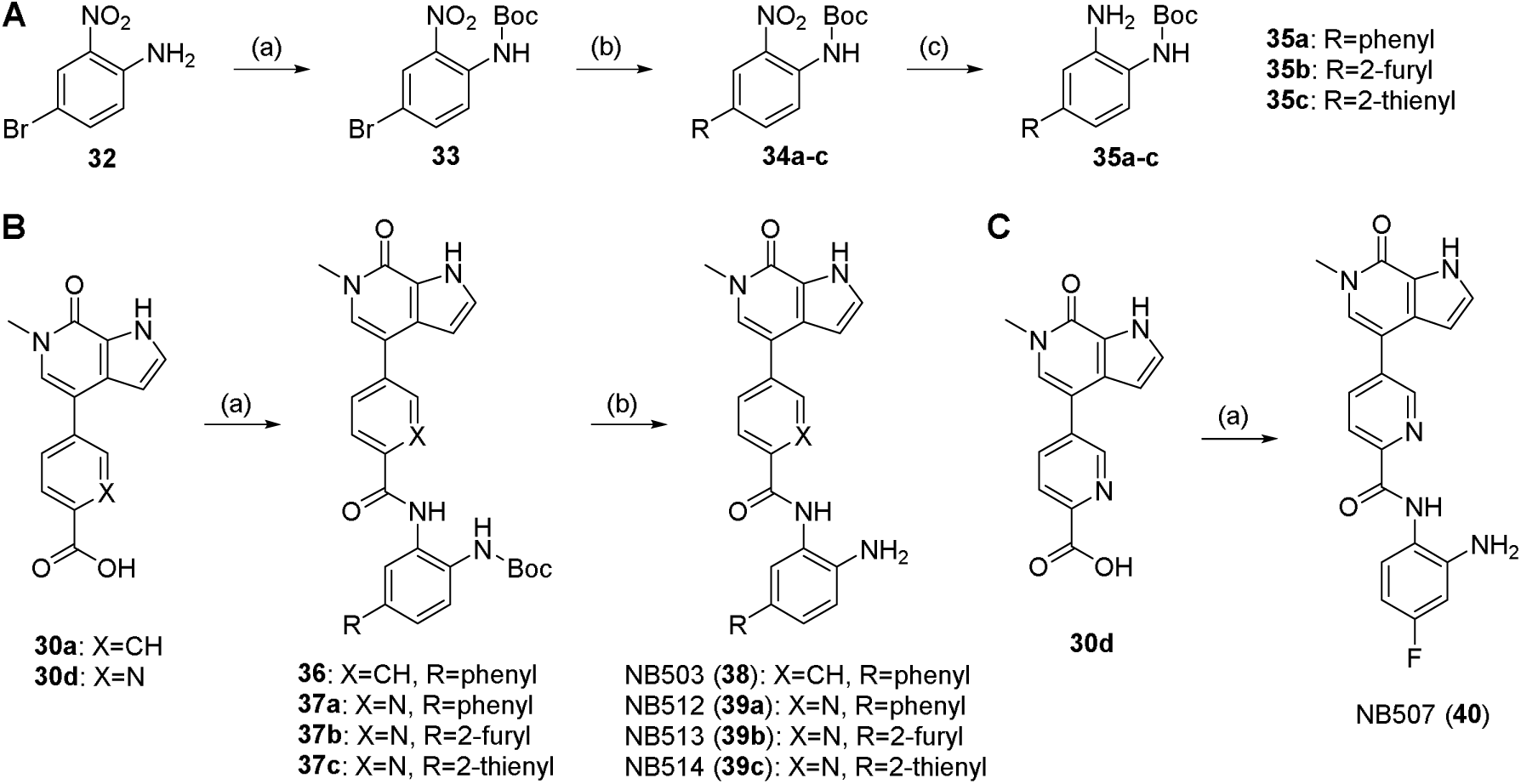
Synthesis of third-generation dual inhibitors (A) Synthesis of *Boc*-protected *o*-phenylenediamines. Reagents and conditions: (a) NaH, Boc_2_O, THF, −10 C to rt, 4 h; (b) arylboronic acid, K_2_CO_3_, Pd XPhos G2, XPhos, DMF/H_2_O, 100°C, 2 h; (c) Fe, NH_4_Cl, MeOH/H_2_O, 85°C, 3 h. (B) Synthesis of inhibitors NB503, NB512, NB513, and NB514. Reagents and conditions: (a) **35a-c**, PyAOP, DIPEA, DMF, rt, 16 h; (b) TFA/DCM, rt, 1 h. (C) Synthesis of inhibitor NB507. Reagents and conditions: (a) 4-fluorobenzene-1,2-diamine, PyAOP, DIPEA, DMF, rt, 16 h.

As expected based on the literature [40], 4-fluorination led to a decreased HDAC1/2 binding affinity in the case of NB507. The modifications on the HDAC moiety in these third-generation inhibitors had only a minimal effect on BRD4 EC_50_ values, consistent with the crystal structures of the BRD4 BD1-NB503/NB512 complexes showing that the phenyl group of the HDAC warhead protrudes into the solvent, albeit also weakly interacting with Trp81 of the WPF shelf region (**Figure 3B**).

A DSF selectivity screen of NB462 against a panel of 32 bromodomains covering all main branches of the bromodomain tree confirmed high stabilization of BET bromodomains but also revealed BRD7 and BRD9 as prominent off-targets (**Supporting Information Table S2**). The BRD7/9 activity of NB462 was not unexpected, given the similarity of the basic scaffold with that of the published BRD9 chemical probes BI-7273 and BI-9564 [41]. Intriguingly, modification of the HDAC warhead, however, drastically improved the selectivity of the dual inhibitors, and the DSF screen of NB503 against our in-house bromodomain panel revealed high selectivity for BET bromodomains (**Figure 3C** and **Supporting Information Table S2**) and elimination of the BRD7/9 off-target activity seen for the precursor molecule NB462. This gain in selectivity suggests that the aniline moiety of NB462 facilitates stabilizing interactions with BRD7/9 that are perturbed upon addition of an aromatic substituent at the 5-position in NB503.

### Biological evaluation of the dual BET/HDAC inhibitors in cancer cell lines

The biological effects of our optimized dual inhibitors were tested in pancreatic cancer cell line PaTu8988T. The most potent second- and third-generation inhibitors effectively reduced cell viability, with IC_50_ values in the low micromolar range (**Figure 4E** and **Supporting Information Table S3**). HDAC activity was evaluated by monitoring histone H3 acetylation 48h after incubation with 1 µM inhibitor by Western blot (**Figure 4A**). The highest histone H3 K9/K14 acetylation levels, indicative of HDAC inhibition, were observed with the third-generation inhibitors with an aromatic substituent in the HDAC warhead. This effect was concentration-dependent, as shown for NB503 and NB512, with NB512 being the more potent inhibitor overall (**Figure 4B**), consistent with its 2-3 times lower EC_50_ values for HDAC1 and HDAC2 in the NanoBRET assays. Surprisingly, one of the inhibitors without optimized HDAC moiety, NB462, was significantly more potent than CI-944 and almost as potent as NB503 and NB512 in inhibiting histone H3 deacetylation. This is particularly interesting, given that NB500, which differs only in having a nitrogen heteroatom in the aromatic linker between the BET and HDAC warhead, was much less potent in preventing histone H3 deacetylation, despite only moderately increased EC_50_ values for HDAC1/2 inhibition in NanoBRET assays.

**Figure 4.**
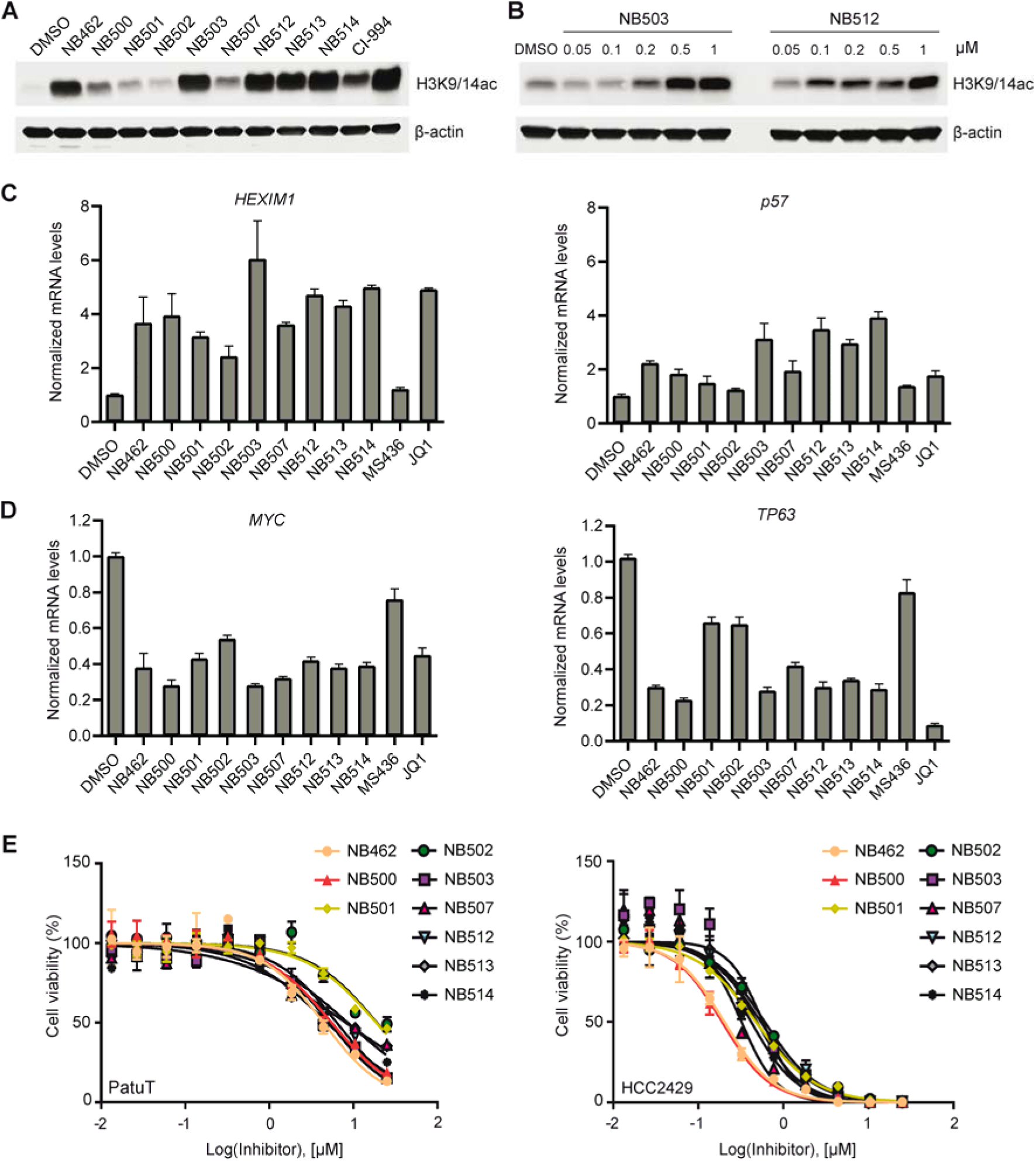
Biological effects of the optimized dual BET/HDAC inhibitors. (A) Effect on histone H3 K9/K14 acetylation in Patu8988T cells 48 h after incubation with 1 μΜ compound monitored by Western blot. (B) Western blot showing the concentration-dependent inhibition of histone H3 K9/K14 deacetylation in Patu8988T cells 48 h after treatment with NB503 and NB512. (C) Upregulation of mRNA levels of BET-inhibition biomarkers *HEXIM1* and *p57* in Patu8988T cells 6 h after treatment with 1 μΜ compound. (D) mRNA levels of oncogenic drivers *MYC* and *TP63* in NMC cells 6 h after treatment with 1 μΜ compound, showing that the optimized dual inhibitors significantly downregulated both transcription factors. (E) Cell viability of pancreatic cancer cell line PatuT (*left*) and NMC cell line HCC2429 (*right*) after 3d-treatment with different concentrations of dual BET/HDAC inhibitors.

To assess the effect of BET inhibition, we analyzed the mRNA levels of a well-characterized BET-targeting marker, *HEXIM1*, by quantitative RT-PCR. After treating PaTu 8988T cells for 6 h with 1 μΜ compound, all tested inhibitors resulted in increased mRNA levels of *HEXIM1*, with NB503 exhibiting the strongest effect (**Figure 4C**). The cell-cycle regulator gene *p57* was previously reported to be induced by combined BET/HDAC inhibition [8]. Consistently, *p57* RNA levels were most increased by the third-generation dual inhibitors (**Figure 4C**), significantly more than with MS463, which was expected given its lower BET inhibition potency, but also more than with (+)-JQ1.

We next analyzed the gene expression levels of the transcription factors *MYC* and *TP63* in NMC HCC2429 cells, which are sensitive to BET inhibition. NMC is driven by the BRD4-NUT fusion oncoprotein, which blocks differentiation and drives growth of NMC cells through the formation of hyperacetylated megadomains, long contiguous stretches of active chromatin resulting, for example, in upregulation of *MYC* and *TP63*, whose ΔN isoform is oncogenic [42, 43]. Disturbing these BRD4-NUT megadomains by BET inhibition should therefore downregulate *MYC* and *TP63* expression. And indeed, all dual inhibitors tested caused a significant decrease in *MYC* and *TP63* mRNA levels compared with untreated controls, with NB500 and NB503 showing the strongest effect (**Figure 4D**). All second and third-generation dual inhibitors were more potent than the parent molecule, MS436, consistent with the NanoBRET BRD4 binding data. Compared with (+)-JQ1, however, the scenario was different: all third-generation dual inhibitors were more potent in downregulating *MYC*, despite lower on-target BRD4 affinity in NanoBRET assays, but significantly less potent in downregulating *TP63* than (+)-JQ1. Interestingly, the inhibitors were about one order of magnitude more potent in reducing the viability of NMC cells than that of PaTu8988T cells, with IC_50_ values ranging from 200-600 nM (**Figure 4E** and **Supporting Information Table S3**).

## Conclusion

Through pharmacophore merging followed by structure-guided design, we have successfully developed a new series of potent and selective dual BET/HDAC inhibitors. The choice of a suitable starting scaffold enabled us to optimally merge and integrate the two functionalities into a single, compact molecule rather than simply linking two existing inhibitors. The best molecules in our lead series bound to both BRD4 bromodomains as well as HDAC1/2 with affinities in the 100-nanomolar range in NanoBRET cellular target engagement assays. Gratifyingly, this on-target potency was also reflected in promising biological effects in pancreatic cancer cells where NB503 and NB512 effectively blocked histone H3 deacetylation and upregulated specific markers for BET inhibition (*HEXIM1* and *p57*). In addition, we could show that our molecules induce the downregulation of oncogenic drivers of NMC, as demonstrated for *MYC* and *TP63*. Overall, this work expands the set of available dual BET/HDAC inhibitors for cancer therapy and provides a basis for future translational studies in different cancer types.

## MATERIALS AND METHODS

### Chemistry

Compound synthesis is described in detail in the Supporting Information, including analytical data for all final products. All commercial chemicals were purchased from common suppliers in reagent grade and used without further purification. For compound purification by flash chromatography, a puriFlash XS 420 device with a UV-VIS multiwave detector (200–400 nm) from Interchim with pre-packed normal-phase PF-SIHP silica columns with particle sizes of 30 μm (Interchim) was used. Synthesized compounds were characterized by NMR and mass spectrometry (ESI). In addition, final inhibitors were identified by high-resolution mass spectrometry (HRMS), and their purity was evaluated by HPLC. ^1^H and ^13^C NMR spectra were measured on an AV300, an AV400, or an AV500 HD AVANCE III spectrometer from Bruker. Chemical shifts (δ) are reported in parts per million (ppm). DMSO-*d*_6_ was used as a solvent, and the spectra were referenced to the residual solvent signal: 2.50 ppm (^1^H NMR) or 39.52 ppm (^13^C NMR). HRMS was measured on a MALDI LTQ Orbitrap XL from ThermoScientific. Determination of the compound purity by HPLC was carried out on an Agilent 1260 Infinity II device with a 1260 DAD HS detector (G7117C; 254 nm, 280 nm, 310 nm) and an LC/MSD device (G6125B, ESI pos. 100-1000). The compounds were analyzed on a Poroshell 120 EC-C18 (Agilent, 3 x 150 mm, 2.7 µm) reversed phase column using 0.1% formic acid in water (A) and 0.1% formic acid in acetonitrile (B) as a mobile phase. The following gradient was used: 0 min: 5% B - 2 min: 80% B - 5 min: 95% B - 7 min: 95% B (flow rate of 0.6 mL/min.). UV-detection was performed at 320 nm (150 nm bandwidth), and all compounds used for further biological characterization showed > 95% purity.

### Protein expression and purification

Bromodomain-containing proteins used in the selectivity panel were expressed and purified as previously described [44]. BRD4 BD1 (residues N44-E168) for structural studies was subcloned in pNIC28-Bsa4 vector (N-terminal His_6_-tag, followed by a TEV cleavage site). The expression plasmids were transformed into *Escherichia coli* BL21(D3)-R3-pRARE2 Rosetta cells. Cells were cultured in Terrific Broth (TB) media at 37 °C to an optical density (OD) of 2.8-3.0, and then expression was induced with 0.5 mM IPTG at 18 °C overnight. Cells were harvested and resuspended in a buffer containing 50 mM HEPES, pH 7.5, 500 mM NaCl, 0.5 mM TCEP, 5% glycerol and subsequently lysed by sonication. The recombinant protein was initially purified by Ni^2+^-affinity chromatography. The histidine tag was then removed by TEV protease treatment overnight, and the cleaved protein was separated by reverse Ni^2+^-affinity purification. The protein was further purified by size exclusion chromatography using a HiLoad 16/600 Superdex 75 column with buffer containing 25 mM HEPES, 150 mM NaCl, 0.5 mM TCEP, and 5% glycerol. Quality control was performed by SDS-polyacrylamide gel electrophoresis and ESI-MS (BRD4 BD1: expected mass 15,083.5 Da, observed mass 15,084.1 Da).

### Crystallization and structure determination

Crystals of BRD4 BD1 in complex with dual inhibitors were grown using the sitting-drop vapor-diffusion technique at 277 K utilizing a mosquito crystallization robot (TTP Labtech, Royston, UK). BRD4-BD1 protein (10 mg/mL in 25 mM HEPES pH 7.5, 150 mM NaCl, 0.5 mM TCEP, 5% glycerol) was incubated with inhibitors at a final concentration of 1 mM prior to setting up crystallization trials. Detailed crystallization conditions for each inhibitor are listed in **Supporting Information Table S4**. Crystals were cryo-protected with mother liquor supplemented with 23% ethylene glycol and flash-frozen in liquid nitrogen. X-ray diffraction data sets were collected at 100 K at beamline X06SA of the Swiss Light Source, Villigen, Switzerland. The obtained diffraction data were integrated with the program XDS [45] and scaled with AIMLESS [46], which is part of the CCP4 package [47]. The structures were then solved by molecular replacement using PHASER [48] or by difference Fourier analysis using PHENIX [49] with PDB entry 6YQN as a starting model. Structure refinement was performed using iterative cycles of manual model building in COOT [50] and refinement in PHENIX. Dictionary files for the compounds were generated using the Grade Web Server (http://grade.globalphasing.org). X-ray data collection and refinement statistics are listed in **Supporting Information Table S5**.

### Differential scanning fluorimetry (DSF)

The effects of inhibitor binding on the apparent melting temperature of recombinant bromodomains were determined by DSF in a 96-well plate (Starlab) at a protein concentration of 2 μM with 10 μΜ compound in buffer containing 25 mM HEPES, pH 7.5, 150 mM NaCl, and 0.5 mM TCEP. SYPRO Orange (5000×, Invitrogen), a dye that shows strong fluorescence upon binding to hydrophobic regions of unfolded proteins, was added at a dilution of 1:1000 (final concentration of 5x). Protein unfolding profiles were recorded using an MX3005P real-time qPCR instrument (Agilent; excitation/emission filters = 492/610 nm) while increasing the temperature from 25 to 95 °C at a heating rate of 3 °C/min. *T*_m_ values were calculated after fitting the fluorescence curves to the Boltzmann equation. Differences in melting temperature upon compound binding are given as Δ*T*_m_ = *T*_m_ (protein with inhibitor) - *T*_m_ (protein without inhibitor). Measurements were performed in triplicates. All bromodomain selectivity-panel data are listed in **Supporting Information Table S2**.

### Isothermal titration calorimetry (ITC)

ITC measurements were performed using a Nano ITC micro-calorimeter (TA Instruments, New Castle, Pennsylvania). For all experiments, reverse titration was performed (syringe containing the protein solution; cell containing the ligand) in ITC buffer containing 25 mM HEPES, pH 7.5, 150 mM NaCl, 0.5 mM TCEP, and 5% glycerol. All compounds were diluted from 50 mM DMSO stocks to 20 μΜ in ITC buffer and BRD4-BD1 was diluted to 120 μΜ in a DMSO-adjusted ITC buffer. BRD4-BD1 (120 μΜ) was titrated into the compound solution (20 μΜ) with an initial injection (4 μL) followed by 29 identical injections (8 μL), at a rate of 0.5 μL/s and with 150 or 200 s intervals. All experiments were performed at 15 °C whilst stirring at 350 rpm. The heat of dilution was determined by independent titrations (protein into buffer) and was subtracted from the experimental raw data. Data were processed using the NanoAnalyze software (version 3.10.0) provided by the instrument manufacturer. The first injection was excluded from the analysis, and fitted curves were generated by applying the independent model (single binding site) to the raw data. Complete thermodynamic data analysis can be found in **Supporting Information Table S1**.

### NanoBRET assay

The assay was performed as described previously [12, 51]. In brief: BRD4 bromodomains and full-length HDACs were obtained as plasmids cloned in frame with a terminal NanoLuc fusion (Promega). Plasmids were transfected into HEK293T cells using FuGENE HD (Promega, E2312), and proteins were allowed to express for 20 h. Serially diluted inhibitor and the corresponding NanoBRET Tracer (Promega) at a concentration determined previously as the tracer *K*_D,app_ were pipetted into white 384-well plates (Greiner #781207) for BRD assays or white 96-well plates (Corning #3600) for HDAC using an Echo acoustic dispenser (Labcyte). The corresponding protein-transfected cells were added and reseeded at a density of 2x 10^5^ cells/mL after trypsinization and resuspending in Opti-MEM without phenol red (Life Technologies). The system was allowed to equilibrate for 2 hours at 37 °C/5% CO_2_ prior to BRET measurements. To measure BRET, NanoBRET NanoGlo substrate + Extracellular NanoLuc Inhibitor (Promega, N2540) was added as per the manufacturer’s protocol, and filtered luminescence was measured on a PHERAstar plate reader (BMG Labtech) equipped with a luminescence filter pair (450 nm BP filter (donor) and 610 nm LP filter (acceptor)). Competitive displacement data were then graphed using a normalized 3-parameter curve fit with the following equation: Y=100/(1+10^(X-LogIC50)). Resulting EC_50_s are an average and standard error of the mean of at least three independent experiments that were themselves performed in technical duplicates (n=2).

### Cell culture and reagents

Pancreatic ductal adenocarcinoma cell line PaTu 8988t was obtained from ATCC and cultured in Dulbecco’s Modified Eagle Medium (DMEM) containing 10% FBS, 25 mM glucose, 4 L-glutamine, 1 mM sodium pyruvate and 1% penicillin-streptomycin. Nut midline carcinoma cell line HCC2429 was kindly provided by Lead Discovery Center GmbH (Dortmund, Germany) and was cultured in RPMI1640 medium containing 10% FBS, 2 mM L-glutamine and 1% penicillin-streptomycin. Cell line PaTu 8988t was authenticated using Multiplex human Cell line Authentication Test (MCA) by Multiplexion GmbH and cell line HCC2429 was authenticated using short tandem repeat (STR) profiling.

### Immunoblot analysis

Protein samples were prepared in RIPA buffer (9806S, Cell Signaling Technology) containing protease inhibitor cocktail (Roche). Proteins were separated in SDS-polyacrylamide gels, transferred to nitrocellulose membranes with Trans-Blot Turbo Transfer System (Bio-Rad) and incubated with antibodies dissolved in TBS buffer containing 5% BSA and 0.1% Tween20. The following primary antibodies were used: rabbit anti-β-actin (ab8227, Abcam) and rabbit anti-acetyl-histone H3 (Lys9/Lys14; 9677, Cell Signaling Technology). Primary antibodies were recognized by a peroxidase-coupled secondary antibody (Jackson), and signals were detected by chemiluminescence.

### RNA extraction and quantitative RT-PCR analysis

Total RNA was extracted from cell culture using Maxwell RSC simplyRNA Cells Kit (Promega) according to the manufacturer’s protocol. cDNA was synthesized using PrimeScript Reverse Transcriptase (TakaRa) and amplified using home-made PCR master mix. The amplicon was detected by EvaGreen Dye using LightCycler 480 instrument (Roche). PCR conditions were 5 min at 95 °C, followed by 45 cycles of 95 °C for 10 sec, 59 °C for 10 sec and 72 °C for 20 sec. The relative gene expression levels were normalized to GUSB and calculated using 2^−ΔΔCt^ method. The PCR primers used are listed in **Supporting Information Table S6**.

### Statistical analyses and figures

Structural images were generated using PyMOL (*Schrödinger LCC*), and graphs were plotted using GraphPad Prism version 8.4, (*GraphPad Software*, San Diego, California, USA, www.graphpad.com).

## Supporting Information

Additional experimental details, materials, and methods, DSF bromodomain selectivity panel, cell-viability data, X-ray data collection and refinement statistics, HPLC and NMR spectra for all compounds (PDF)”.

## Acknowledgments

The authors are grateful for support by the Structural Genomics Consortium (SGC), a registered charity (No:1097737) a registered charity (no: 1097737) that receives funds from Bayer AG, Boehringer Ingelheim, Bristol Myers Squibb, Genentech, Genome Canada through Ontario Genomics Institute [OGI-196], EU/EFPIA/OICR/McGill/KTH/Diamond Innovative Medicines Initiative 2 Joint Undertaking [EUbOPEN grant 875510], Janssen, Pfizer and Takeda. S.K. is funded by the German Cancer Research Center (DKTK) and the Frankfurt Cancer Institute (FCI). S.K. and B-T.B. also received support from the Collaborative Research Center CRC1399 “Mechanisms of drug sensitivity and resistance in small cell lung cancer”. A.C.J. and D.-I.B. are funded by German Research Foundation (DFG) grant JO 1473/1-3 (A.C.J.). J.T.S. is grateful for support by the German Cancer Consortium (DKTK), by the Deutsche Forschungsgemeinschaft (DFG, German Research Foundation; #405344257/SI 1549/3-2 and SI1549/4-1), by the German Cancer Aid (#70112505/PIPAC, #70113834/PREDICT-PACA) and by the German Federal Ministry of Education and Research (BMBF; 01KD2206A/SATURN3).

## Data availability

The atomic coordinates and structure factors of BRD4 BD1 in complex with the dual inhibitors have been deposited in the Protein Data Bank (PDB) under accession codes 8P9F (NB161), 8P9G (NB390), 8P9H (NB437), 8P9I (NB462), 8P9J (NB500), 8P9K (NB503), and 8P9L (NB512).

