## Supporting information for "Development of potent dual BET/HDAC inhibitors *via* pharmacophore merging and structure-guided optimization"

### Chemical Synthesis

#### General procedure for azo coupling

The aniline (1.0 eq) was dissolved in MeOH/ACN (1/1) and cooled to -10 °C. To this solution were added conc. HCl (1.0 eq) and *iso*-amyl nitrite (1.0 eq), and the mixture was stirred for 1 h, during which the mixture turned yellow due to the formation of the diazonium salt. In a separate flask, the phenol (1.0 eq) and $K_2CO_3$ (5.0 eq) were dissolved in MeOH/$H_2O$/ACN (3/2/1), the mixture was purged with argon for 10 min and cooled to -10 °C. The dissolved diazonium compound was slowly added to the second mixture, which instantly turned red. The mixture was stirred for 2 h while being slowly warmed to ambient temperature. Then, the mixture was partitioned between water and ethyl acetate, and the organic layer was washed with brine, dried over $MgSO_4$, filtered and concentrated under reduced pressure. The crude product was purified by flash chromatography to provide the title compound.

#### General procedure for *N-Boc* deprotection

The *Boc*-protected aniline was dissolved in DCM/TFA (3/1) and stirred for 1 h at ambient temperature. Afterwards, volatiles were removed under reduced pressure, the residue was dissolved in ethyl acetate and washed with sat. aq $NaHCO_3$ and brine. The organic phase was dried over $MgSO_4$, filtered, and volatiles were removed under reduced pressure.

#### General procedure for Suzuki coupling A

The respective arylboronic acid (1.2 eq), *tert*-Butyl (4-bromo-2-nitrophenyl)carbamate (**33**, 1.0 eq), potassium carbonate (3.0 eq), Pd XPhos G2 (0.05 eq) and XPhos (0.05 eq) were dissolved in DMF/water (2/1, 85 mM) and purged with argon for 10 min. The mixture was heated at 100 °C for 2 h, cooled to ambient temperature and partitioned between water and DCM. The DCM layer was washed with brine, dried over $MgSO_4$, filtered, and volatiles were removed under reduced pressure to provide the crude product. The residue was then triturated with methanol and filtered.

#### General procedure for Suzuki coupling B

6-Methyl-4-(4,4,5,5-tetramethyl-1,3,2-dioxaborolan-2-yl)-1-tosyl-1,6-dihydro-7*H*-pyrrolo[2,3-*c*]pyridin-7-one (**15**, 1.0 eq), the aryl halide (1.2 eq), $K_3PO_4$ (2.5 eq), Pd XPhos G2 (0.05 eq) and XPhos (0.05 eq) were dissolved in dioxane/water (4/1, 50 mM) and purged with argon for 10 minutes. The mixture was heated at 70 °C for 1 h, cooled to ambient temperature and partitioned between water and ethyl acetate. The ethyl acetate layer was washed with brine, dried over $MgSO_4$, filtered, and volatiles were removed under reduced pressure to provide the crude product.

#### General procedure for reduction of nitro groups

A mixture of the starting material (1.0 eq), ammonium chloride (7.0 eq) and iron powder (7.0 eq) in methanol/water (9/1) was heated at 85 °C for 3 h. After cooling to ambient temperature, the mixture was filtered through a plug of celite, and the filtrate was partitioned between water and DCM. The DCM layer was washed with brine, dried over $MgSO_4$, filtered, and volatiles were removed under reduced pressure.

##### General procedure for ester and tosylamide hydrolysis

The ester and LiOH · $H_2O$ (7 eq) were dissolved in dioxane/water (4/1, 100 mM) and heated at 80 °C for 2 h. The mixture was cooled to ambient temperature and brought to a pH of 3 by the addition of 5% aq HCl. The resulting precipitate was filtered and dried under reduced pressure to provide the product.

##### General procedure for amide coupling

Carboxylic acid (1.0 eq), amine (1.2 eq) and (7-Azabenzotriazol-1-yloxy)tripyrrolidinophosphonium hexafluorophosphate (PyAOP, 1.3 eq) were dissolved in anh. DMF (40 mM) and *N*,*N*-diisopropylethylamine (1.3 eq) was added. The mixture was stirred at ambient temperature for 16 h and partitioned between water and ethyl acetate. The ethyl acetate layer was washed with brine, dried over $MgSO_4$, filtered, and volatiles were removed under reduced pressure.

***tert*-butyl (2-aminophenyl)carbamate (2)**

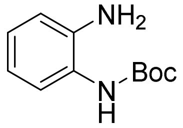


Benzene-1,2-diamine (5.00 g, 46.2 mmol, 1.0 eq) was dissolved in DCM (200 mL) and cooled to 0 °C. A solution of di-*tert*-butyl dicarbonate (10.09 g, 46.2 mmol, 1.0 eq) in DCM (100 mL) was added dropwise and the mixture was stirred for 16 h. Afterwards, all volatiles were removed under reduced pressure and the residue was crystallized from hexane/ethyl acetate to provide the title compound as colorless crystals (7.89 g, 82%). $^1$H NMR (400 MHz, DMSO-$d_6$) δ 8.26 (s, 1H), 7.19 (d, $J$ = 7.9 Hz, 1H), 6.84 (td, $J$ = 7.6, 1.5 Hz, 1H), 6.69 (dd, $J$ = 8.0, 1.5 Hz, 1H), 6.53 (td, $J$ = 7.5, 1.5 Hz, 1H), 4.80 (s, 2H), 1.47 (d, $J$ = 4.5 Hz, 9H). MS (ESI): m/z calc. for $[C_{11}H_{16}N_2O_2 + Na^+]^+$ = 231.11, found = 231.05

**(9*H*-Fluoren-9-yl)methyl (4-((2-((tert-butoxycarbonyl)amino)phenyl)carbamoyl)phenyl)-carbamate (4)**

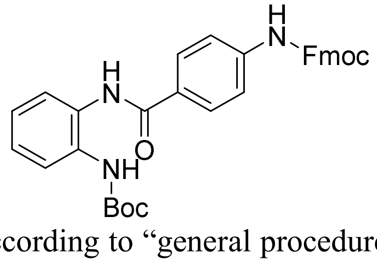


The synthesis was performed according to "general procedure for amide coupling". 4-((((9*H*-fluoren-9-yl)methoxy)carbonyl)amino)benzoic acid (**3**, 5.30 g, 14.8 mmol, 1.0 eq) and **2** (3.07 g, 14.8 mmol, 1.0 eq) were used. The crude product (12.74 g (64% purity), quant yield assumed) was used in the next step without further purification. MS (ESI): m/z calc. for $[C_{33}H_{31}N_3O_5+Na^+]^+$ = 572.22, found = 572.20

**tert-Butyl (2-(4-aminobenzamido)phenyl)carbamate (5)**

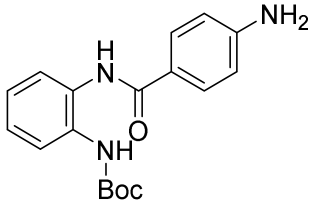

**4** (12.74 g, 64% purity, 14.8 mmol) was dissolved in ACN/morpholine (1/1, 200 mL) and stirred at ambient temperature for 16 h. Afterwards, ice water was added and the resulting precipitate was filtered. The filtrate was extracted with DCM, the organic phase was washed with brine, dried over $MgSO_4$ and volatiles were removed under reduced pressure. The crude product was purified by flash chromatography (silica, DCM/MeOH) to provide the title compound as an off-white solid (3.43 g, 71%). $^1$H NMR (400 MHz, DMSO-$d_6$) δ 9.48 (s, 1H), 8.65 (s, 1H), 7.73 – 7.65 (m, 2H), 7.55 – 7.52 (m, 1H), 7.50 – 7.46 (m, 1H), 7.14 (ddd, *J* = 6.6, 3.7, 2.1 Hz, 2H), 6.65 – 6.59 (m, 2H), 5.81 (s, 2H), 1.46 (s, 9H). MS (ESI): m/z calc. for $[C_{18}H_{21}N_3O_3+H^+]^+$ = 328.16, found = 328.20

**(*E*)-4-((2-Amino-4-hydroxy-5-methylphenyl)diazenyl)-*N*-(2-aminophenyl)benzamide NB161 (1)**

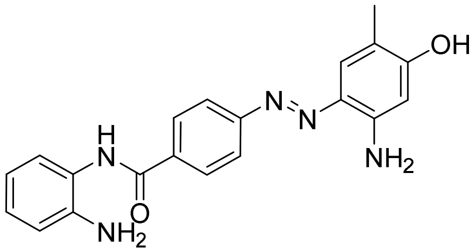

The synthesis was performed according to "general procedure for azo coupling". *tert*-Butyl (2-(4-aminobenzamido)phenyl)carbamate (**5**, 75 mg, 0.23 mmol, 1.0 eq) and 5-amino-2-methylphenol (27 mg, 0.23 mmol, 1.0 eq) were used. The crude product was purified by flash chromatography (hexane/ethyl acetate) to provide the *Boc*-protected intermediate as a red solid (90 mg, 85%). $^1$H NMR (400 MHz, DMSO-$d_6$) δ 10.22 (s, 1H), 9.87 (s, 1H), 8.69 (s, 1H), 8.05 (d, *J* = 8.5 Hz, 2H), 7.87 (d, *J* = 8.2 Hz, 2H), 7.55 (ddd, *J* = 7.8, 4.5, 1.7 Hz, 2H), 7.43 (d, *J* = 1.1 Hz, 1H), 7.21 (td, *J* = 7.7, 1.8 Hz, 1H), 7.16 (td, *J* = 7.4, 1.6 Hz, 1H), 7.12 (br s, 2H), 6.24 (s, 1H), 2.04 (s, 3H), 1.46 (s, 9H). MS (ESI): m/z calc. for $[C_{25}H_{27}N_5O_4+H^+]^+$ = 462.21, found = 462.27

Deprotection of *tert*-butyl (*E*)-(2-(4-((2-amino-4-hydroxy-5-methylphenyl)diazenyl)-benzamido)phenyl)carbamate (85 mg, 0.18 mmol) was performed according to "general procedure for *N-Boc* deprotection". The crude product was purified by flash chromatography (C18 silica, ACN/$H_2O$) to provide the title compound as a red solid (42 mg, 63%). $^1$H NMR (500 MHz, DMSO-$d_6$) δ 10.20 (s, 1H), 9.71 (s, 1H), 8.08 (d, *J* = 8.2 Hz, 2H), 7.86 (d, *J* = 8.2 Hz, 2H), 7.44 (s, 1H), 7.19 (d, *J* = 7.8 Hz, 1H), 7.09 (s, 2H), 6.98 (td, *J* = 7.6, 1.6 Hz, 1H), 6.80 (dd, *J* = 8.0, 1.5 Hz, 1H), 6.61 (td, *J* = 7.5, 1.4 Hz, 1H), 6.26 (s, 1H), 4.92 (s, 2H), 2.05 (s, 3H). MS (ESI): m/z calc. for $[C_{20}H_{19}N_5O_2+H^+]^+$ = 362.15, found = 362.18. HRMS (MALDI): m/z calc. for $[C_{20}H_{19}N_5O_2+H^+]^+$ = 362.1612, found = 362.1613

**2,2-Dimethyl-1-(6-methyl-1*H*-indol-1-yl)propan-1-one**

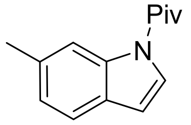


The synthesis was performed following a published protocol (1). $^{1}H$ NMR (400 MHz, DMSO-$d_6$) δ 8.24 – 8.21 (m, 1H), 7.99 (d, $J$ = 3.8 Hz, 1H), 7.47 (d, $J$ = 7.9 Hz, 1H), 7.09 (ddd, $J$ = 7.9, 1.6, 0.7 Hz, 1H), 6.67 (dd, $J$ = 3.9, 0.8 Hz, 1H), 2.41 (s, 3H), 1.44 (s, 9H). MS (ESI): m/z calc. for $[C_{14}H_{17}NO+H^+]^+$ = 216.13, found = 216.10

**6-Methyl-1*H*-indol-7-ol**

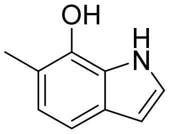


The synthesis was performed according to literature (2). MS (ESI): m/z calc. for $[C_9H_9NO+H^+]^+$ = 148.07, found = 148.10

**(*E*)-*N*-(2-Aminophenyl)-4-((7-hydroxy-6-methyl-1*H*-indol-4-yl)diazenyl)benzamide NB390 (6)**

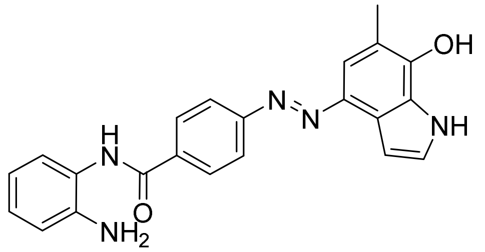


The synthesis was performed according to "general procedure for azo coupling". *tert*-Butyl (2-(4-aminobenzamido)phenyl)carbamate (160 mg, 489 µmol, 1.0 eq) and 6-methyl-1*H*-indol-7-ol (86 mg, 0.59 mmol, 1.2 eq) were used. The *Boc*-protected intermediate was used in the next step without further purification. MS (ESI): m/z calc. for $[C_9H_9NO+H^+]^+$ = 486.21, found = 486.20

Deprotection of crude *tert*-butyl (*E*)-(2-(4-((7-hydroxy-6-methyl-1*H*-indol-4-yl)diazenyl)benzamido)phenyl)carbamate (max. 237 mg, 488 µmol) was performed according to "general procedure for *N-Boc* deprotection. The crude product was purified by flash chromatography (C18 silica, ACN/$H_2O$) to provide the title compound as a red solid (19 mg, 8%).$^{1}H$ NMR (500 MHz, DMSO-$d_6$) δ 11.93 (s, 1H), 11.30 (s, 1H), 9.52 (s, 1H), 7.99 (d, $J$ = 8.6 Hz, 2H), 7.81 (s, 1H), 7.59 (d, $J$ = 8.7 Hz, 1H), 7.46 (d, $J$ = 8.8 Hz, 2H), 7.18 (d, $J$ = 3.0 Hz, 1H), 6.97 (t, $J$ = 7.6 Hz, 1H), 6.79 (dd, $J$ = 7.9, 1.4 Hz, 1H), 6.63 (t, $J$ = 2.5 Hz, 1H), 6.60 (dd, $J$ = 7.5, 1.5 Hz, 1H), 4.90 (br s, 2H), 2.11 (s, 3H). HRMS (MALDI): m/z calc. for $[C_{22}H_{19}N_5O_2+H^+]^+$ = 386.1612, found = 386.1611

**2-Methoxy-4-methyl-3-nitropyridine (8)**

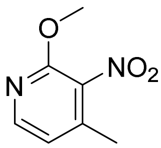


The procedure was adapted from literature (3). Sodium methoxide (100 g, 1.85 mol, 3.8 eq) was dissolved in methanol (300 mL) and cooled to 0 °C. A solution of 2-chloro-4-methyl-3-nitropyridine (**7**, 85.0 g, 0.493 mol, 1.0 eq) in methanol (400 mL) was added dropwise, and the mixture was heated to reflux for 16 h. About half of the solvent was reduced under reduced pressure, and ice water was added. The resulting precipitate was filtered and dried to provide a beige solid (79.5 g, 96%). $^{1}$H NMR (400 MHz, DMSO-$d_6$) δ 8.34 – 8.05 (m, 1H), 7.09 (d, $J$ = 5.7 Hz, 1H), 3.95 (s, 3H), 2.29 (s, 3H). $^{13}$C NMR (101 MHz, DMSO) δ 154.19, 148.05, 141.69, 135.62, 119.47, 54.34, 16.06.

**5-Bromo-2-methoxy-4-methyl-3-nitropyridine (9)**

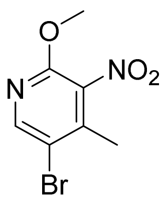


(3)

The procedure was adapted from literature (3). Bromine (65.0 mL, 1.27 mol, 2.7 eq) was slowly added to a suspension of 2-Methoxy-4-methyl-3-nitropyridine (**8**, 79.0 g, 0.470 mol, 1.0 eq) and sodium acetate (139 g, 1.69 mol, 3.6 eq) in acetic acid (450 mL) and the mixture was heated at 80 °C for 16 h. Afterwards, ice water and sat. aq $Na_2SO_3$ were added (1.8 L), and the resulting precipitate was filtered and dried to provide a beige solid (100.5 g, 87%). $^{1}$H NMR (400 MHz, DMSO-$d_6$) δ 8.52 (s, 1H), 3.97 (s, 3H), 2.31 (s, 3H). $^{13}$C NMR (101 MHz, DMSO) δ 153.46, 149.19, 140.97, 135.78, 114.36, 54.87, 17.44.

**(*E*)-2-(5-Bromo-2-methoxy-3-nitropyridin-4-yl)-*N*,*N*-dimethylethen-1-amine (10)**

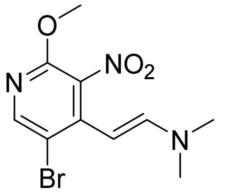


The procedure was adapted from literature (3). 5-Bromo-2-methoxy-4-methyl-3-nitropyridine (**9**, 100 g, 0.405 mol, 1.0 eq) was dissolved in DMF (700 mL) and heated at 80 °C. *N*,*N*-Dimethylformamide dimethyl acetal (500 mL, 3.75 mol, 9.3 eq) was added dropwise, and the mixture was heated at 90 °C for 16 h. After cooling to ambient temperature, water (3 L) was added, and the resulting precipitate was filtered and dried to provide a red solid (120 g, 98%). $^{1}$H NMR (300 MHz, DMSO-$d_6$) δ 8.22 (s, 1H), 7.04 (d, $J$ = 13.5 Hz, 1H), 4.80 (d, $J$ = 13.5 Hz, 1H), 3.88 (s, 3H), 2.90 (s, 6H). $^{13}$C NMR (75 MHz, DMSO) δ 154.49, 148.74, 148.26, 139.83, 111.26, 86.27, 54.84.

**4-Bromo-7-methoxy-1*H*-pyrrolo[2,3-*c*]pyridine (11)**

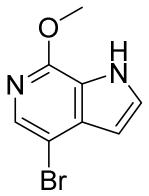

The procedure was adapted from literature (3). A mixture of (*E*)-2-(5-bromo-2-methoxy-3-nitropyridin-4-yl)-*N*,*N*-dimethylethen-1-amine (**10**, 60.0 g, 199 mmol, 1.0 eq) and iron powder (50.0 g, 895 mmol, 4.5 eq) in methanol/acetic acid/water (6/2/1, 450 mL) was heated under reflux for 2 h. The mixture was filtered over a plug of celite, and the solvent was removed under reduced pressure. The residue was partitioned between water and ethyl acetate. The organic layer was washed with brine, dried over $MgSO_4$, filtered, and volatiles were removed under reduced pressure to provide an off-white solid. The procedure was performed twice to provide a combined yield of 85 g (94%). $^{1}$H NMR (300 MHz, DMSO-$d_6$) δ 12.15 (s, 1H), 7.76 (s, 1H), 7.54 (t, $J$ = 2.7 Hz, 1H), 6.54 – 6.34 (m, 1H), 4.02 (s, 3H). $^{13}$C NMR (75 MHz, DMSO) δ 150.80, 134.83, 134.10, 129.13, 120.92, 105.17, 101.98, 53.51.

**4-Bromo-7-methoxy-1-tosyl-1*H*-pyrrolo[2,3-*c*]pyridine (12)**

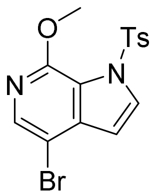

The procedure was adapted from literature (3). Sodium hydride (60%, 28.8 g, 1.20 mol, 3.2 eq) was slowly added to a solution of 4-bromo-7-methoxy-1*H*-pyrrolo[2,3-*c*]pyridine (**11**, 85.0 g, 374 mmol, 1.0 eq) in THF (900 mL) at 0 °C. After stirring for 30 min, tosyl chloride (99.9 g, 524 mmol, 1.4 eq) was slowly added. After 2 h, the reaction was quenched by adding ice water and extracted with ethyl acetate. The organic layer was washed with brine, dried over $MgSO_4$, filtered, and volatiles were removed under reduced pressure. The residue was recrystallized from acetonitrile to provide a beige solid (110 g, 77%). $^{1}$H NMR (400 MHz, DMSO-$d_6$) δ 8.15 (d, $J$ = 3.7 Hz, 1H), 7.94 (s, 1H), 7.84 (d, $J$ = 8.4 Hz, 2H), 7.41 (d, $J$ = 8.3 Hz, 2H), 6.75 (d, $J$ = 3.7 Hz, 1H), 3.81 (s, 3H), 2.33 (s, 3H). $^{13}$C NMR (101 MHz, DMSO) δ 149.95, 145.53, 139.14, 138.55, 134.93, 132.20, 129.89, 127.72, 118.37, 105.78, 104.14, 53.18, 21.05.

**4-Bromo-1-tosyl-1,6-dihydro-7*H*-pyrrolo[2,3-*c*]pyridin-7-one (13)**

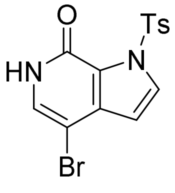

(4)

The procedure was adapted from literature (4). 4-Bromo-7-methoxy-1-tosyl-1*H*-pyrrolo[2,3-*c*]pyridine (**12**, 80.0 g, 210 mmol) was dissolved in dioxane (300 mL) and HCl in dioxane (4 M, 250 mL) and heated at 50 °C for 2 h. About 300 mL of dioxane were removed under reduced pressure, and the residue was triturated with diethyl ether. The precipitate was filtered and dried to provide an off-white solid (62 g, 81%). $^{1}$H NMR (400 MHz, DMSO-$d_6$) δ 11.63 (s, 1H), 8.02 (d, $J$ = 3.5 Hz, 1H), 7.93 (d, $J$ = 8.4 Hz, 2H), 7.39 (d, $J$ = 8.2 Hz, 2H), 7.34 (s,

1H), 6.58 (d, $J$ = 3.5 Hz, 1H), 2.35 (s, 3H). $^{13}$C NMR (101 MHz, DMSO) δ 151.97, 145.36, 136.70, 135.03, 131.11, 129.67, 129.60, 128.39, 121.88, 106.24, 91.19, 21.09.

**4-Bromo-6-methyl-1-tosyl-1,6-dihydro-7*H*-pyrrolo[2,3-*c*]pyridin-7-one (14)**

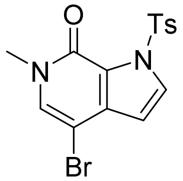


The procedure was adapted from literature (4). Sodium hydride (60%, 3.63 g, 151 mmol, 1.5 eq) was slowly added to a solution of 4-bromo-7-methoxy-1*H*-pyrrolo[2,3-*c*]pyridine (**13**, 37.0 g, 101 mmol, 1.0 eq) in DMF (400 mL) at 0 °C. After stirring for 20 min, iodomethane (9.41 mL, 151 mmol, 1.5 eq) was slowly added. After 2 h, the reaction was quenched by adding ice water. The resulting precipitate was filtered and dried to provide an off-white solid (36.37 g, 95%). $^{1}$H NMR (400 MHz, DMSO-$d_6$) δ 8.05 (d, $J$ = 3.5 Hz, 1H), 7.95 (d, $J$ = 8.4 Hz, 2H), 7.78 (s, 1H), 7.41 (d, $J$ = 8.2 Hz, 2H), 6.58 (d, $J$ = 3.5 Hz, 1H), 3.39 (s, 3H), 2.36 (s, 3H). $^{13}$C NMR (101 MHz, DMSO) δ 151.78, 145.30, 136.10, 135.09, 134.17, 131.38, 129.62, 128.40, 121.42, 105.95, 90.79, 36.31, 21.08.

**6-Methyl-4-(4,4,5,5-tetramethyl-1,3,2-dioxaborolan-2-yl)-1-tosyl-1,6-dihydro-7*H*-pyrrolo-[2,3-*c*]pyridin-7-one (15)**

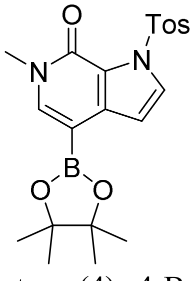


The procedure was adapted from literature (4). 4-Bromo-6-methyl-1-tosyl-1,6-dihydro-7*H*-pyrrolo[2,3-*c*]pyridin-7-one (**14**, 10.0 g, 26.2 mmol, 1.0 eq), 4,4,4′,4′,5,5,5′,5′-octamethyl-2,2′-bi(1,3,2-dioxaborolane) (13.3 g, 52.4 mmol, 2.0 eq), potassium acetate (5.66 g, 57.7 mmol, 2.2 eq), Pd XPhos G2 (825 mg, 1.05 mmol, 0.04 eq) and XPhos (125 mg, 262 µmol, 0.01 eq) were dissolved in dioxane (150 mL) and purged with argon for 10 min. The mixture was heated at 80 °C for 2 h, cooled to ambient temperature, partitioned between water and ethyl acetate and filtered over a plug of celite. The ethyl acetate layer was washed with brine, dried over $MgSO_4$, filtered, and volatiles were removed under reduced pressure. The crude residue was triturated with hexane/diethyl ether (2/1), filtered and washed with hexane to provide the title compound as a colorless solid (8.68 g, 77%). $^{1}$H NMR (400 MHz, DMSO-$d_6$) δ 7.97 (d, $J$ = 3.5 Hz, 1H), 7.91 (d, $J$ = 1.8 Hz, 1H), 7.89 (d, $J$ = 1.8 Hz, 1H), 7.72 (s, 1H), 7.42 – 7.40 (m, 1H), 7.42 – 7.36 (m, 1H), 6.81 (d, $J$ = 3.5 Hz, 1H), 3.43 (s, 3H), 2.36 (s, 3H), 1.29 (s, 12H). $^{13}$C NMR (101 MHz, DMSO) δ 152.86, 144.91, 143.05, 138.93, 135.53, 131.06, 129.55, 129.52, 128.11, 121.29, 107.80, 82.80, 36.29, 24.60, 21.06. MS (ESI): m/z calc. for $[C_{21}H_{25}BN_2O_5S+H^+]^+$ = 429.16, found = 429.10

**6-(Methoxycarbonyl)quinoline 1-oxide (17)**

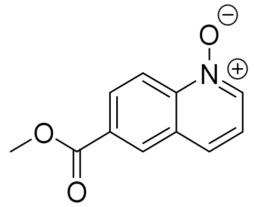

The synthesis was adapted from literature (5). Methyl quinoline-6-carboxlate (**16**, 8.00 g, 42.7 mmol, 1.0 eq) was dissolved in DCM (200 mL) and cooled to 0 °C. *meta*-chloroperoxybenzoic acid (75 %, wet with water; 19.67 g, 85.47 mmol, 2.0 eq) was added in small portions to the stirring solution. After addition, the cooling bath was removed, and the solution was stirred for 3 h at ambient temperature. Afterwards, the reaction mixture was quenched with sat. aq $NaHCO_3$, and the aqueous layer was extracted with DCM. The combined organic layers were washed with brine, dried over $MgSO_4$, filtered, and the solvent was removed under reduced pressure to provide a colorless solid (7.97 g, 92%). $^1$H NMR (400 MHz, DMSO-$d_6$) δ 8.70 (d, $J$ = 1.6 Hz, 1H), 8.67 (d, $J$ = 5.9 Hz, 1H), 8.58 (d, $J$ = 9.0 Hz, 1H), 8.18 (dd, $J$ = 9.0, 1.6 Hz, 1H), 8.09 (d, $J$ = 8.3 Hz, 1H), 7.54 (dd, $J$ = 8.5, 5.8 Hz, 1H), 3.92 (s, 3H). $^{13}$C NMR (101 MHz, DMSO) δ 165.26, 142.43, 136.99, 131.09, 129.85, 129.43, 128.99, 126.06, 122.87, 119.75, 52.58. MS (ESI): m/z calc. for $[C_{11}H_9NO_3+H^+]^+$ = 204.06, found = 204.05

**Methyl 2-hydroxyquinoline-6-carboxylate (18)**

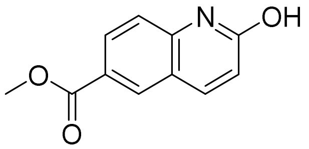

The synthesis was adapted from literature (6). Mesyl chloride (6.03 mL, 77.7 mmol, 2.0 eq) was added to a stirred solution of 6-(methoxycarbonyl)quinoline 1-oxide (**17**, 7.90 g, 38.9 mmol, 1.0 eq) in acetonitrile/water (1/1, 140 mL), and the mixture was stirred at ambient temperature for 45 min. The resulting precipitate was filtered, washed with hexane and dried to give a colorless solid (5.68 g, 72%). $^1$H NMR (400 MHz, DMSO-$d_6$) δ 12.03 (s, 1H), 8.29 (d, $J$ = 1.9 Hz, 1H), 8.07 – 7.97 (m, 2H), 7.35 (d, $J$ = 8.6 Hz, 1H), 6.56 (d, $J$ = 9.6 Hz, 1H), 3.85 (s, 3H). $^{13}$C NMR (101 MHz, DMSO) δ 165.66, 162.03, 142.08, 140.37, 130.54, 129.86, 122.82, 122.72, 118.63, 115.37, 52.01. MS (ESI): m/z calc. for $[C_{11}H_9NO_3+H^+]^+$ = 204.06, found = 204.10

**Methyl 2-chloroquinoline-6-carboxylate (19)**

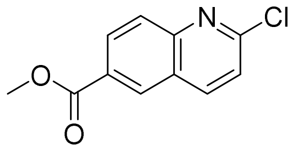

Methyl 2-hydroxyquinoline-6-carboxylate (**18**, 5.64 g, 27.8 mmol) was suspended in DCM (80 mL) and cooled to 0 °C. DMF (5.0 mL) and $SOCl_2$ (5.0 mL, 68.9 mmol, 2.5 eq) were subsequently added, and the mixture was stirred for 18 h. The reaction was quenched with ice and sat. aq $NaHCO_3$ and extracted with DCM. The organic phase was washed with sat. aq $NaHCO_3$ and brine, dried over $MgSO_4$, filtered, and the solvent was removed under reduced pressure. The crude product (8.80 g) was recrystallized from chloroform to provide the title compound as a colorless solid (4.28 g, 70%). $^{1}H$ NMR (400 MHz, DMSO-$d_6$) δ 8.73 (d, $J$ = 2.0 Hz, 1H), 8.64 (dd, $J$ = 8.7, 0.8 Hz, 1H), 8.24 (dd, $J$ = 8.8, 2.0 Hz, 1H), 8.02 (d, $J$ = 8.8 Hz, 1H), 7.69 (d, $J$ = 8.6 Hz, 1H), 3.93 (s, 3H). $^{13}C$ NMR (101 MHz, DMSO) δ 165.55, 152.28, 148.94, 141.29, 130.79, 129.77, 128.43, 127.90, 126.10, 123.41, 52.49. MS (ESI): m/z calc. for $[C_{11}H_8ClNO_2+H^+]^+$ = 222.02, found = 222.00

**Methyl 2-(6-methyl-7-oxo-1-tosyl-6,7-dihydro-1*H*-pyrrolo[2,3-*c*]pyridin-4-yl)quinoline-6-carboxylate (20)**

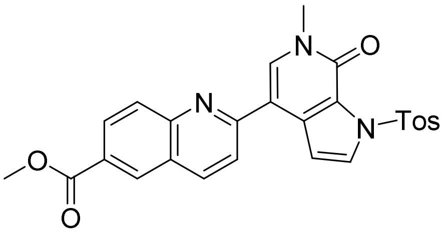


The synthesis was performed according to “general procedure for Suzuki coupling B”. **15** (1.20 g, 2.80 mmol, 1.0 eq) and methyl 2-chloroquinoline-6-carboxylate (**19**, 869 mg, 3.92 mmol, 1.4 eq) were used. The crude product could be filtered from the reaction mixture to give a colorless solid (1.27 g, 93%). The compound was not soluble enough in DMSO to give an NMR spectrum. MS (ESI): m/z calc. for $[C_{26}H_{21}N_3O_5S+H^+]^+$ = 488.12, found = 488.15

**2-(6-Methyl-7-oxo-6,7-dihydro-1*H*-pyrrolo[2,3-*c*]pyridin-4-yl)quinoline-6-carboxylic acid (21)**

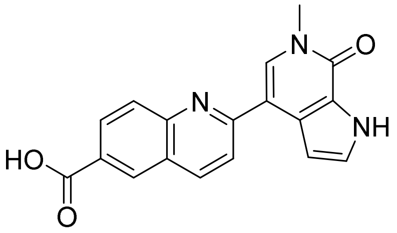


The synthesis was performed according to “general procedure for ester and tosylamide hydrolysis”. Methyl 2-(6-methyl-7-oxo-1-tosyl-6,7-dihydro-1*H*-pyrrolo[2,3-*c*]pyridin-4-yl)quinoline-6-carboxylate (**20**, 1.20 g, 2.46 mmol) was used. The resulting yellow solid (0.85 g, quant) was not soluble enough in DMSO to give an NMR spectrum. MS (ESI): m/z calc. for $[C_{18}H_{13}N_3O_3+H^+]^+$ = 320.10, found = 320.10

***N*-(2-Aminophenyl)-2-(6-methyl-7-oxo-6,7-dihydro-1*H*-pyrrolo[2,3-*c*]pyridin-4-yl)quinoline-6-carboxamide NB437 (22)**

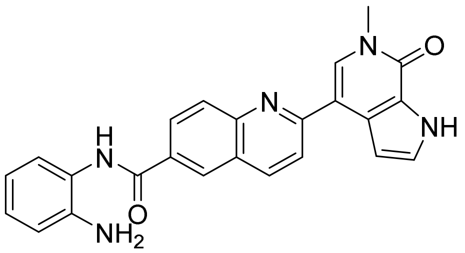

The synthesis was performed according to "general procedure for amide coupling". **21** (250 mg, 783 µmol, 1.0 eq) and *tert*-butyl (2-aminophenyl)carbamate (179 mg, 861 µmol, 1.1 eq) were used. The *Boc*-protected intermediate (399 mg, quant) was used in the next step without further purification. MS (ESI): m/z calc. for $[C_{29}H_{27}N_5O_4+H^+]^+$ = 510.21, found = 510.25

The next step was performed according to "general procedure for *N-Boc* deprotection". *tert*-Butyl (2-(2-(6-methyl-7-oxo-6,7-dihydro-1*H*-pyrrolo[2,3-*c*]pyridin-4-yl)quinoline-6-carboxamido)-phenyl)carbamate (399 mg, 783 µmol) was dissolved in DCM/TFA (10 mL, 3/1). The crude product was purified by flash chromatography (DCM/MeOH) to give a beige solid (124 mg, 39%). $^1$H NMR (500 MHz, DMSO-$d_6$) δ 12.18 (s, 1H), 9.88 (s, 1H), 8.63 (d, *J* = 2.1 Hz, 1H), 8.49 (d, *J* = 8.7 Hz, 1H), 8.30 (dd, *J* = 8.7, 2.1 Hz, 1H), 8.27 (s, 1H), 8.15 (dd, *J* = 8.8, 2.2 Hz, 2H), 7.43 (t, *J* = 2.8 Hz, 1H), 7.38 (t, *J* = 2.4 Hz, 1H), 7.25 (dd, *J* = 7.9, 1.5 Hz, 1H), 7.00 (td, *J* = 7.7, 1.6 Hz, 1H), 6.82 (dd, *J* = 8.1, 1.5 Hz, 1H), 6.63 (td, *J* = 7.5, 1.5 Hz, 1H), 4.99 (s, 2H), 3.69 (s, 3H). $^{13}$C NMR (126 MHz, DMSO) δ 157.54, 154.91, 149.22, 143.67, 137.71, 132.65, 132.12, 129.12, 129.06, 128.93, 128.66, 128.17, 127.63, 127.19, 127.04, 125.85, 123.99, 123.77, 119.96, 116.73, 116.61, 113.69, 105.65, 36.47. MS (ESI): m/z calc. for $[C_{24}H_{19}N_5O_2+H^+]^+$ = 410.15, found = 410.10 HRMS (MALDI): m/z calc. for $[C_{24}H_{19}N_5O_2+H^+]^+$ = 410.1612, found = 410.1618

**6-Methyl-7-oxo-1-tosyl-6,7-dihydro-1*H*-pyrrolo[2,3-*c*]pyridine-4-carboxylic acid (23)**

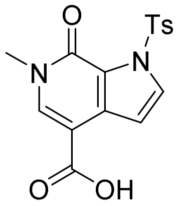


**14** (3.00 g, 7.87 mmol, 1.0 eq) was dissolved in anh. THF (50 mL) and cooled to -40 °C. A solution of isopropylmagnesium chloride lithium chloride complex in THF (1.3 M, 12.1 mL, 15.7 mmol, 2.0 eq) was slowly added. The mixture was stirred for 2 h and poured onto powdered dry ice. After stirring for 30 minutes, the reaction was quenched by adding sat. aq $NH_4Cl$ and extracted with ethyl acetate. The ethyl acetate layer was washed with brine, dried over $MgSO_4$, filtered, and volatiles were removed under reduced pressure to provide the crude product (2.73 g, quant.). MS (ESI): m/z calc. for $[C_{16}H_{14}N_2O_5S+H^+]^+$ = 347.06, found = 347.00

**6-Methyl-7-oxo-6,7-dihydro-1*H*-pyrrolo[2,3-*c*]pyridine-4-carboxylic acid (24)**

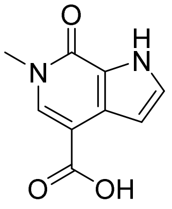


**23** (2.73 g, 7.88 mmol, 1 eq) and LiOH · $H_2O$ (2.65 g, 63.1 mmol, 8 eq) were dissolved in dioxane/water (60 mL, 3/1) and heated at 90 °C for 1 h. The mixture was cooled to ambient temperature and brought to a pH of 3 by the addition of 5% aq HCl. The resulting precipitate was filtered and dried under reduced pressure to provide the product as a colorless solid (1.27 g, 84 %). $^1$H NMR (400 MHz, DMSO-$d_6$) δ 12.50 (s, 1H), 12.13 (s, 1H), 8.07 (s, 1H), 7.34 (t, *J* = 2.8 Hz, 1H), 6.73 (dd, *J* = 2.7, 2.1 Hz, 1H), 3.59 (s, 3H). $^{13}$C NMR (101 MHz, DMSO) δ

166.58, 154.72, 136.59, 127.70, 127.59, 122.58, 104.78, 104.03, 36.02. MS (ESI): m/z calc. for $[C_9H_8N_2O_3+H^+]^+$ = 193.05, found = 193.00

**Methyl 4-(6-methyl-7-oxo-6,7-dihydro-1*H*-pyrrolo[2,3-*c*]pyridine-4-carboxamido)benzoate (25)**

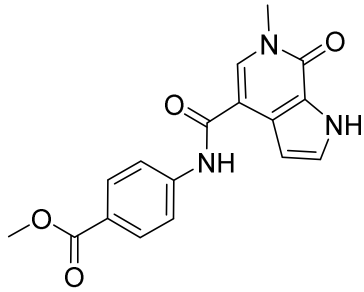

**24** (350 mg, 1.81 mmol, 1.0 eq) and $SOCl_2$ (211 µL, 2.91 mmol, 1.6 eq) were added to dioxane (15 mL). The mixture was heated at 80 °C for 16 h and cooled to ambient temperature. Then, a solution of methyl 4-aminobenzoate (441 mg, 2.91 mmol, 1.6 eq) and *N*,*N*-diisopropylethylamine (381 µL, 2.19 mmol, 1.2 eq) in *N*,*N*-dimethylacetamide (8 mL) was added. The mixture was stirred for 1 h and partitioned between water and ethyl acetate. The ethyl acetate layer was washed with brine, dried over $MgSO_4$, filtered, and volatiles were removed under reduced pressure. The crude product was purified by flash chromatography (DCM -> 10% MeOH in DCM) to provide the title compound as a colorless solid (550 mg, 93 %). $^1$H NMR (400 MHz, DMSO-$d_6$) δ 12.17 (s, 1H), 10.25 (s, 1H), 8.11 (s, 1H), 7.98 – 7.93 (m, 2H), 7.90 – 7.86 (m, 2H), 7.37 (t, *J* = 2.7 Hz, 1H), 6.74 (dd, *J* = 2.7, 2.0 Hz, 1H), 3.84 (s, 3H), 3.61 (s, 3H). $^{13}$C NMR (101 MHz, DMSO) δ 165.87, 164.48, 154.49, 143.94, 133.34, 130.14, 127.59, 127.55, 123.77, 122.81, 119.12, 108.77, 103.77, 51.84, 36.05. MS (ESI): m/z calc. for $[C_{17}H_{15}N_3O_4+H^+]^+$ = 326.11, found = 326.05

**4-(6-Methyl-7-oxo-6,7-dihydro-1*H*-pyrrolo[2,3-*c*]pyridine-4-carboxamido)benzoic acid (26)**

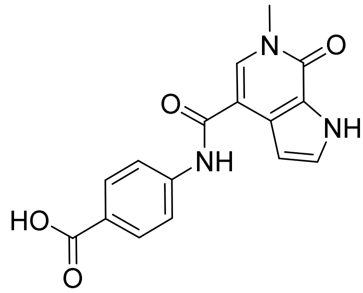

**25** (425 mg, 1.31 mmol, 1 eq) and LiOH · $H_2O$ (329 mg, 7.84 mmol, 6 eq) were dissolved in THF/methanol/water (40 mL, 3/3/2) and heated at 60 °C for 1 h. The mixture was cooled to ambient temperature and brought to a pH of 3 by the addition of 5% aq HCl. The resulting precipitate was filtered and dried under reduced pressure to provide the product as an off-white solid (397 mg, 98%). $^1$H NMR (400 MHz, DMSO-$d_6$) δ 12.70 (s, 1H), 12.18 (s, 1H), 10.27 (s, 1H), 8.16 (s, 1H), 7.97 – 7.93 (m, 2H), 7.91 – 7.86 (m, 2H), 7.38 (t, *J* = 2.8 Hz, 1H), 6.76 (t, *J* = 2.4 Hz, 1H), 3.63 (s, 3H). $^{13}$C NMR (101 MHz, DMSO) δ 166.97, 164.45, 154.49, 143.58, 133.32, 130.24, 127.64, 127.54, 124.96, 122.81, 119.04, 108.80, 103.79, 36.03. MS (ESI): m/z calc. for $[C_{16}H_{13}N_3O_4+H^+]^+$ = 312.09, found = 312.05

**_N_-(4-((2-Aminophenyl)carbamoyl)phenyl)-6-methyl-7-oxo-6,7-dihydro-1_H_-pyrrolo[2,3-_c_]pyridine-4-carboxamide NB480 (27)**

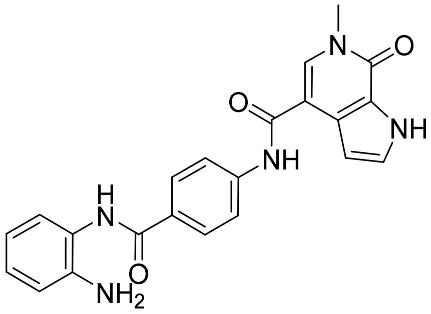


The synthesis was performed according to "general procedure for amide coupling". **26** (190 mg, 793 µmol, 1.0 eq) and benzene-1,2-diamine (86 mg, 0.79 mmol, 1.3 eq) were used. The crude product was triturated with methanol and filtered to provide the title compound as a colorless solid (161 mg, 66%). $^1$H NMR (500 MHz, DMSO-$d_6$) δ 12.18 (s, 1H), 10.18 (s, 1H), 9.60 (s, 1H), 8.12 (s, 1H), 8.01 (d, $J$ = 8.4 Hz, 2H), 7.86 (d, $J$ = 8.5 Hz, 2H), 7.38 (t, $J$ = 2.8 Hz, 1H), 7.19 (d, $J$ = 7.2 Hz, 1H), 6.98 (td, $J$ = 7.7, 1.6 Hz, 1H), 6.80 (dd, $J$ = 8.0, 1.5 Hz, 1H), 6.76 (t, $J$ = 2.4 Hz, 1H), 6.61 (td, $J$ = 7.5, 1.4 Hz, 1H), 4.89 (s, 2H), 3.63 (s, 3H). $^{13}$C NMR (126 MHz, DMSO) δ 164.78, 164.40, 154.52, 143.13, 142.29, 133.17, 128.89, 128.58, 127.66, 127.55, 126.66, 126.37, 123.54, 122.86, 118.94, 116.31, 116.17, 108.94, 103.82, 36.06. MS (ESI): m/z calc. for $[C_{22}H_{19}N_5O_3+H^+]^+$ = 402.15, found = 402.10. HRMS (MALDI): m/z calc. for $[C_{22}H_{19}N_5O_3+Na^+]^+$ = 424.1380, found = 424.1390

**Methyl 4-(6-methyl-7-oxo-1-tosyl-6,7-dihydro-1_H_-pyrrolo[2,3-_c_]pyridin-4-yl)benzoate (29a)**

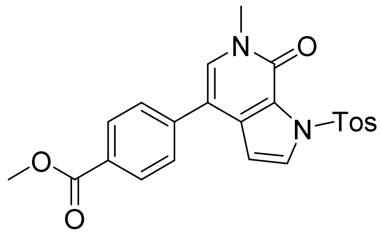


The synthesis was performed according to "general procedure for Suzuki coupling B". **15** (1.00 g, 2.33 mmol, 1.0 eq) and methyl 4-bromobenzoate (**28a**, 603 mg, 2.80 mmol, 1.2 eq) were used. The crude product (1.00 g, 98%) was used in the next step without further characterization. MS (ESI): m/z calc. for $[C_{23}H_{20}N_2O_5S+H^+]^+$ = 437.11, found = 437.05

**Methyl 3-amino-4-(6-methyl-7-oxo-1-tosyl-6,7-dihydro-1_H_-pyrrolo[2,3-_c_]pyridin-4-yl)benzoate (29b)**

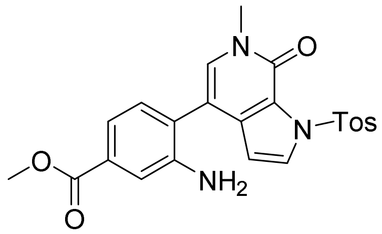


The synthesis was performed according to "general procedure for Suzuki coupling B". **15** (1.60 g, 3.74 mmol, 1.0 eq) and methyl 3-amino-4-bromobenzoate (**28b**, 1.12 g, 4.86 mmol,

1.3 eq) were used. The crude product (1.60 g, 95%) was used in the next step without further characterization. MS (ESI): m/z calc. for $[C_{23}H_{21}N_3O_5S+H^+]^+$ = 452.12, found = 452.10

**Methyl 3-methoxy-4-(6-methyl-7-oxo-1-tosyl-6,7-dihydro-1*H*-pyrrolo[2,3-*c*]pyridin-4-yl)benzoate (29c)**

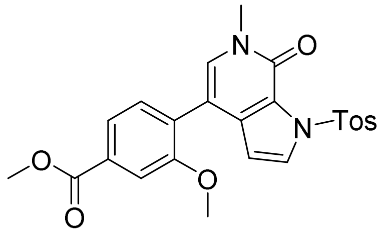


The synthesis was performed according to "general procedure for Suzuki coupling B". **15** (1.00 g, 2.33 mmol, 1.0 eq) and methyl 4-bromo-3-methoxybenzoate (**28c**, 629 mg, 2.57 mmol, 1.1 eq) were used. The crude product (1.00 g, 92%) was used in the next step without further characterization. MS (ESI): m/z calc. for $[C_{24}H_{22}N_2O_6S+H^+]^+$ = 467.00, found = 467.12

**Methyl 5-(6-methyl-7-oxo-1-tosyl-6,7-dihydro-1*H*-pyrrolo[2,3-*c*]pyridin-4-yl)picolinate (29d)**

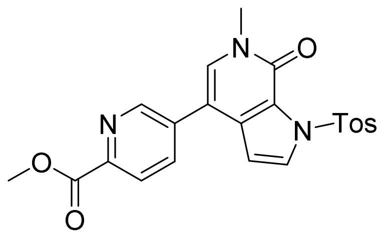


The synthesis was performed according to "general procedure for Suzuki coupling B". **15** (3.50 g, 8.17 mmol, 1.0 eq) and methyl 5-chloropicolinate (**28d**, 1.68 g, 9.81 mmol, 1.2 eq) were used. After completion of the reaction, additional water was added, and the resulting solid was filtered. Dioxane was removed from the filtrate under reduced pressure, the solution was cooled to 4 °C overnight, and the resulting precipitate was filtered. The combined solids were dried under reduced pressure to provide the crude product (1.90 g, 53%). $^1$H NMR (400 MHz, DMSO-$d_6$) δ 8.86 (s, 1H), 8.13 (s, 2H), 8.08 (d, $J$ = 3.6 Hz, 1H), 7.97 (d, $J$ = 8.0 Hz, 2H), 7.85 (s, 1H), 7.43 (d, $J$ = 8.2 Hz, 2H), 6.79 (d, $J$ = 3.6 Hz, 1H), 3.91 (s, 3H), 3.49 (s, 3H), 2.38 (s, 3H). MS (ESI): m/z calc. for $[C_{22}H_{19}N_3O_5S+H^+]^+$ = 438.10, found = 438.00

**Methyl 6-(6-methyl-7-oxo-1-tosyl-6,7-dihydro-1*H*-pyrrolo[2,3-*c*]pyridin-4-yl)pyridazine-3-carboxylate (29e)**

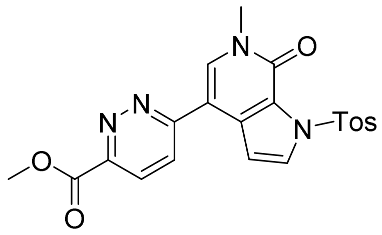

The synthesis was performed according to "general procedure for Suzuki coupling B". **15** (400 mg, 934 µmol, 1.0 eq) and methyl 6-chloropyridazine-3-carboxylate (**28e**, 209 mg, 1.21 mmol, 1.2 eq) were used. MS (ESI): m/z calc. for $[C_{21}H_{18}N_4O_5S+H^+]^+$ = 439.10, found = 439.05
The product was hydrolyzed in the next step without being isolated first.

**Methyl 6-(6-methyl-7-oxo-1-tosyl-6,7-dihydro-1*H*-pyrrolo[2,3-*c*]pyridin-4-yl)nicotinate (29f)**

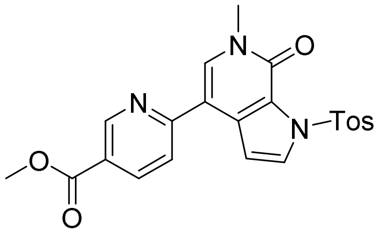

The synthesis was performed according to "general procedure for Suzuki coupling B". **15** (400 mg, 934 µmol, 1.0 eq) and methyl 6-chloronicotinate (**28f**, 208 mg, 1.21 mmol, 1.2 eq) were used. MS (ESI): m/z calc. for $[C_{22}H_{19}N_3O_5S+H^+]^+$ = 438.10, found = 438.05
The product was hydrolyzed in the next step without being isolated first.

**4-(6-Methyl-7-oxo-6,7-dihydro-1*H*-pyrrolo[2,3-*c*]pyridin-4-yl)benzoic acid (30a)**

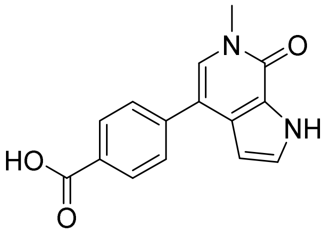

The synthesis was performed according to "general procedure for ester and tosylamide hydrolysis". **29a** (1.00 g, 2.29 mmol) was used. The product was obtained as a colorless solid (602 mg, 98%). $^1$H NMR (500 MHz, DMSO-$d_6$) δ 12.97 (s, 1H), 12.19 (s, 1H), 8.02 (d, $J$ = 8.0 Hz, 2H), 7.73 (d, $J$ = 8.0 Hz, 2H), 7.51 (s, 1H), 7.37 (d, $J$ = 3.1 Hz, 1H), 6.50 (d, $J$ = 2.4 Hz, 1H), 3.60 (s, 3H). $^{13}$C NMR (126 MHz, DMSO) δ 167.19, 154.15, 141.70, 131.70, 131.30, 129.88, 128.94, 128.81, 127.97, 127.31, 127.14, 123.40, 113.71, 102.17, 35.64. MS (ESI): m/z calc. for $[C_{15}H_{12}N_2O_3+H^+]^+$ = 269.08, found = 269.05

**3-Amino-4-(6-methyl-7-oxo-6,7-dihydro-1H-pyrrolo[2,3-c]pyridin-4-yl)benzoic acid (30b)**

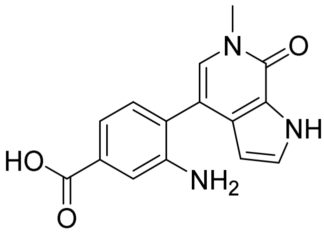

The synthesis was performed according to "general procedure for ester and tosylamide hydrolysis". **29b** (1.60 g, 3.54 mmol) was used. The product (970 mg, 97%) was used in the next step without further characterization. MS (ESI): m/z calc. for $[C_{15}H_{13}N_3O_3+H^+]^+$ = 284.10, found = 284.05

**3-Methoxy-4-(6-methyl-7-oxo-6,7-dihydro-1*H*-pyrrolo[2,3-*c*]pyridin-4-yl)benzoic acid (30c)**

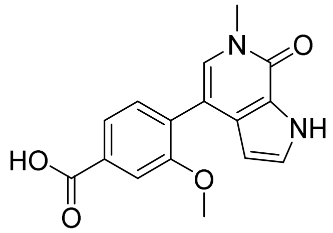


The synthesis was performed according to "general procedure for ester and tosylamide hydrolysis". **29c** (1.00 g, 2.14 mmol) was used. The product was obtained as a colorless solid (621 mg, 97%). $^1$H NMR (400 MHz, DMSO-$d_6$) δ 12.99 (s, 1H), 12.00 (s, 1H), 7.66 – 7.57 (m, 2H), 7.45 (d, $J$ = 7.8 Hz, 1H), 7.30 – 7.24 (m, 2H), 6.07 (t, $J$ = 2.5 Hz, 1H), 3.80 (s, 3H), 3.56 (s, 3H). $^{13}$C NMR (101 MHz, DMSO) δ 167.11, 156.58, 154.11, 130.89, 130.63, 130.31, 129.57, 129.28, 126.62, 122.87, 121.69, 111.66, 111.07, 102.88, 55.35, 35.54. MS (ESI): m/z calc. for $[C_{16}H_{14}N_2O_4+H^+]^+$ = 299.10, found = 299.05

**5-(6-Methyl-7-oxo-6,7-dihydro-1*H*-pyrrolo[2,3-*c*]pyridin-4-yl)picolinic acid (30d)**

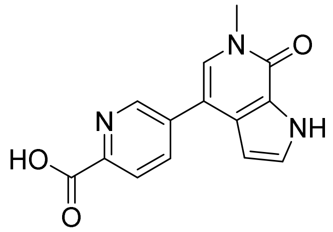


The synthesis was performed according to "general procedure for ester and tosylamide hydrolysis". **29d** (1.85 g, 4.23 mmol) was used. The product was obtained as a yellow solid (1.09 g, 96%). $^1$H NMR (500 MHz, DMSO-$d_6$) δ 12.26 (s, 1H), 8.96 – 8.92 (m, 2H), 8.18 (dd, $J$ = 8.1, 2.3 Hz, 1H), 8.12 (d, $J$ = 7.9 Hz, 1H), 7.65 (s, 1H), 7.40 (t, $J$ = 2.8 Hz, 1H), 6.52 (dd, $J$ = 2.8, 2.0 Hz, 1H), 3.61 (s, 3H). $^{13}$C NMR (126 MHz, DMSO) δ 166.05, 154.18, 147.54, 146.18, 136.09, 135.27, 129.75, 127.65, 127.61, 124.89, 123.41, 110.43, 101.92, 35.74. MS (ESI): m/z calc. for $[C_{14}H_{11}N_3O_3+H^+]^+$ = 270.08, found = 270.00

**6-(6-methyl-7-oxo-6,7-dihydro-1*H*-pyrrolo[2,3-*c*]pyridin-4-yl)pyridazine-3-carboxylic acid (30e)**

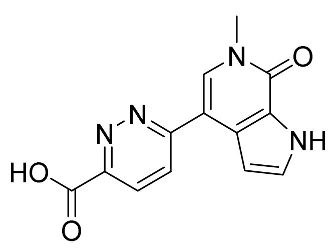


The synthesis was performed according to "general procedure for ester and tosylamide hydrolysis". Hydrolysis was performed directly after Suzuki coupling. LiOH · $H_2O$ (274 mg,

6.53 mmol, 7 eq) was added to the solution of **29e** (409 mg, 933 µmol, 1 eq). The product was obtained as a yellow solid (130 mg, 52%). MS (ESI): m/z calc. for $[C_{13}H_{10}N_4O_3+H^+]^+$ = 271.08, found = 271.05

**6-(6-methyl-7-oxo-6,7-dihydro-1*H*-pyrrolo[2,3-*c*]pyridin-4-yl)nicotinic acid (30f)**

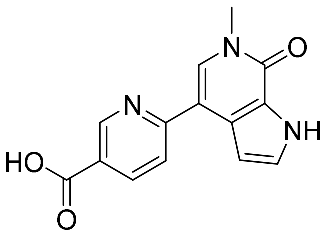


The synthesis was performed according to "general procedure for ester and tosylamide hydrolysis". Hydrolysis was performed directly after Suzuki coupling. LiOH · $H_2O$ (274 mg, 6.53 mmol, 7 eq) was added to the solution of **29f** (409 mg, 933 µmol, 1 eq). The product was obtained as a yellow solid (170 mg, 68%). MS (ESI): m/z calc. for $[C_{14}H_{11}N_3O_3+H^+]^+$ = 270.08, found = 270.00

***N*-(2-Aminophenyl)-4-(6-methyl-7-oxo-6,7-dihydro-1*H*-pyrrolo[2,3-*c*]pyridin-4-yl)benzamide NB462 (31a)**

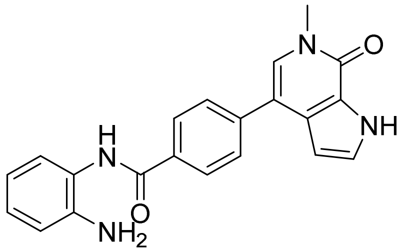


The synthesis was performed according to "general procedure for amide coupling". **30a** (350 mg, 1.30 mmol, 1.0 eq) and benzene-1,2-diamine (169 mg, 1.57 mmol, 1.2 eq) were used. The crude product was triturated with methanol and filtered to provide the title compound as a colorless solid (198 mg, 42%). $^1H$ NMR (500 MHz, DMSO-$d_6$) δ 12.20 (s, 1H), 9.72 (s, 1H), 8.10 (d, $J$ = 7.9 Hz, 2H), 7.74 (d, $J$ = 8.0 Hz, 2H), 7.52 (s, 1H), 7.39 (d, $J$ = 3.7 Hz, 1H), 7.21 (d, $J$ = 7.8 Hz, 1H), 6.99 (t, $J$ = 7.7 Hz, 1H), 6.82 (d, $J$ = 8.0 Hz, 1H), 6.63 (t, $J$ = 7.5 Hz, 1H), 6.51 (s, 1H), 4.94 (s, 2H), 3.62 (s, 3H). $^{13}C$ NMR (126 MHz, DMSO) δ 165.03, 154.16, 143.18, 140.26, 132.67, 128.65, 128.36, 128.12, 127.30, 126.90, 126.74, 126.51, 123.44, 123.42, 116.34, 116.20, 113.87, 102.14, 54.91, 35.64. MS (ESI): m/z calc. for $[C_{21}H_{18}N_4O_2+H^+]^+$ = 359.14, found = 359.10. HRMS (MALDI): m/z calc. for $[C_{21}H_{18}N_4O_2+Na^+]^+$ = 381.1322, found = 381.1332

**3-Amino-*N*-(2-aminophenyl)-4-(6-methyl-7-oxo-6,7-dihydro-1*H*-pyrrolo[2,3-*c*]pyridin-4-yl)benzamide NB469 (31b)**

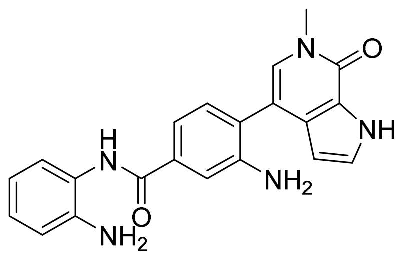


The synthesis was performed according to "general procedure for amide coupling". **30b** (970 mg, 3.42 mmol, 1.0 eq) and benzene-1,2-diamine (555 mg, 5.14 mmol, 1.5 eq) were used. The crude product was triturated with methanol and filtered to provide the title compound as a beige solid (417 mg, 33%). $^1$H NMR (500 MHz, DMSO-$d_6$) δ 12.08 (s, 1H), 9.54 (s, 1H), 7.35 (d, $J$ = 1.8 Hz, 1H), 7.30 (t, $J$ = 2.8 Hz, 1H), 7.22 (ddd, $J$ = 17.0, 7.7, 1.6 Hz, 2H), 7.20 (s, 1H), 7.17 (d, $J$ = 7.8 Hz, 1H), 6.97 (ddd, $J$ = 8.1, 7.3, 1.5 Hz, 1H), 6.79 (dd, $J$ = 8.0, 1.5 Hz, 1H), 6.61 (td, $J$ = 7.5, 1.4 Hz, 1H), 6.07 (dd, $J$ = 2.7, 2.0 Hz, 1H), 4.99 (s, 2H), 4.88 (s, 2H), 3.57 (s, 3H). $^{13}$C NMR (126 MHz, DMSO) δ 165.78, 154.22, 146.03, 142.85, 134.57, 130.20, 129.32, 128.85, 126.78, 126.35, 126.21, 123.87, 123.78, 123.31, 116.38, 116.26, 115.18, 114.41, 111.71, 102.50, 35.55. MS (ESI): m/z calc. for $[C_{21}H_{19}N_5O_2+H^+]^+$ = 374.15, found = 374.10. HRMS (MALDI): m/z calc. for $[C_{21}H_{19}N_5O_2+Na^+]^+$ = 396.1431, found = 396.1441

***N*-(2-Aminophenyl)-3-methoxy-4-(6-methyl-7-oxo-6,7-dihydro-1*H*-pyrrolo[2,3-*c*]pyridin-4-yl)benzamide NB470 (31c)**

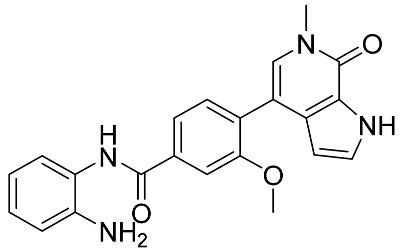


The synthesis was performed according to "general procedure for amide coupling". **30c** (400 mg, 1.34 mmol, 1.0 eq) and benzene-1,2-diamine (189 mg, 1.74 mmol, 1.3 eq) were used. The crude product was triturated with methanol and filtered to provide the title compound as a colorless solid (388 mg, 74%). $^1$H NMR (500 MHz, DMSO-$d_6$) δ 12.01 (s, 1H), 9.76 (s, 1H), 7.71 (d, $J$ = 1.7 Hz, 1H), 7.67 (dd, $J$ = 7.8, 1.7 Hz, 1H), 7.45 (d, $J$ = 7.8 Hz, 1H), 7.28 (t, $J$ = 2.8 Hz, 1H), 7.26 (s, 1H), 7.19 (dd, $J$ = 7.9, 1.6 Hz, 1H), 7.00 (td, $J$ = 7.6, 1.5 Hz, 1H), 6.81 (dd, $J$ = 8.0, 1.5 Hz, 1H), 6.63 (td, $J$ = 7.5, 1.4 Hz, 1H), 6.07 (t, $J$ = 2.4 Hz, 1H), 4.93 (s, 2H), 3.83 (s, 3H), 3.57 (s, 3H). $^{13}$C NMR (126 MHz, DMSO) δ 164.91, 156.54, 154.14, 143.29, 134.76, 130.36, 129.75, 129.10, 128.77, 126.84, 126.63, 126.58, 123.30, 122.89, 120.06, 116.29, 116.16, 111.30, 110.77, 102.87, 55.51, 35.57. MS (ESI): m/z calc. for $[C_{22}H_{20}N_4O_3+H^+]^+$ = 389.15, found = 389.10. HRMS (MALDI): m/z calc. for $[C_{22}H_{20}N_4O_3+Na^+]^+$ = 411.1428, found = 411.1434

***N*-(2-Aminophenyl)-5-(6-methyl-7-oxo-6,7-dihydro-1*H*-pyrrolo[2,3-*c*]pyridin-4-yl)picolin-amide NB500 (31d)**

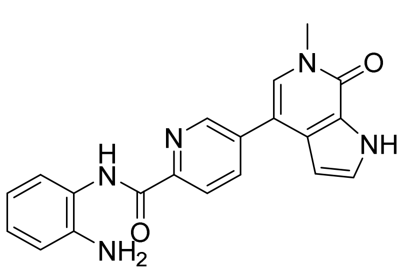


The synthesis was performed according to "general procedure for amide coupling". **30d** (170 mg, 631 μmol, 1.0 eq) and benzene-1,2-diamine (82 mg, 0.76 mmol, 1.2 eq) were used. The crude product was triturated with methanol and filtered to provide the title compound as a pale yellow solid (80 mg, 35%). $^1$H NMR (500 MHz, DMSO-$d_6$) δ 12.28 (s, 1H), 10.08 (s, 1H), 8.95 (dd, *J* = 2.2, 0.9 Hz, 1H), 8.26 (dd, *J* = 8.1, 2.2 Hz, 1H), 8.22 (dd, *J* = 8.1, 0.9 Hz, 1H), 7.67 (s, 1H), 7.54 (dd, *J* = 8.0, 1.5 Hz, 1H), 7.42 (t, *J* = 2.8 Hz, 1H), 6.97 (td, *J* = 7.6, 1.5 Hz, 1H), 6.85 (dd, *J* = 8.0, 1.5 Hz, 1H), 6.67 (td, *J* = 7.6, 1.5 Hz, 1H), 6.53 (dd, *J* = 2.8, 2.0 Hz, 1H), 4.92 (s, 2H), 3.63 (s, 3H). $^{13}$C NMR (126 MHz, DMSO) δ 162.07, 148.02, 146.48, 141.67, 135.86, 135.76, 129.57, 127.69, 127.56, 125.82, 124.39, 124.13, 123.39, 122.33, 117.06, 116.80, 110.49, 101.84, 35.72. MS (ESI): m/z calc. for $[C_{20}H_{17}N_5O_2+H^+]^+$ = 360.14, found = 360.05. HRMS (MALDI): m/z calc. for $[C_{20}H_{17}N_5O_2+Na^+]^+$ = 382.1275, found = 382.1286

***N*-(2-Aminophenyl)-6-(6-methyl-7-oxo-6,7-dihydro-1*H*-pyrrolo[2,3-*c*]pyridin-4-yl)pyridazine-3-carboxamide NB501 (31e)**

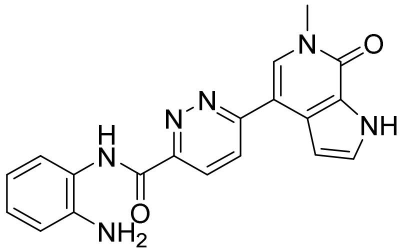


The synthesis was performed according to "general procedure for amide coupling". **30e** (130 mg, 481 μmol, 1.0 eq) and benzene-1,2-diamine (62 mg, 0.58 mmol, 1.2 eq) were used. The crude product was triturated with methanol and filtered to provide the title compound as a pale yellow solid (100 mg, 58%). $^1$H NMR (500 MHz, DMSO-$d_6$) δ 12.26 (s, 1H), 10.38 (s, 1H), 8.36 (d, *J* = 9.0 Hz, 1H), 8.28 (d, *J* = 8.9 Hz, 1H), 8.24 (s, 1H), 7.45 (dd, *J* = 7.9, 1.5 Hz, 1H), 7.43 (t, *J* = 2.8 Hz, 1H), 7.09 (t, *J* = 2.4 Hz, 1H), 7.00 (td, *J* = 7.6, 1.5 Hz, 1H), 6.84 (dd, *J* = 8.1, 1.5 Hz, 1H), 6.66 (td, *J* = 7.5, 1.5 Hz, 1H), 5.01 (s, 2H), 3.67 (s, 3H). $^{13}$C NMR (126 MHz, DMSO) δ 161.32, 159.32, 154.39, 150.75, 142.43, 132.49, 127.45, 126.99, 126.35, 125.87, 125.47, 124.69, 123.41, 123.37, 116.72, 116.56, 109.96, 104.53, 36.04. MS (ESI): m/z calc. for $[C_{19}H_{16}N_6O_2+H^+]^+$ = 361.13, found = 361.05. HRMS (MALDI): m/z calc. for $[C_{19}H_{16}N_6O_2+Na^+]^+$ = 383.1227, found = 383.1240

***N*-(2-Aminophenyl)-6-(6-methyl-7-oxo-6,7-dihydro-1*H*-pyrrolo[2,3-*c*]pyridin-4-yl)nicotin-amide NB502 (31f)**

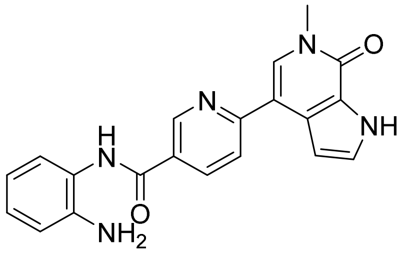


The synthesis was performed according to "general procedure for amide coupling". **30f** (170 mg, 631 µmol, 1.0 eq) and benzene-1,2-diamine (82 mg, 0.76 mmol, 1.2 eq) were used. The crude product was triturated with methanol and filtered to provide the title compound as a beige solid (126 mg, 56%). $^1$H NMR (500 MHz, DMSO-$d_6$) δ 12.18 (s, 1H), 9.81 (s, 1H), 9.21 (d, $J$ = 2.3 Hz, 1H), 8.38 (dd, $J$ = 8.3, 2.4 Hz, 1H), 8.10 (s, 1H), 7.98 (d, $J$ = 8.3 Hz, 1H), 7.40 (t, $J$ = 2.8 Hz, 1H), 7.20 (dd, $J$ = 7.9, 1.4 Hz, 1H), 7.03 – 6.94 (m, 2H), 6.80 (dd, $J$ = 8.1, 1.4 Hz, 1H), 6.61 (td, $J$ = 7.5, 1.4 Hz, 1H), 4.99 (s, 2H), 3.66 (s, 3H). $^{13}$C NMR (126 MHz, DMSO) δ 163.83, 157.53, 154.35, 148.74, 143.38, 136.14, 131.19, 127.41, 127.22, 126.92, 126.73, 123.45, 122.81, 119.47, 116.12, 115.97, 112.76, 104.00, 35.90. MS (ESI): m/z calc. for $[C_{20}H_{17}N_5O_2+H^+]^+$ = 360.14, found = 360.10. HRMS (MALDI): m/z calc. for $[C_{20}H_{17}N_5O_2+Na^+]^+$ = 382.1275, found = 382.1285

***tert*-Butyl (4-bromo-2-nitrophenyl)carbamate (33)**

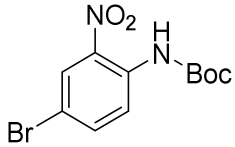


4-Bromo-2-nitroaniline (**32**, 7.00 g, 32.3 mmol, 1.0 eq) was dissolved in anh. THF (100 mL), cooled to -10 °C and sodium hydride (60%, 1.70 g, 71.0 mmol, 2.2 eq) was carefully added. After 10 min, a solution of di-*tert*-butyl dicarbonate (7.74 g, 35.5 mmol, 1.1 eq) in THF (60 mL) was added dropwise, and the mixture was stirred for 4 h. The reaction was quenched with ice, and the mixture was partitioned between water and ethyl acetate. The ethyl acetate layer was washed with brine, dried over $MgSO_4$, filtered, and volatiles were removed under reduced pressure. Purification by column chromatography (silica, hexane/ethyl acetate) provided the product as a pale-yellow solid (9.45 g, 92%). $^1$H-NMR (300 MHz, DMSO-d6) δ 9.66 (s, 1H), 8.12 (d, $J$ = 2.3 Hz, 1H), 7.86 (dd, $J$ = 8.8, $J$ = 2.4 Hz, 1H), 7.60 (d, $J$ = 8.8 Hz, 1H), 1.44 (s, 9H). MS (ESI): m/z calc. for $[C_{11}H_{13}BrN_2O_4+Na^+]^+$ = 339.00, found = 338.92

***tert*-Butyl (3-nitro-[1,1'-biphenyl]-4-yl)carbamate (34a)**

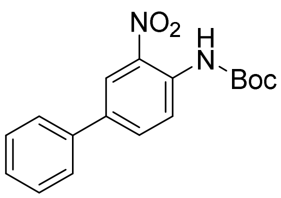


The synthesis was performed according to "general procedure for Suzuki coupling A". *tert*-Butyl (4-bromo-2-nitrophenyl)carbamate (**33**, 2.00 g, 6.31 mmol, 1.0 eq) and phenylboronic acid (923 mg, 7.57 mmol, 1.2 eq) were used. The product was isolated as an orange solid

(1.65 g, 83%). $^1$H-NMR (300 MHz, DMSO-d$_6$) δ 9.64 (s, 1H), 8.18 (d, *J* = 2.2 Hz, 1H), 7.99 (dd, *J* = 8.6, *J* = 2.2 Hz, 1H), 7.81 – 7.68 (m, 3H), 7.54 – 7.36 (m, 3H), 1.46 (s, 9H). MS (ESI): m/z calc. for $[C_{17}H_{18}N_2O_4+Na^+]^+$ = 337.12, found = 337.11

**_tert_-Butyl (4-(furan-2-yl)-2-nitrophenyl)carbamate (34b)**

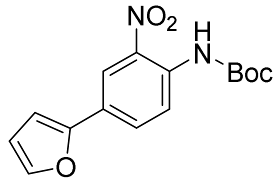


The synthesis was performed according to "general procedure for Suzuki coupling A". *tert*-Butyl (4-bromo-2-nitrophenyl)carbamate (**33**, 1.94 g, 6.12 mmol, 1.0 eq) and furan-2-ylboronic acid (821 mg, 7.34 mmol, 1.2 eq) were used. The product was isolated as an orange solid (1.27 g, 68%). $^1$H-NMR (300 MHz, DMSO-d$_6$) δ 9.63 (s, 1H), 8.19 (d, *J* = 2.1 Hz, 1H), 7.97 (dd, *J* = 8.6, *J* = 2.1 Hz, 1H), 7.80 (dd, *J* = 1.8, *J* = 0.6 Hz, 1H), 7.73 (d, *J* = 8.6 Hz, 1H), 7.10 (dd, *J* = 3.4, *J* = 0.6 Hz, 1H), 6.63 (dd, *J* = 3.4, *J* = 1.8 Hz, 1H), 1.45 (s, 9H). MS (ESI): m/z calc. for $[C_{15}H_{16}N_2O_5+Na^+]^+$ = 327.10, found = 327.09

**_tert_-Butyl (2-nitro-4-(thiophen-2-yl)phenyl)carbamate (34c)**

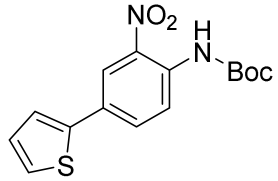


The synthesis was performed according to "general procedure for Suzuki coupling A". *tert*-Butyl (**33**, 4-bromo-2-nitrophenyl)carbamate (1.90 g, 5.99 mmol, 1.0 eq) and thiophen-2-ylboronic acid (920 mg, 7.19 mmol, 1.2 eq) were used. The product was isolated as an orange solid (1.13 g, 59%). $^1$H-NMR (300 MHz, DMSO-d$_6$) δ 9.63 (s, 1H, g), 8.15 (d, *J* = 2.2 Hz, 1H), 7.94 (dd, *J* = 8.6, *J* = 2.3 Hz, 1H), 7.70 (d, *J* = 8.7 Hz, 1H), 7.64 – 7.60 (m, 2H), 7.17 (dd, *J* = 5.0, *J* = 3.7 Hz, 1H), 1.45 (s, 9H). MS (ESI): m/z calc. for $[C_{15}H_{16}N_2O_4S+Na^+]^+$ = 343.07, found = 343.09

**_tert_-Butyl (3-amino-[1,1'-biphenyl]-4-yl)carbamate (35a)**

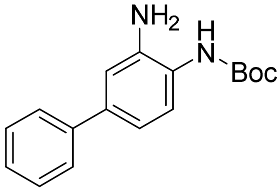


The synthesis was performed according to "general procedure for reduction of nitro groups". *tert*-Butyl (3-nitro-[1,1'-biphenyl]-4-yl)carbamate (**34a**, 1.47 g, 4.68 mmol) was used. The product was isolated as a colorless solid (1.288 g, 97%). $^1$H-NMR (300 MHz, DMSO-d6) δ 8.34 (s, 1H), 7.56 – 7.51 (m, 2H), 7.46 – 7.39 (m, 2H), 7.34 – 7.28 (m, 2H), 6.99 (d, *J* = 2.1 Hz, 1H), 6.83 (dd, *J* = 8.2, *J* = 2.2 Hz, 1H), 4.95 (s, 2H), 1.47 (s, 9H). MS (ESI): m/z calc. for $[C_{17}H_{20}N_2O_2+Na^+]^+$ = 307.14, found = 307.05

**tert-Butyl (2-amino-4-(furan-2-yl)phenyl)carbamate (35b)**

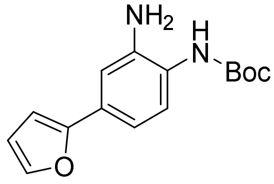


The synthesis was performed according to “general procedure for reduction of nitro groups”. *tert*-Butyl (4-(furan-2-yl)-2-nitrophenyl)carbamate (**34b**, 1.23 g, 4.04 mmol) was used. The product was isolated as a colorless solid (0.989 g, 89%). $^{1}$H-NMR (300 MHz, DMSO-d6) δ 8.32 (s, 1H), 7.66 (dd, *J* = 1.8, *J* = 0.7 Hz, 1H), 7.28 (d, *J* = 8.3 Hz, 1H), 7.04 (d, *J* = 2.0 Hz, 1H), 6.88 (dd, *J* = 8.2, *J* = 2.0 Hz, 1H), 6.68 (dd, *J* = 3.3, *J* = 0.7 Hz, 1H), 6.53 (dd, *J* = 3.4, *J* = 1.8 Hz, 1H), 4.98 (s, 2H), 1.46 (s, 9H). MS (ESI): m/z calc. for $[C_{17}H_{20}N_2O_2+Na^+]^+$ = 297.12, found = 297.11

**tert-Butyl (2-amino-4-(thiophen-2-yl)phenyl)carbamate (35c)**

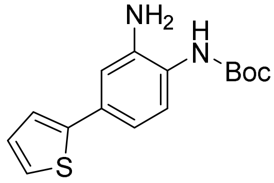


The synthesis was performed according to “general procedure for reduction of nitro groups”. *tert*-Butyl (2-nitro-4-(thiophen-2-yl)phenyl)carbamate (**34c**, 1.10 g, 3.43 mmol) was used. The product was isolated as a colorless solid (0.928 g, 93%). $^{1}$H-NMR (300 MHz, DMSO-d6) δ 8.33 (s, 1H), 7.44 (dd, *J* = 5.1, *J* = 1.1 Hz, 1H), 7.29 (dd, *J* = 3.6, *J* = 1.2 Hz, 1H), 7.27 (d, *J* = 8.4 Hz, 1H), 7.08 (dd, *J* = 5.1, *J* = 3.6 Hz, 1H), 6.98 (d, *J* = 2.1 Hz, 1H), 6.84 (dd, *J* = 8.2, *J* = 2.0 Hz, 1H), 5.00 (s, 2H), 1.47 (s, 9H). MS (ESI): m/z calc. for $[C_{15}H_{18}N_2O_2S+Na^+]^+$ = 313.10, found = 313.06

**tert-Butyl (3-(4-(6-methyl-7-oxo-6,7-dihydro-1*H*-pyrrolo[2,3-*c*]pyridin-4-yl)benzamido)-[1,1'-biphenyl]-4-yl)carbamate (36)**

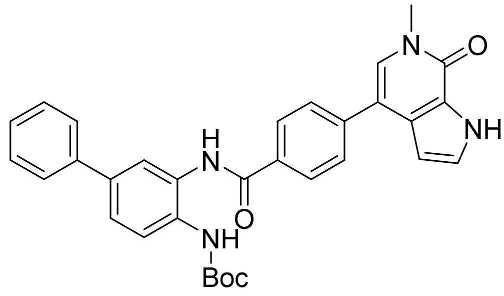


The synthesis was performed according to “general procedure for amide coupling”. **30a** (160 mg, 596 µmol, 1.0 eq) and **35a** (170 mg, 596 µmol, 1.0 eq) were used. The obtained beige solid (210 mg, 66%) was used in the next step without further characterization. MS (ESI): m/z calc. for $[C_{32}H_{30}N_4O_4+H^+]^+$ = 535.23, found = 535.20

***tert*-Butyl (3-(5-(6-methyl-7-oxo-6,7-dihydro-1*H*-pyrrolo[2,3-*c*]pyridin-4-yl)picolinamido)-[1,1'-biphenyl]-4-yl)carbamate (37a)**

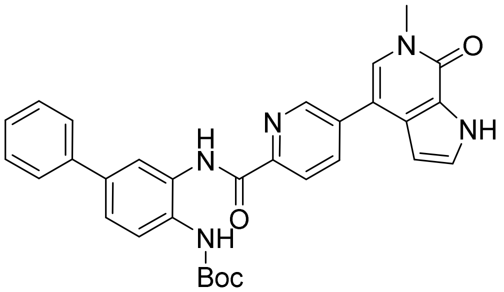

The synthesis was performed according to "general procedure for amide coupling". **30d** (100 mg, 371 µmol, 1.0 eq) and **35a** (116 mg, 409 µmol, 1.1 eq) were used. The product was obtained as a beige solid (198 mg, quant.). $^{1}$H NMR (500 MHz, DMSO-$d_6$) δ 12.29 (s, 1H), 10.58 (s, 1H), 9.24 (s, 1H), 8.88 (dd, *J* = 2.2, 0.9 Hz, 1H), 8.33 (d, *J* = 2.1 Hz, 1H), 8.29 (dd, *J* = 8.2, 2.2 Hz, 1H), 8.26 (dd, *J* = 8.1, 0.9 Hz, 1H), 7.71 – 7.63 (m, 3H), 7.51 – 7.46 (m, 3H), 7.43 (t, *J* = 2.8 Hz, 1H), 7.41 – 7.36 (m, 2H), 6.50 (dd, *J* = 2.9, 2.0 Hz, 1H), 3.63 (s, 3H), 1.52 (s, 9H). $^{13}$C NMR (126 MHz, DMSO) δ 161.98, 154.14, 153.93, 147.41, 146.41, 139.54, 137.37, 136.23, 136.11, 132.13, 129.78, 129.17, 129.00, 127.63, 127.48, 126.54, 123.37, 123.10, 122.49, 110.34, 101.69, 79.85, 35.73, 28.06. MS (ESI): m/z calc. for $[C_{31}H_{29}N_5O_4+H^+]^+$ = 536.22, found = 536.20

***tert*-Butyl (4-(furan-2-yl)-2-(5-(6-methyl-7-oxo-6,7-dihydro-1*H*-pyrrolo[2,3-*c*]pyridin-4-yl)picolinamido)phenyl)carbamate (37b)**

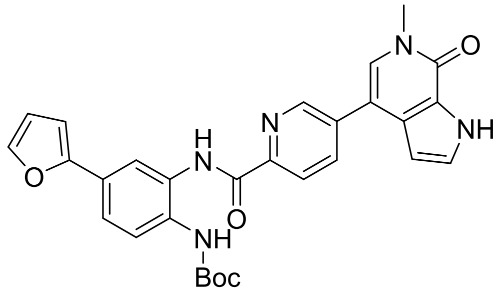

The synthesis was performed according to "general procedure for amide coupling". **30d** (100 mg, 371 µmol, 1.0 eq) and **35b** (112 mg, 409 µmol, 1.1 eq) were used. The crude product (195 mg, quant.) was used in the next step without further characterization. MS (ESI): m/z calc. for $[C_{29}H_{27}N_5O_5+H^+]^+$ = 526.20, found = 526.15

***tert*-Butyl (2-(5-(6-methyl-7-oxo-6,7-dihydro-1*H*-pyrrolo[2,3-*c*]pyridin-4-yl)picolinamido)-4-(thiophen-2-yl)phenyl)carbamate (37c)**

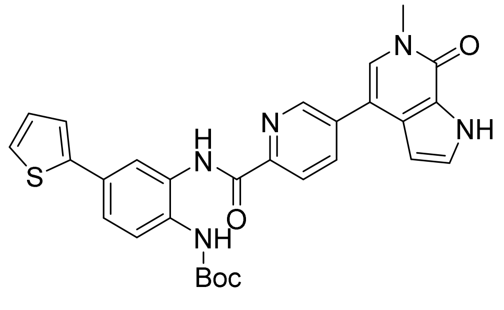

The synthesis was performed according to "general procedure for amide coupling". **30d** (50 mg, 186 µmol, 1.0 eq) and **35c** (54 mg, 0.19 mmol, 1.0 eq) were used. The crude product (100 mg, quant.) was used in the next step without further characterization. MS (ESI): m/z calc. for $[C_{29}H_{27}N_5O_4S+H^+]^+$ = 542.18, found = 542.15

**_N_-(4-Amino-[1,1'-biphenyl]-3-yl)-4-(6-methyl-7-oxo-6,7-dihydro-1_H_-pyrrolo[2,3-_c_]pyridin-4-yl)benzamide NB503 (38)**

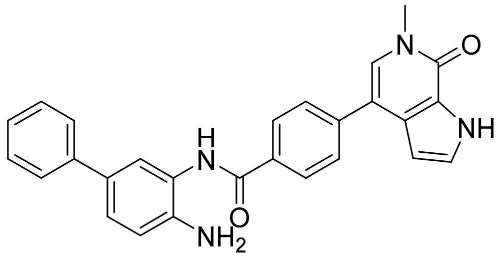

The synthesis was performed according to "general procedure for *N-Boc* deprotection". **36** (210 mg, 393 µmol) was dissolved in DCM/TFA (10 mL, 3/1). The crude product was triturated with methanol and filtered to provide the title compound as a beige solid (128 mg, 75%). $^1$H NMR (500 MHz, DMSO-$d_6$) δ 12.22 (s, 1H), 10.29 (s, 1H), 8.18 (d, $J$ = 8.3 Hz, 2H), 7.79 (d, $J$ = 8.3 Hz, 2H), 7.74 (d, $J$ = 2.2 Hz, 1H), 7.64 (d, $J$ = 7.1 Hz, 2H), 7.56 (s, 1H), 7.54 (dd, $J$ = 8.4, 2.3 Hz, 1H), 7.45 (t, $J$ = 7.7 Hz, 2H), 7.41 (t, $J$ = 2.8 Hz, 1H), 7.34 (t, $J$ = 7.4 Hz, 1H), 7.27 (d, $J$ = 8.3 Hz, 1H), 6.70 (br s, 2H), 6.52 (t, $J$ = 2.4 Hz, 1H), 3.63 (s, 3H). $^{13}$C NMR (126 MHz, DMSO) δ 165.54, 154.17, 140.75, 139.31, 131.97, 129.00, 128.82, 128.60, 128.08, 127.36, 127.17, 126.96, 126.17, 124.97, 124.85, 123.44, 113.77, 102.14, 35.67. MS (ESI): m/z calc. for $[C_{27}H_{22}N_4O_2+H^+]^+$ = 435.17, found = 435.10. HRMS (MALDI): m/z calc. for $[C_{27}H_{22}N_4O_2+H^+]^+$ = 457.1635, found = 457.1639

**_N_-(4-Amino-[1,1'-biphenyl]-3-yl)-5-(6-methyl-7-oxo-6,7-dihydro-1_H_-pyrrolo[2,3-_c_]pyridin-4-yl)picolinamide NB512 (39a)**

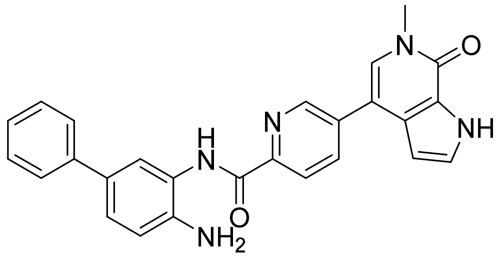

The synthesis was performed according to "general procedure for *N-Boc* deprotection". **37a** (198 mg, 370 µmol) was dissolved in DCM/TFA (10 mL, 3/1). The crude product was triturated with methanol and filtered to provide the title compound as a pale yellow solid (100 mg, 62%). $^1$H NMR (500 MHz, DMSO-$d_6$) δ 12.28 (s, 1H), 10.17 (s, 1H), 8.97 (dd, $J$ = 2.2, 0.9 Hz, 1H), 8.28 (dd, $J$ = 8.1, 2.2 Hz, 1H), 8.24 (dd, $J$ = 8.2, 0.8 Hz, 1H), 7.88 (d, $J$ = 2.1 Hz, 1H), 7.68 (s, 1H), 7.60 – 7.55 (m, 2H), 7.44 – 7.38 (m, 3H), 7.32 (dd, $J$ = 8.3, 2.2 Hz, 1H), 7.28 – 7.24 (m, 1H), 6.93 (d, $J$ = 8.3 Hz, 1H), 6.54 (dd, $J$ = 2.8, 2.0 Hz, 1H), 5.11 (s, 2H), 3.63 (s, 3H). $^{13}$C NMR (126 MHz, DMSO) δ 162.29, 154.15, 147.99, 146.50, 141.51, 140.31, 135.89, 135.82, 129.60, 128.90, 128.80, 127.70, 127.58, 126.12, 125.65, 124.27, 124.16, 123.39, 122.80, 122.40, 117.11, 110.50, 101.84, 35.73. MS (ESI): m/z calc. for $[C_{26}H_{21}N_5O_2+H^+]^+$ = 436.17, found = 436.15. HRMS (MALDI): m/z calc. for $[C_{26}H_{21}N_5O_2+Na^+]^+$ = 458.1588, found = 458.1582

***N*-(2-Amino-5-(furan-2-yl)phenyl)-5-(6-methyl-7-oxo-6,7-dihydro-1*H*-pyrrolo[2,3-*c*]pyridin-4-yl)picolinamide NB513 (39b)**

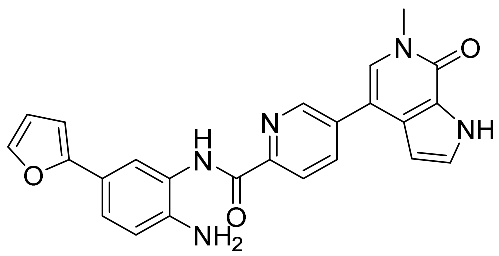


The synthesis was performed according to "general procedure for *N-Boc* deprotection". **37b** (195 mg, 371 µmol) was dissolved in DCM/TFA (10 mL, 3/1). The crude product was triturated with methanol and filtered to provide the title compound as a beige solid (87 mg, 55%). $^1$H NMR (500 MHz, DMSO-$d_6$) δ 12.28 (s, 1H), 10.12 (s, 1H), 8.96 (d, $J$ = 2.2 Hz, 1H), 8.30 – 8.20 (m, 2H), 7.89 (d, $J$ = 2.1 Hz, 1H), 7.72 – 7.59 (m, 2H), 7.42 (t, $J$ = 2.8 Hz, 1H), 7.33 (dd, $J$ = 8.3, 2.1 Hz, 1H), 6.88 (d, $J$ = 8.4 Hz, 1H), 6.63 (d, $J$ = 3.3 Hz, 1H), 6.54 (t, $J$ = 2.4 Hz, 1H), 6.52 (dd, $J$ = 3.3, 1.8 Hz, 1H), 5.26 – 5.01 (m, 2H), 3.63 (s, 3H). $^{13}$C NMR (126 MHz, DMSO) δ 162.35, 154.15, 153.80, 147.94, 146.49, 141.62, 141.34, 135.88, 135.83, 129.61, 127.69, 127.58, 123.98, 123.39, 122.40, 121.70, 120.17, 119.87, 116.80, 111.80, 110.49, 102.66, 101.84, 35.73. MS (ESI): m/z calc. for $[C_{24}H_{19}N_5O_3+H^+]^+$ = 426.15, found = 426.10. HRMS (MALDI): m/z calc. for $[C_{24}H_{19}N_5O_3+Na^+]^+$ = 448.1380, found = 448.1385

***N*-(2-Amino-5-(thiophen-2-yl)phenyl)-5-(6-methyl-7-oxo-6,7-dihydro-1*H*-pyrrolo[2,3-*c*]pyridin-4-yl)picolinamide NB514 (39c)**

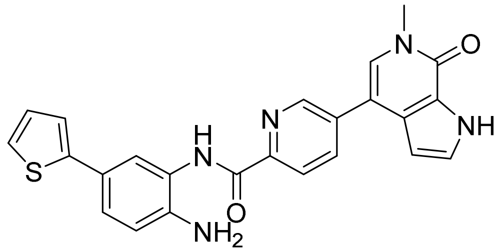


The synthesis was performed according to "general procedure for *N-Boc* deprotection". **37c** (100 mg, 185 µmol) was dissolved in DCM/TFA (8 mL, 3/1). The crude product was triturated with methanol and filtered to provide the title compound as a beige solid (34 mg, 41%). $^1$H NMR (500 MHz, DMSO-$d_6$) δ 12.29 (s, 1H), 10.19 (s, 1H), 8.98 – 8.96 (m, 1H), 8.29 (dd, $J$ = 8.2, 2.2 Hz, 1H), 8.27 – 8.22 (m, 1H), 7.85 (d, $J$ = 2.2 Hz, 1H), 7.69 (s, 1H), 7.43 (t, $J$ = 2.8 Hz, 1H), 7.39 (dd, $J$ = 5.1, 1.2 Hz, 1H), 7.32 (dd, $J$ = 8.3, 2.2 Hz, 1H), 7.28 (dd, $J$ = 3.6, 1.2 Hz, 1H), 7.08 (dd, $J$ = 5.1, 3.6 Hz, 1H), 6.90 (d, $J$ = 8.3 Hz, 1H), 6.56 – 6.53 (m, 1H), 5.08 (br s, 2H), 3.64 (s, 3H). $^{13}$C NMR (126 MHz, DMSO) δ 162.40, 154.15, 147.88, 146.50, 144.18, 135.88, 135.87, 129.62, 128.24, 127.69, 127.59, 124.39, 123.54, 123.39, 123.37, 123.33, 123.31, 122.45, 121.98, 121.35, 117.24, 110.48, 101.84, 35.73. MS (ESI): m/z calc. for $[C_{24}H_{19}N_5O_2S+H^+]^+$ = 442.13, found = 442.10. HRMS (MALDI): m/z calc. for $[C_{24}H_{19}N_5O_2S+Na^+]^+$ = 464.1152, found = 464.1151

***N*-(2-Amino-4-fluorophenyl)-5-(6-methyl-7-oxo-6,7-dihydro-1*H*-pyrrolo[2,3-*c*]pyridin-4-yl)picolinamide NB507 (40)**

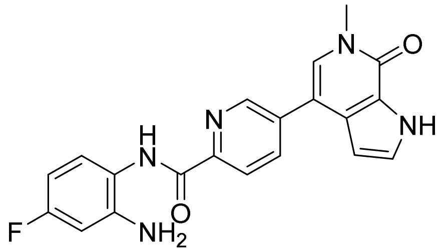


The synthesis was performed according to "general procedure for amide coupling". **30d** (90 mg, 0.33 mmol, 1.0 eq) and 4-fluorobenzene-1,2-diamine (51 mg, 0.40 mmol, 1.2 eq) were used. The crude product was triturated with methanol and filtered to provide the title compound as a beige solid (48 mg, 38%). $^{1}$H NMR (500 MHz, DMSO-*d*$_6$) δ 12.27 (s, 1H), 9.99 (s, 1H), 8.94 (dd, *J* = 2.2, 0.8 Hz, 1H), 8.25 (dd, *J* = 8.2, 2.3 Hz, 1H), 8.20 (dd, *J* = 8.2, 0.8 Hz, 1H), 7.66 (s, 1H), 7.42 (t, *J* = 2.8 Hz, 1H), 7.37 (dd, *J* = 8.7, 6.3 Hz, 1H), 6.60 (dd, *J* = 11.1, 2.9 Hz, 1H), 6.52 (dd, *J* = 2.8, 2.0 Hz, 1H), 6.42 (td, *J* = 8.6, 2.9 Hz, 1H), 5.25 (s, 2H), 3.62 (s, 3H). $^{19}$F NMR (471 MHz, DMSO-*d*$_6$) δ -116.72 (ddd, *J* = 11.7, 8.9, 6.6 Hz). $^{13}$C NMR (126 MHz, DMSO) δ 162.51, 154.14, 148.00, 146.45, 144.60, 144.50, 135.81, 135.75, 129.55, 127.70, 127.57, 126.95, 126.87, 123.38, 122.40, 119.70, 119.68, 110.50, 102.63, 102.45, 102.17, 101.97, 101.82, 35.72. MS (ESI): m/z calc. for $[C_{20}H_{16}FN_5O_2+H^+]^+$ = 378.13, found = 360.10. HRMS (MALDI): m/z calc. for $[C_{20}H_{16}FN_5O_2+Na^+]^+$ = 400.1180, found = 400.1183

**Supporting Table S1.** ITC thermodynamic data of NBx binding to BRD4 BD1

| | Compound | |
|---|---|---|
| | NB462 | NB500 |
| $K_d$ (M) | 1.52E-07 ± 7.29E-08 | 4.58E-08 ± 3.51E-08 |
| n | 0.996 ± 0.029 | 1.003 ± 0.022 |
| ΔH (kcal/mol) | -10.36 ± 0.501 | -7.844 ± 0.360 |
| ΔS (cal/mol·K) | -4.759 | 6.358 |
| ΔG (kcal/mol) | -8.99 | -9.676 |
| -TΔS (kcal/mol) | 1.371 | -1.832 |
| $K_a$ ($M^{-1}$) | 6.58E+06 | 2.18E+07 |
| Confidence Level (%) | 95 | 95 |

**Supporting Table S2.** Bromodomain panel selectivity data

| Protein | Mean $\Delta T_m$ (K) | |
|---|---|---|
| | NB462 | NB503 |
| ATAD2A | 0.4 ± 0.2 | -1.3 ± 0.1 |
| BAZ2B | 0.5 ± 0.0 | 1.4 ± 0.1 |
| BRD1 | 0.2 ± 0.1 | 1.1 ± 0.1 |
| BRD2 (1) | 3.1 ± 0.5 | 3.2 ± 0.5 |
| BRD2 (2) | 5.3 ± 0.1 | 3.6 ± 0.4 |
| BRD3 (1) | 3.8 ± 0.3 | 3.8 ± 0.4 |
| BRD3 (2) | 6.1 ± 0.0 | 4.6 ± 0.7 |
| BRD4 (1) | 3.6 ± 0.1 | 5.5 ± 0.1 |
| BRD4 (2) | 4.6 ± 0.1 | 5.3 ± 0.2 |
| BRD7 | 5.6 ± 0.3 | 1.1 ± 0.2 |
| BRD9 | 5.2 ± 1.2 | 1.0 ± 0.8 |
| BRDT(1) | 3.0 ± 0.2 | 3.3 ± 0.1 |
| BRDT(2) * | n.d. | n.d. |
| BRPF1B | 0.8 ± 0.1 | 0.8 ± 0.3 |
| BRPF3 | -0.5 ± 0.1 | 1.2 ± 0.1 |
| CREBBPA | 1.8 ± 0.3 | 1.4 ± 0.3 |
| EP300A | 2.0 ± 0.3 | 1.0 ± 0.3 |
| PB1A(3) | 0.5 ± 0.2 | 0.0 ± 0.0 |
| PB1A(4) | 0.1 ± 0.1 | 1.0 ± 0.6 |
| PB1A(5) | 0.0 ± 0.1 | 1.1 ± 0.2 |
| PB1A(6) | 0.3 ± 0.1 | 1.3 ± 0.1 |
| PCAFA | 0.1 ± 0.4 | 1.9 ± 0.2 |
| SMARCA2A | -0.1 ± 0.1 | 1.2 ± 0.5 |
| SP100 | -0.2 ± 0.1 | 0.4 ± 0.1 |
| TAF1 (1) | 0.4 ± 0.1 | 1.0 ± 0.2 |
| TAF1 (2) | 1.3 ± 0.0 | 1.7 ± 0.1 |
| TAF1LA (1) | 0.3 ± 0.1 | 0.6 ± 0.3 |
| TAF1LA (2) | 1.7 ± 0.1 | 1.2 ± 0.0 |
| TRIM24** | 0.6 ± 0.1 | 0.7 ± 0.5 |
| TRIM28** | 0.6 ± 0.4 | 0.0 ± 0.2 |
| TRIM33B** | 0.4 ± 0.1 | 0.5 ± 0.0 |
| WDR9A(1) | -0.3 ± 0.2 | 0.0 ± 0.0 |

* Boltzmann fitting failed

** Construct containing tandem PHD-BD

**Supporting Table S3.** Cell viability upon treatment with dual BET/HDAC inhibitors for 3 days in PaTu8988t and HCC2429 cell lines.

| **Compound** | **Cell viability [$IC_{50}$; µM]** | |
|---|---|---|
| | **PaTu8988t** | **HCC2429** |
| NB462 | 4.4 | 0.20 |
| NB500 | 5.5 | 0.19 |
| NB501 | 18.6 | 0.48 |
| NB502 | 19.0 | 0.61 |
| NB503 | 5.2 | 0.58 |
| NB507 | 8.1 | 0.32 |
| NB512 | 6.3 | 0.55 |
| NB513 | 8.6 | 0.49 |
| NB514 | 4.9 | 0.42 |

**Supporting Table S4.** Crystallization conditions for BRD4 BD1-inhibitor complexes

| Compound | Temperature | Reservoir buffer | Drop volume ratio (protein:reservoir solution) |
|---|---|---|---|
| NB161 | 4 °C | 25% PEG 3350, 0.15 M Na nitrate, 15% ethylene glycol, 0.1 M bis-tris propane pH 7.6 | 1:1 |
| NB390 | 4 °C | 25% PEG 3350, 0.2 M Na formate, 15% ethylene glycol, 0.1 M bis-tris propane pH 7.9 | 1:1 |
| NB437 | 4 °C | 25% PEG 3350, 0.2 M Na nitrate, 15% ethylene glycol, 0.1 M bis-tris propane pH 7.9 | 1:2 |
| NB462 | 4 °C | 24% PEG 3350, 0.15 M Na formate, 15% ethylene glycol, 0.1 M bis-tris propane pH 7.6 | 1:2 |
| NB500 | 4 °C | 25% PEG 3350, 0.1 M Na nitrate, 15% ethylene glycol, 0.1 M bis-tris propane pH 7.3 | 1:1 |
| NB503 | 4 °C | 25% PEG 3350, 0.1 M Na malonate pH 7, 15% ethylene glycol, 0.1 M bis-tris propane pH 7.3 | 1:2 |
| NB512 | 4 °C | 24% PEG 3350, 0.1 M Na/K Tartrate, 10% ethylene glycol, 0.1 M bis-tris propane pH 7.3 | 1:2 |

**Supporting Table S5.** X-ray data collection and refinement statistics

| Compound | NB161 | NB390 | NB437 | NB462 | NB500 | NB503 | NB512 |
|---|---|---|---|---|---|---|---|
| *Data Collection* | | | | | | | |
| Space Group | $P2_12_12_1$ | $P2_12_12_1$ | $P2_12_12_1$ | $P2_1$ | $P2_12_12_1$ | $P2_12_12_1$ | $P2_12_12_1$ |
| a (Å) | 42.24 | 42.24 | 42.62 | 44.69 | 44.30 | 42.32 | 42.23 |
| b (Å) | 52.69 | 52.73 | 52.50 | 51.49 | 50.96 | 52.51 | 52.40 |
| c (Å) | 56.87 | 56.81 | 56.94 | 53.11 | 53.56 | 55.05 | 54.98 |
| α (°) | 90.0 | 90.0 | 90.0 | 90.0 | 90.0 | 90.0 | 90.0 |
| β (°) | 90.0 | 90.0 | 90.0 | 91.0 | 90.0 | 90.0 | 90.0 |
| γ (°) | 90.0 | 90.0 | 90.0 | 90.0 | 90.0 | 90.0 | 90.0 |
| Molecules/AU | 1 | 1 | 1 | 2 | 1 | 1 | 1 |
| Resolution (Å)[a] | 42.2-1.30 (1.32-1.30) | 42.2-1.10 (1.12-1.10) | 42.6-1.19 (1.21-1.19) | 44.7-1.23 (1.25-1.23) | 44.3-1.42 (1.44-1.42) | 42.3-1.25 (1.27-1.25) | 42.2-1.29 (1.31-1.29) |
| Unique reflections | 31,913 | 51,898 | 40,101 | 68,247 | 22,649 | 34,045 | 31,310 |
| Completeness (%)[a] | 99.9 (100) | 99.4 (97.3) | 96.4 (93.7) | 97.6 (95.4) | 96.8 (96.7) | 98.5 (97.0) | 99.7 (99.8) |
| Multiplicity[a] | 6.5 (6.5) | 6.4 (6.2) | 6.7 (6.3) | 4.9 (4.8) | 6.6 (6.8) | 6.6 (6.2) | 8.6 (8.7) |
| $R_{merge}$ (%)[a] | 0.037 (0.873) | 0.039 (0.795) | 0.090 (0.789) | 0.072 (0.781) | 0.051 (0.903) | 0.036 (0.392) | 0.027 (1.005) |
| CC(1/2)[a] | 0.999 (0.846) | 0.999 (0.825) | 0.996 (0.815) | 0.991 (0.835) | 0.998 (0.844) | 0.999 (0.935) | 0.999 (0.876) |
| Mean $I/\sigma(I)$[a] | 22.5 (2.0) | 22.0 (2.4) | 11.1 (2.3) | 8.8 (2.3) | 16.8 (2.2) | 26.5 (4.6) | 31.8 (2.1) |
| *Refinement* | | | | | | | |
| $R_{work}$, (%)[b] | 17.9 | 15.5 | 15.1 | 16.7 | 17.0 | 15.6 | 16.3 |
| $R_{free}$, (%)[b] | 21.5 | 18.3 | 17.3 | 19.3 | 20.6 | 17.5 | 18.8 |
| No. of atoms | | | | | | | |
| Protein[c] | 1028 | 1031 | 1054 | 2097 | 1047 | 1052 | 1028 |
| Water | 124 | 150 | 155 | 303 | 86 | 135 | 121 |
| Ligands[c] | 74 | 70 | 82 | 129 | 55 | 57 | 49 |
| RMSD bonds (Å) | 0.007 | 0.006 | 0.006 | 0.006 | 0.006 | 0.005 | 0.005 |
| RMSD angles (°) | 0.91 | 0.88 | 0.99 | 0.97 | 0.94 | 0.96 | 0.83 |
| Mean $B$ (Å$^2$) | 25.4 | 20.5 | 19.8 | 20.1 | 28.5 | 18.5 | 26.8 |
| PDB entry | 8P9F | 8P9G | 8P9H | 8P9I | 8P9J | 8P9K | 8P9L |

[a]Values in parentheses are for the highest-resolution shell.
[b]$R_{work}$ and $R_{free}$ = $\sum||F_{obs}| - |F_{calc}||/\sum|F_{obs}|$, where $R_{free}$ was calculated with 5 % of the reflections chosen at random and not used in the refinement.
[c]Number includes alternative conformations.

**Supporting Table S6.** List of PCR primers used

| Gene | Forward sequence | Reverse sequence |
|---|---|---|
| *HEXIM1* | CTAGGGAACTGGGAGCTTGG | AAGGGTTAAATCCCCTGCCG |
| *p57* | AGATCAGCGCCTGAGAAGTCGT | CTCGGGGCTCTTTGGGCTCT |
| *MYC* | CAGCTGCTTAGACGCTGGATT | GTAGAAATACGGCTGCACCGA |
| *TP63* | TGGAAACCAGAGATGGGCAA | CGGGCGCTTCGTACCATC |
| *GUSB* | TGCAGGTGATGGAAGAAGTG | TTGCTCACAAAGGTCACAGG |

A B

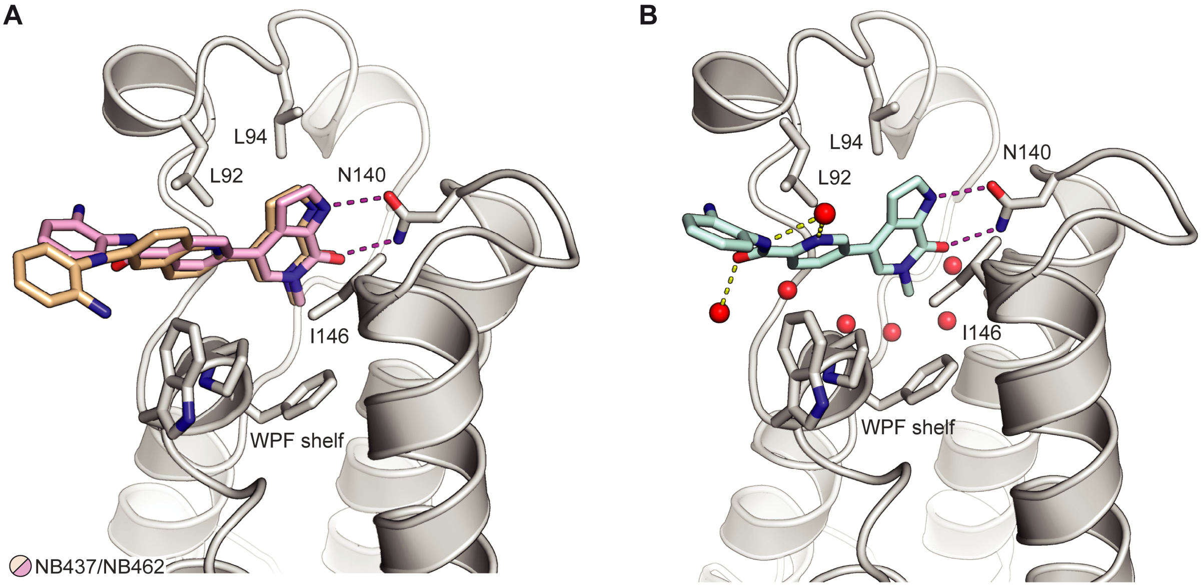


**Supporting Figure S1.** Structures of BRD4 BD1 with bound dual BET/HDAC inhibitors. (A) Superimposition of the crystal structures of BRD4 BD1 in complex with NB437 (light orange stick model) and NB462 (pink stick model). For clarity, only the structure of the protein chain of the NB462 complex is shown (gray ribbon model), with selected side chains interacting with the inhibitor highlighted as stick model. (B) Structure of BRD4 BD1 in complex with NB500. Selected water molecules in the binding pocket are shown as red spheres. Hydrogen bonds of the inhibitor with the protein (magenta dashed lines) or water molecules (yellow dashed lines) are highlighted.

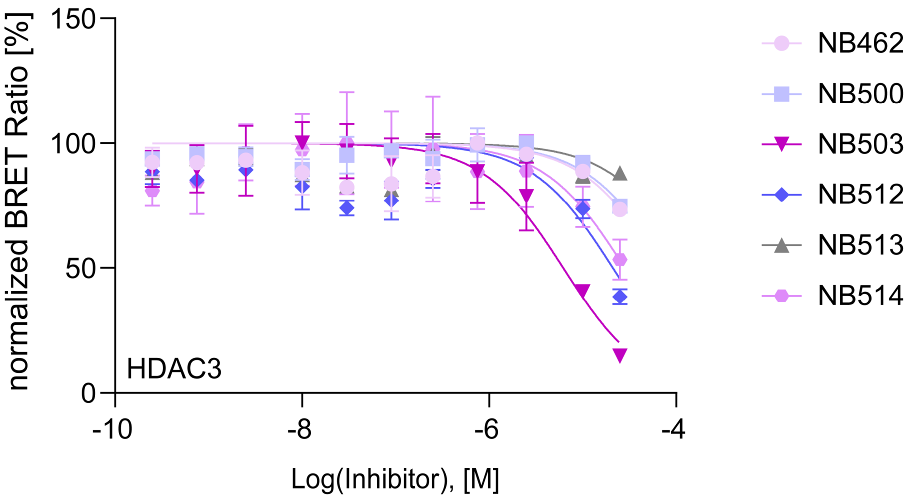


**Supporting Figure S2.** Representative NanoBRET data and fits of dual BET/HDAC inhibitors binding to HDAC3.

### HPLC purity

#### Blank measurement

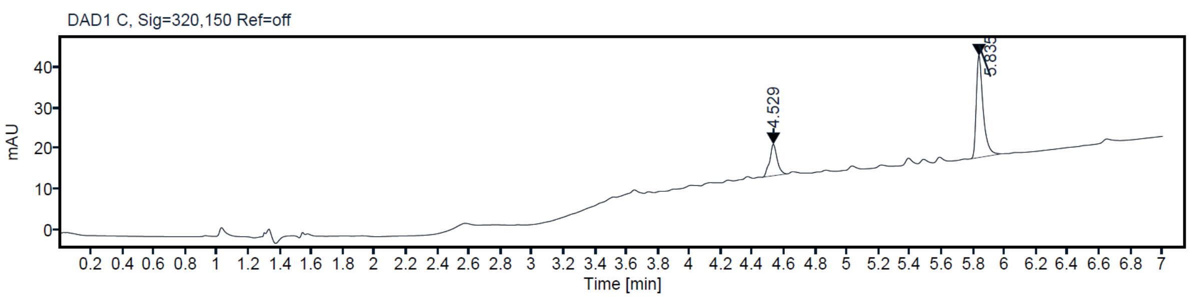


## NB161

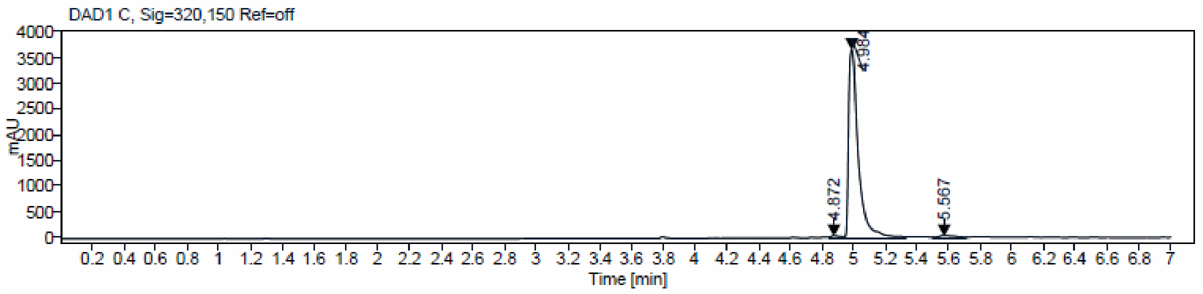


**Sample Purity**

Signal Description DAD1 C, Sig=320,150 Ref=off

| Sample Name | Name | RT | Width | Area | Area% | Height |
|---|---|---|---|---|---|---|
| NB161 | | 4.872 | 0.088 | 200.9542 | 1.24 | 46.0820 |
| NB161 | | 4.984 | 0.058 | 15612.18 55 | 96.48 | 3644.0142 |
| NB161 | | 5.567 | 0.132 | 368.6589 | 2.28 | 45.0985 |

| | |
|---|---|
| Max Area% | 96.480 |
| UV Signal Purity>95% | Pass |

## NB390

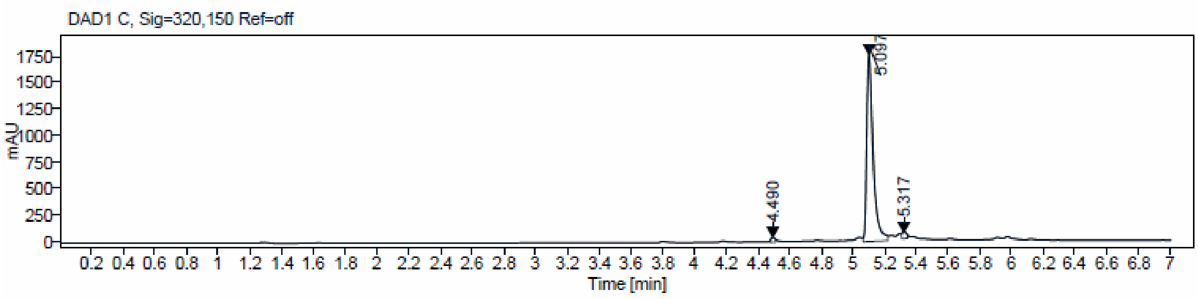


**Sample Purity**

Signal Description DAD1 C, Sig=320,150 Ref=off

| Sample Name | Name | RT | Width | Area | Area% | Height |
|---|---|---|---|---|---|---|
| NB390 | | 4.490 | 0.029 | 54.1471 | 1.10 | 36.5637 |
| NB390 | | 5.097 | 0.040 | 4800.5356 | 97.39 | 1741.9381 |
| NB390 | | 5.317 | 0.031 | 74.5046 | 1.51 | 45.1006 |

Max Area% 97.390

UV Signal Purity>95% Pass

## NB437

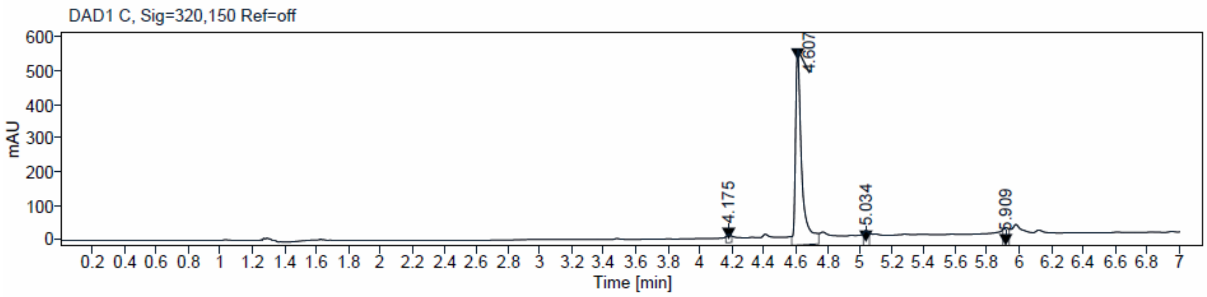


**Sample Purity**

Signal Description DAD1 C, Sig=320,150 Ref=off

| Sample Name | Name | RT | Width | Area | Area% | Height |
|---|---|---|---|---|---|---|
| NB437 | | 4.175 | 0.033 | 14.8471 | 1.11 | 8.6162 |
| NB437 | | 4.607 | 0.033 | 1286.3580 | 95.93 | 547.6205 |
| NB437 | | 5.034 | 0.026 | 14.7228 | 1.10 | 9.8781 |
| NB437 | | 5.909 | 0.038 | 25.0186 | 1.87 | 13.0276 |

Max Area% 95.929

UV Signal Purity>95% Pass

## NB462

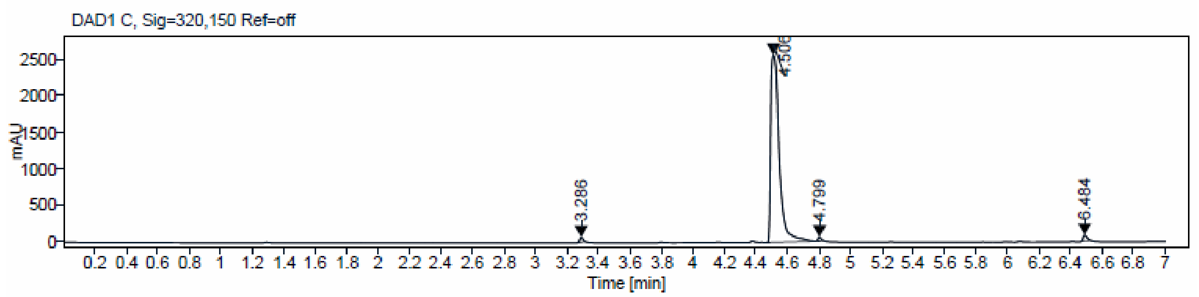


**Sample Purity**

Signal Description DAD1 C, Sig=320,150 Ref=off

| Sample Name | Name | RT | Width | Area | Area% | Height |
|---|---|---|---|---|---|---|
| NB462 | | 3.286 | 0.024 | 91.5371 | 0.92 | 66.2465 |
| NB462 | | 4.506 | 0.053 | 9631.3271 | 97.13 | 2577.7595 |
| NB462 | | 4.799 | 0.023 | 65.2411 | 0.66 | 52.4126 |
| NB462 | | 6.484 | 0.030 | 127.9308 | 1.29 | 76.5870 |

Max Area% 97.129

UV Signal Purity>95% Pass

## NB469

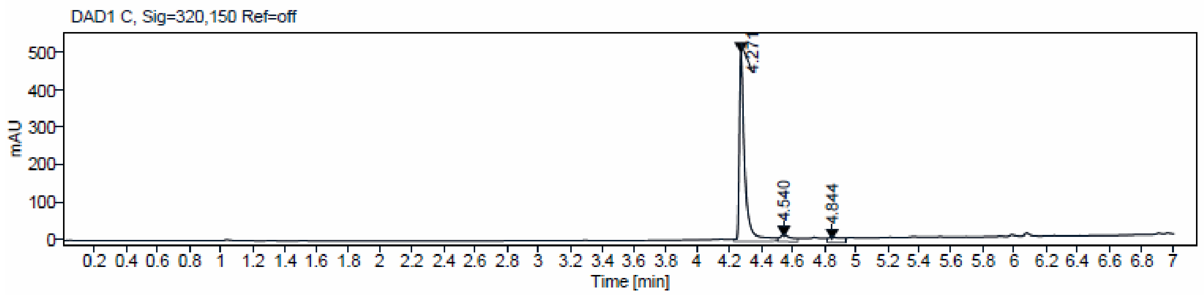


**Sample Purity**

Signal Description DAD1 C, Sig=320,150 Ref=off

| Sample Name | Name | RT | Width | Area | Area% | Height |
|---|---|---|---|---|---|---|
| NB469 | | 4.271 | 0.031 | 1119.2117 | 97.09 | 499.0727 |
| NB469 | | 4.540 | 0.034 | 23.7507 | 2.06 | 10.0126 |
| NB469 | | 4.844 | 0.029 | 9.8229 | 0.85 | 4.3816 |

Max Area% 97.088

UV Signal Purity>95% Pass

## NB470

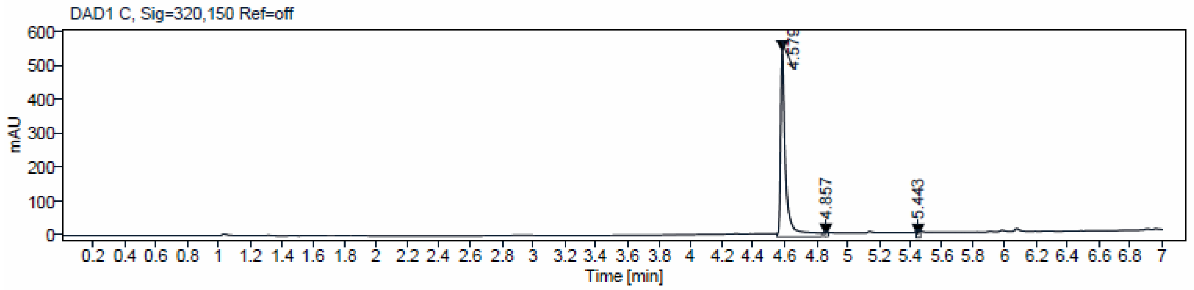


**Sample Purity**

Signal Description DAD1 C, Sig=320,150 Ref=off

| Sample Name | Name | RT | Width | Area | Area% | Height |
|---|---|---|---|---|---|---|
| NB470 | | 4.579 | 0.027 | 1101.4595 | 98.78 | 544.6414 |
| NB470 | | 4.857 | 0.024 | 5.1854 | 0.47 | 4.1291 |
| NB470 | | 5.443 | 0.026 | 8.4550 | 0.76 | 6.4661 |

Max Area% 98.777

UV Signal Purity>95% Pass

## NB480

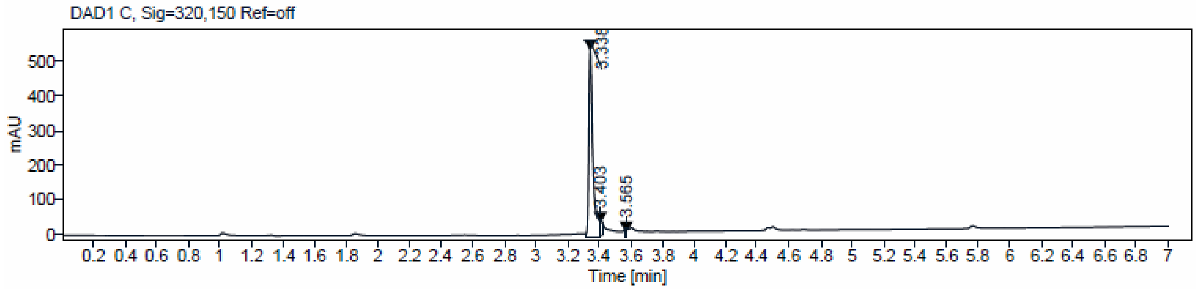


**Sample Purity**

Signal Description DAD1 C, Sig=320,150 Ref=off

| Sample Name | Name | RT | Width | Area | Area% | Height |
|---|---|---|---|---|---|---|
| NB480 | | 3.338 | 0.024 | 948.8326 | 95.77 | 538.9740 |
| NB480 | | 3.403 | 0.020 | 34.6199 | 3.49 | 34.1772 |
| NB480 | | 3.565 | 0.015 | 7.3165 | 0.74 | 9.2308 |

Max Area% 95.767

UV Signal Purity>95% Pass

**NB500**

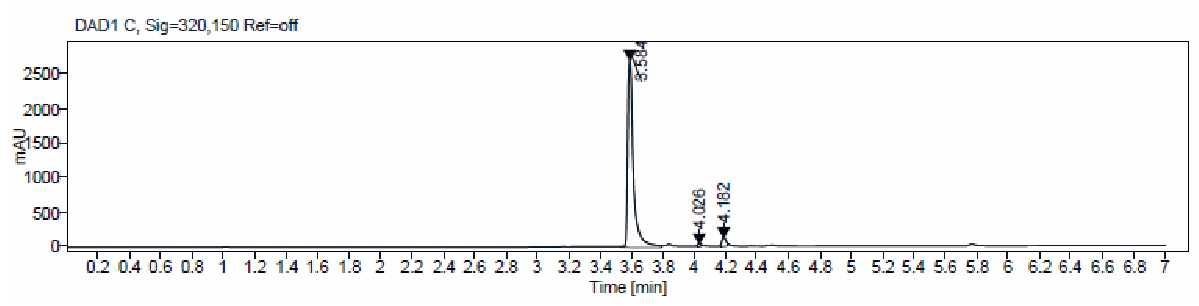


**Sample Purity**

Signal Description DAD1 C, Sig=320,150 Ref=off

| Sample Name | Name | RT | Width | Area | Area% | Height |
|---|---|---|---|---|---|---|
| NB500 | | 3.584 | 0.033 | 6617.2954 | 97.23 | 2705.4158 |
| NB500 | | 4.026 | 0.020 | 24.7576 | 0.36 | 22.2138 |
| NB500 | | 4.182 | 0.026 | 163.5289 | 2.40 | 108.7853 |

Max Area% 97.233

UV Signal Purity>95% Pass

**NB501**

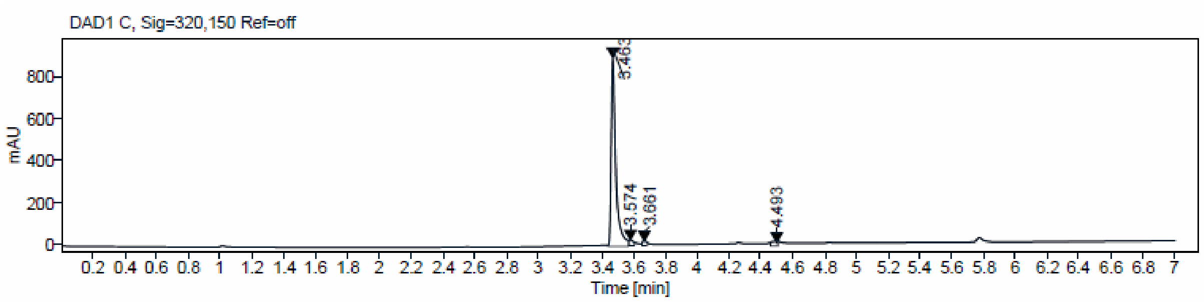


**Sample Purity**

Signal Description DAD1 C, Sig=320,150 Ref=off

| Sample Name | Name | RT | Width | Area | Area% | Height |
|---|---|---|---|---|---|---|
| NB501 | | 3.463 | 0.027 | 1781.5571 | 96.17 | 889.6945 |
| NB501 | | 3.574 | 0.029 | 30.4943 | 1.65 | 19.2803 |
| NB501 | | 3.661 | 0.023 | 18.5934 | 1.00 | 13.7266 |
| NB501 | | 4.493 | 0.047 | 21.9083 | 1.18 | 9.7645 |

Max Area% 96.168

UV Signal Purity>95% Pass

**NB502**

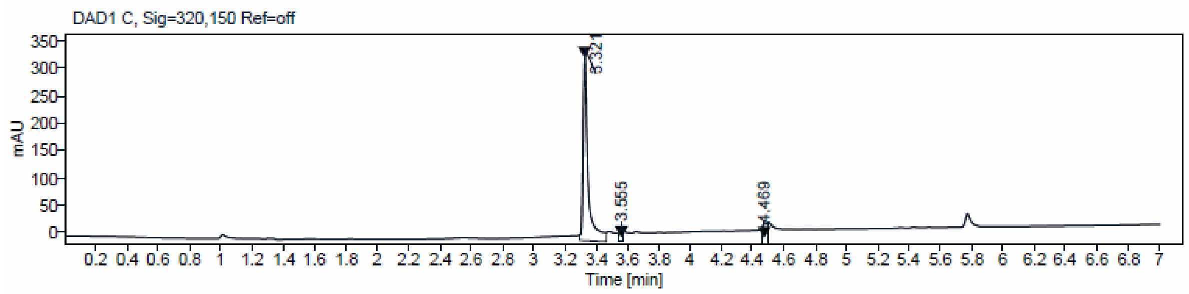


**Sample Purity**

Signal Description DAD1 C, Sig=320,150 Ref=off

| Sample Name | Name | RT | Width | Area | Area% | Height |
|---|---|---|---|---|---|---|
| NB502 | | 3.321 | 0.025 | 634.7424 | 95.96 | 334.3840 |
| NB502 | | 3.555 | 0.018 | 6.8211 | 1.03 | 6.7317 |
| NB502 | | 4.469 | 0.039 | 19.9128 | 3.01 | 9.8161 |

Max Area% 95.958

UV Signal Purity>95% Pass

**NB503**

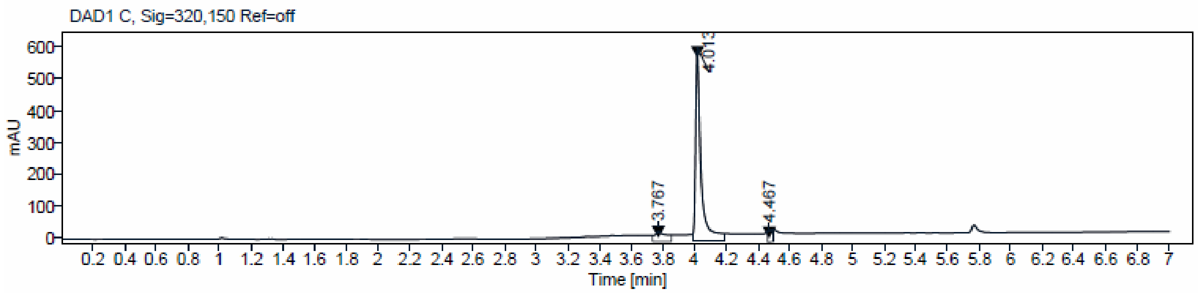


**Sample Purity**

Signal Description DAD1 C, Sig=320,150 Ref=off

| Sample Name | Name | RT | Width | Area | Area% | Height |
|---|---|---|---|---|---|---|
| NB503 | | 3.767 | 0.031 | 25.0035 | 2.02 | 11.1851 |
| NB503 | | 4.013 | 0.028 | 1195.5093 | 96.53 | 574.0806 |
| NB503 | | 4.467 | 0.037 | 17.9701 | 1.45 | 9.6678 |

Max Area% 96.530

UV Signal Purity>95% Pass

**NB507**

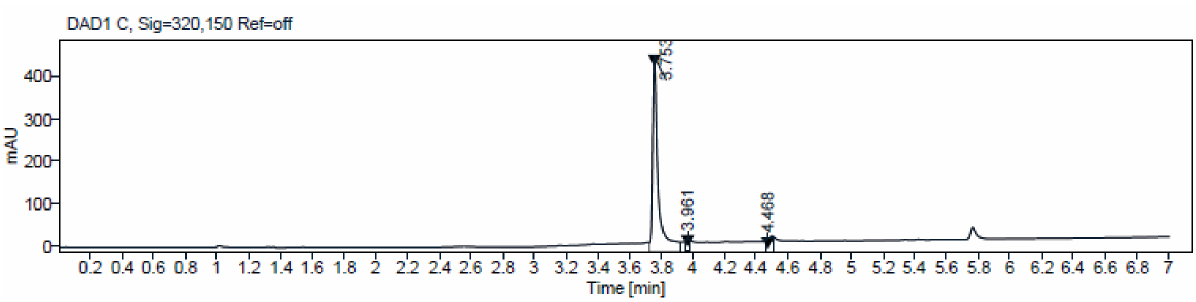


**Sample Purity**

Signal Description DAD1 C, Sig=320,150 Ref=off

| Sample Name | Name | RT | Width | Area | Area% | Height |
|---|---|---|---|---|---|---|
| NB507 | | 3.753 | 0.028 | 914.1124 | 97.29 | 433.9325 |
| NB507 | | 3.961 | 0.023 | 9.2851 | 0.99 | 7.2258 |
| NB507 | | 4.468 | 0.046 | 16.1563 | 1.72 | 6.7971 |

| | |
|---|---|
| Max Area% | 97.292 |
| UV Signal Purity>95% | Pass |

**NB512**

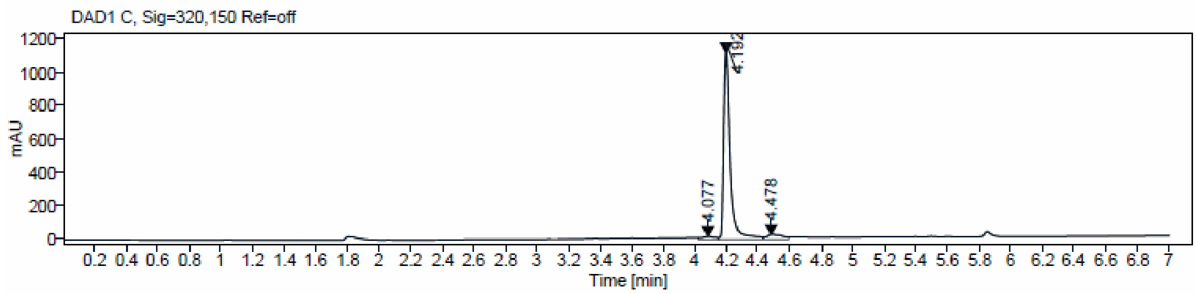


**Sample Purity**

Signal Description DAD1 C, Sig=320,150 Ref=off

| Sample Name | Name | RT | Width | Area | Area% | Height |
|---|---|---|---|---|---|---|
| NB533 | | 4.077 | 0.047 | 33.9675 | 1.10 | 10.5429 |
| NB533 | | 4.192 | 0.038 | 2975.9663 | 96.45 | 1112.9037 |
| NB533 | | 4.478 | 0.055 | 75.6843 | 2.45 | 19.4026 |

| | |
|---|---|
| Max Area% | 96.446 |
| UV Signal Purity>95% | Pass |

## NB513

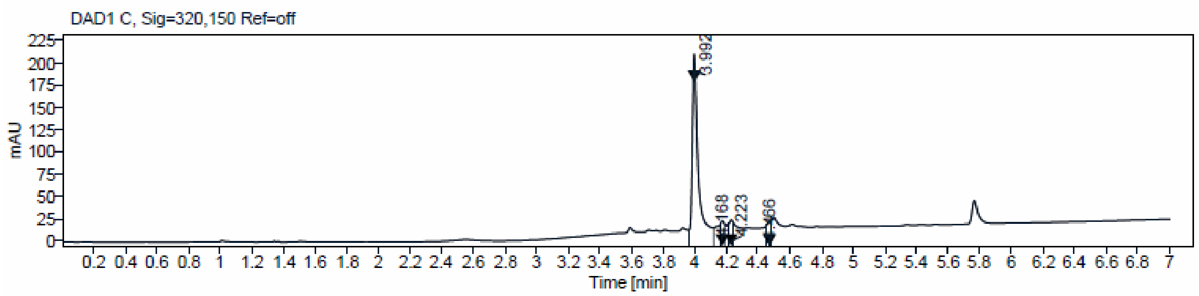


**Sample Purity**

Signal Description DAD1 C, Sig=320,150 Ref=off

| Sample Name | Name | RT | Width | Area | Area% | Height |
|---|---|---|---|---|---|---|
| NB513 | | 3.992 | 0.031 | 527.8755 | 95.67 | 207.0565 |
| NB513 | | 4.168 | 0.023 | 7.1770 | 1.30 | 5.7963 |
| NB513 | | 4.223 | 0.024 | 6.6385 | 1.20 | 5.2439 |
| NB513 | | 4.466 | 0.026 | 10.0878 | 1.83 | 6.9494 |

Max Area% 95.668

UV Signal Purity>95% Pass

## NB514

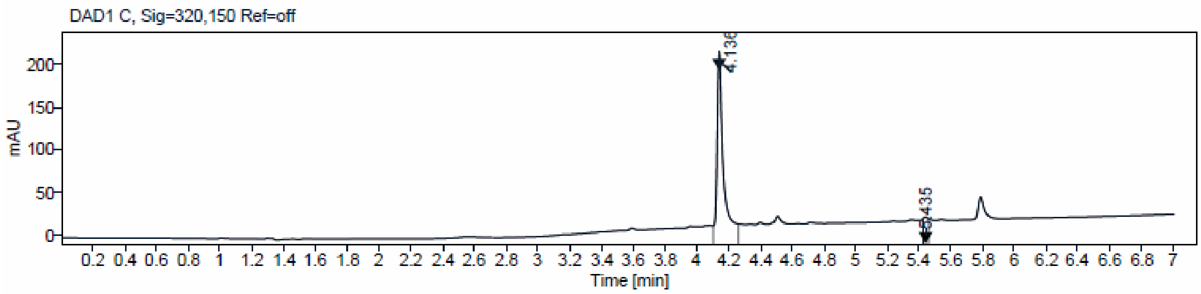


**Sample Purity**

Signal Description DAD1 C, Sig=320,150 Ref=off

| Sample Name | Name | RT | Width | Area | Area% | Height |
|---|---|---|---|---|---|---|
| NB514 | | 4.136 | 0.030 | 455.5850 | 97.95 | 203.8892 |
| NB514 | | 5.435 | 0.035 | 9.5244 | 2.05 | 5.4359 |

Max Area% 97.952

UV Signal Purity>95% Pass

### NMR Spectra

$^{1}H$ and $^{13}C$ Spectra of NB437 (**22**)

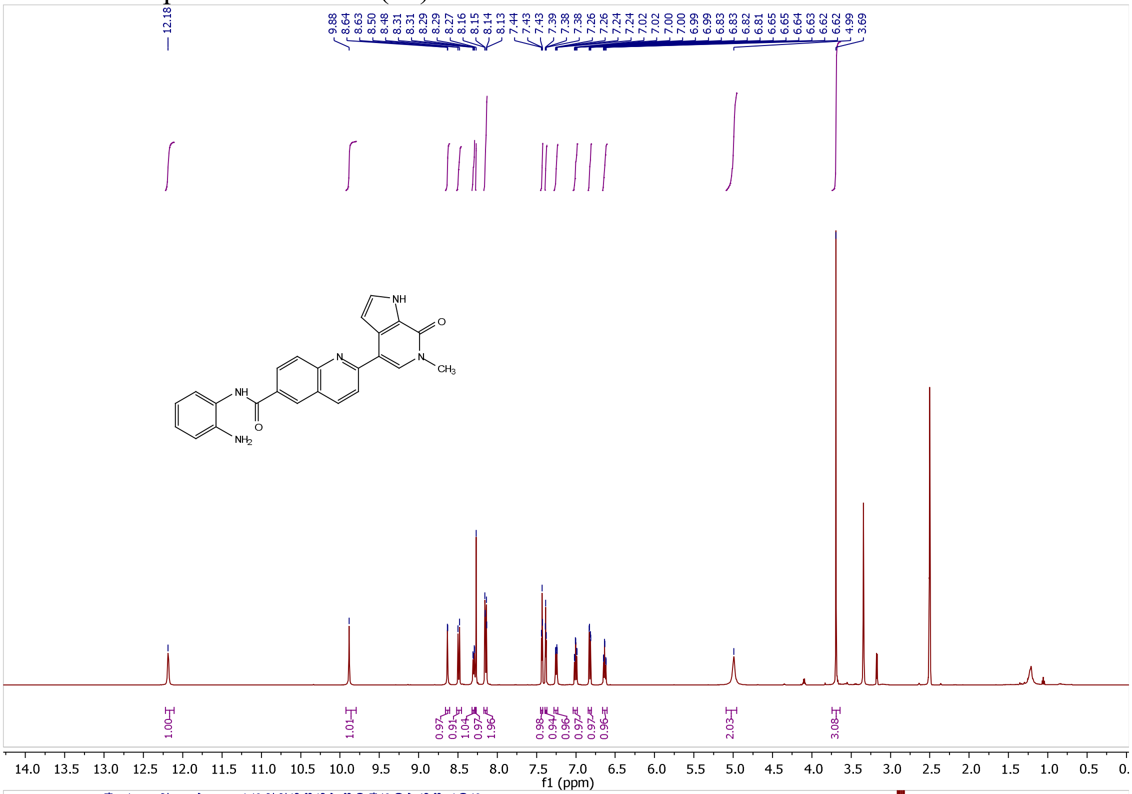


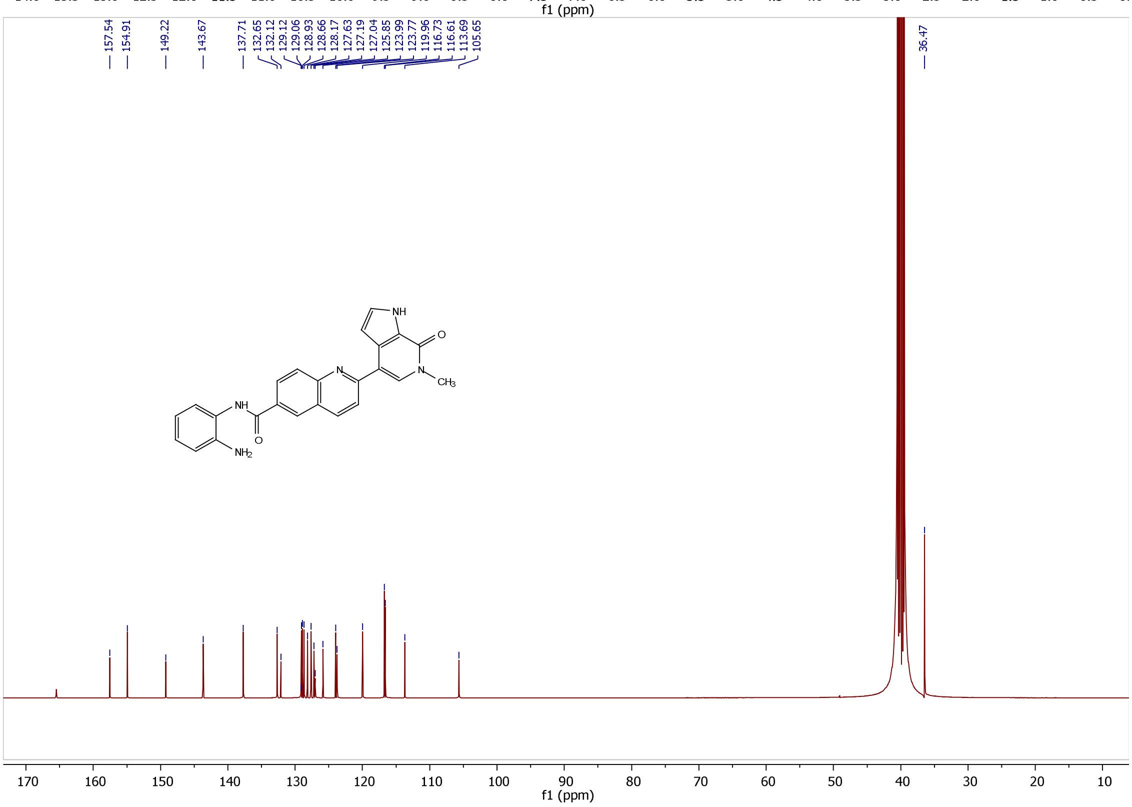

$^{1}H$ and $^{13}C$ Spectra of NB462 (**31a**)

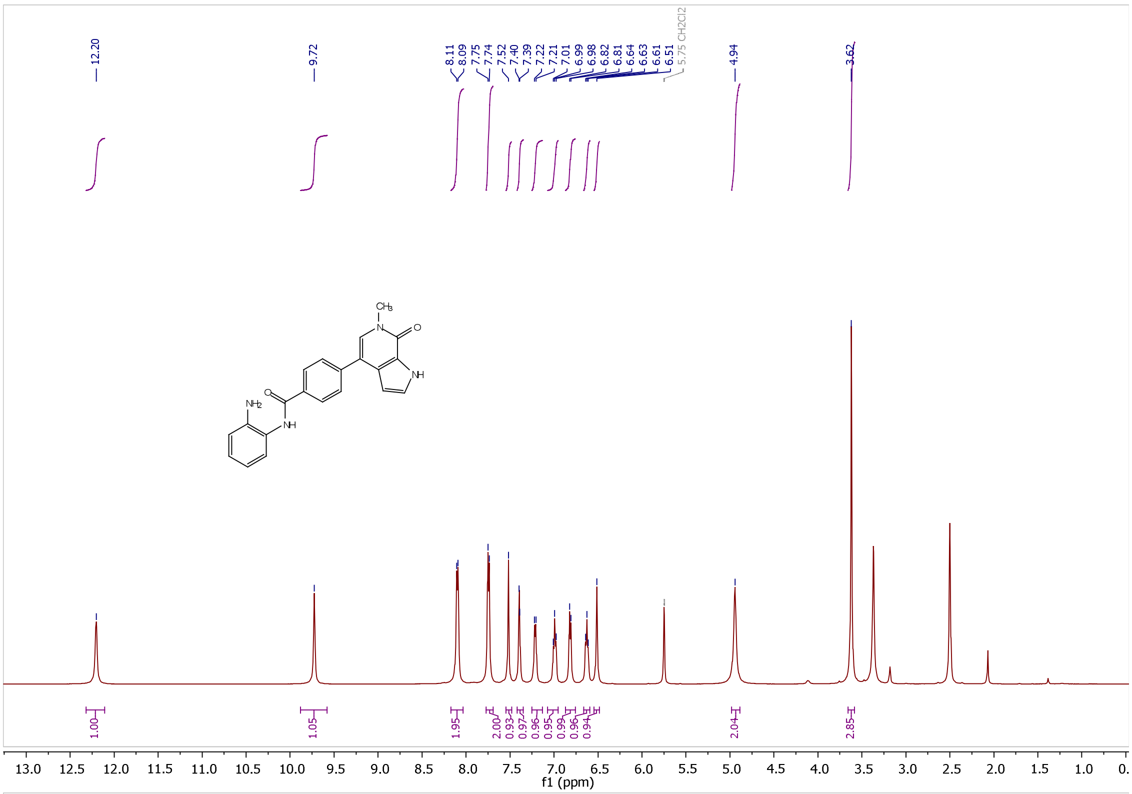


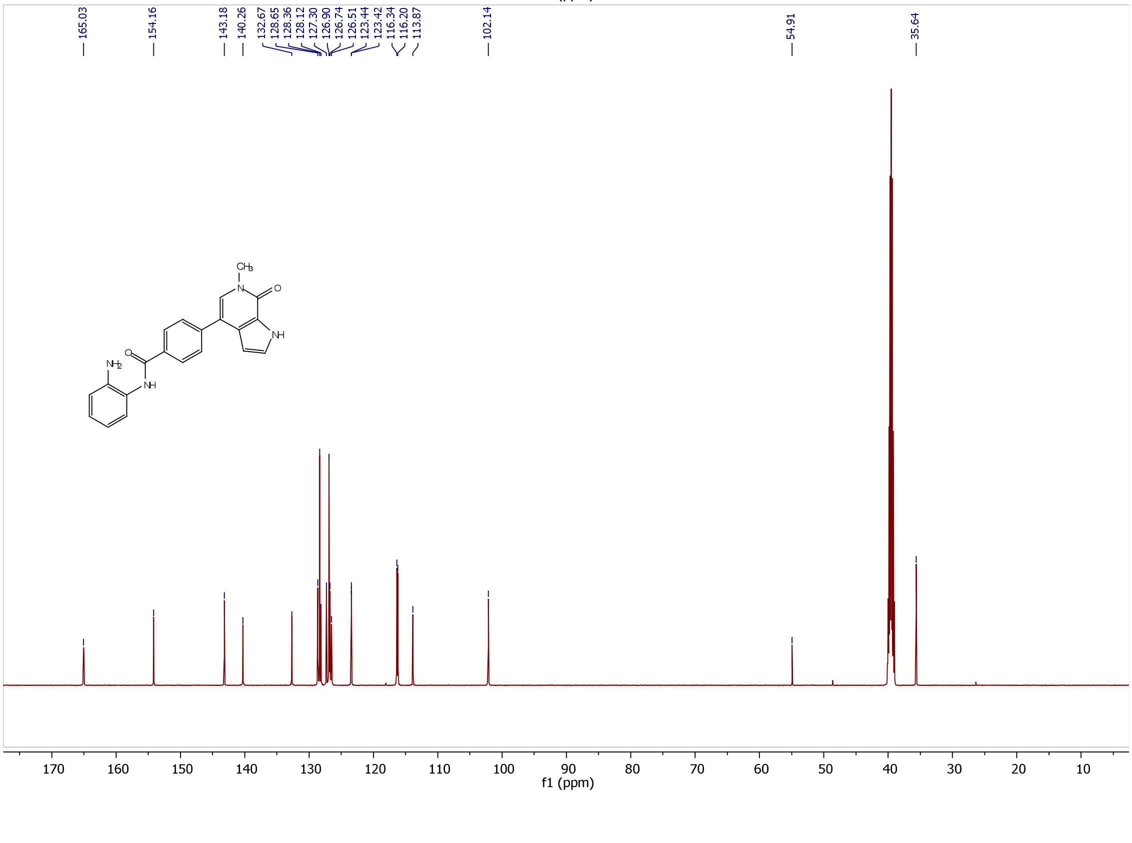

$^{1}H$ and $^{13}C$ Spectra of NB469 (**31b**)

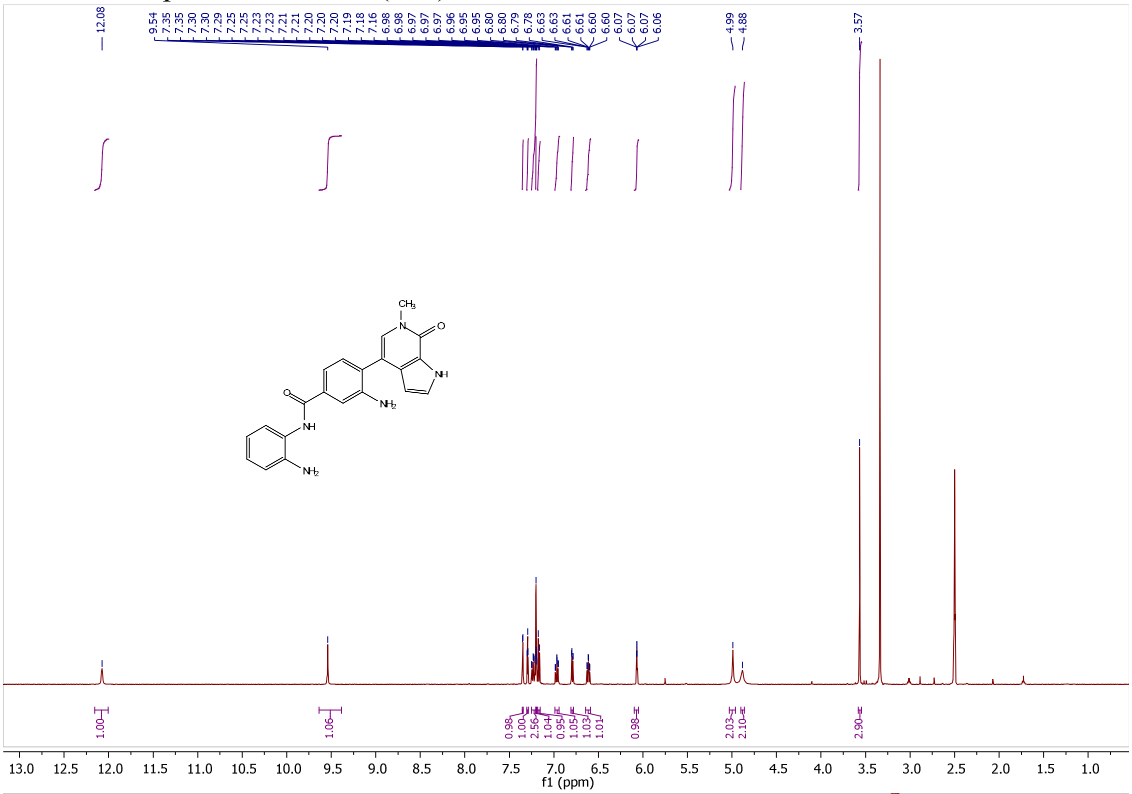


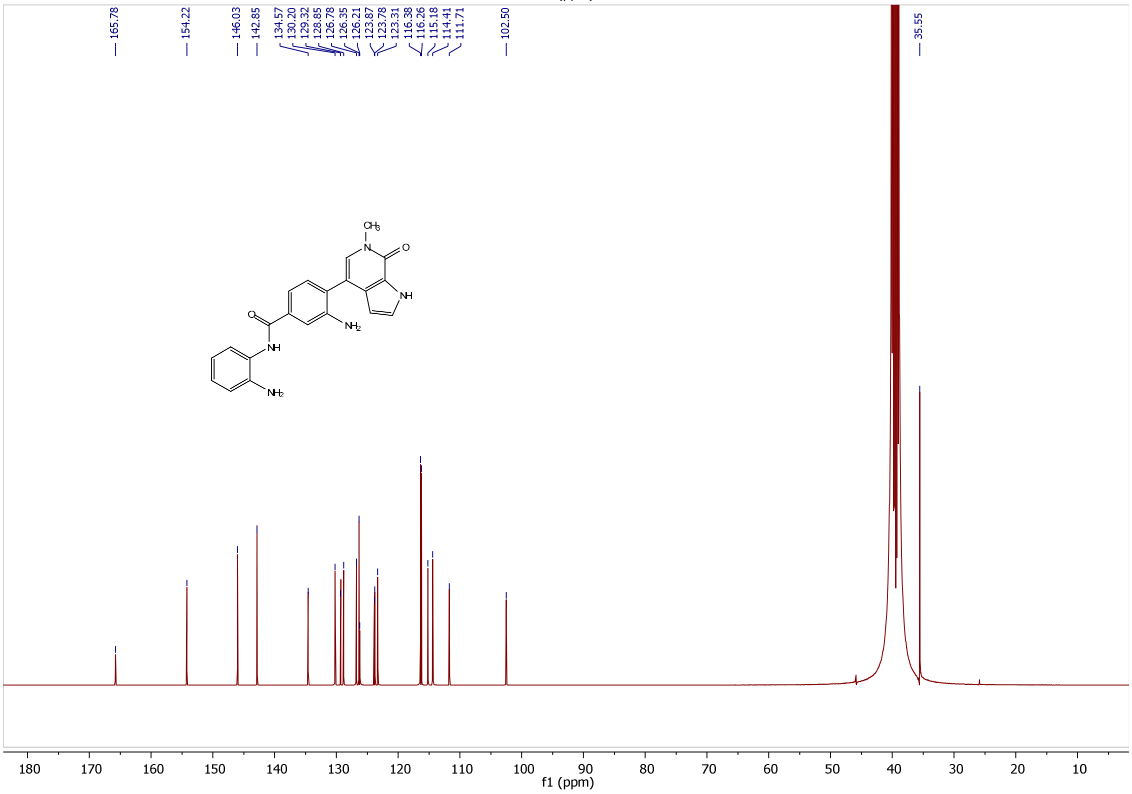

$^{1}H$ and $^{13}C$ Spectra of NB470 (**31c**)

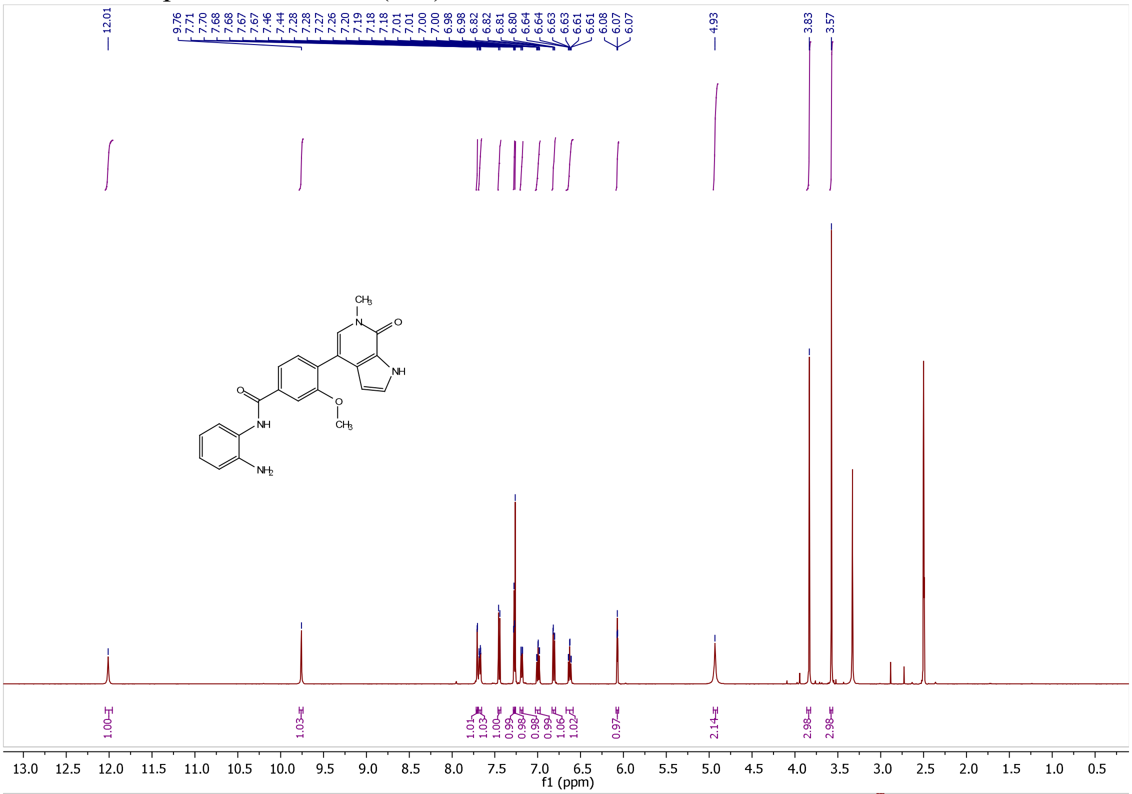


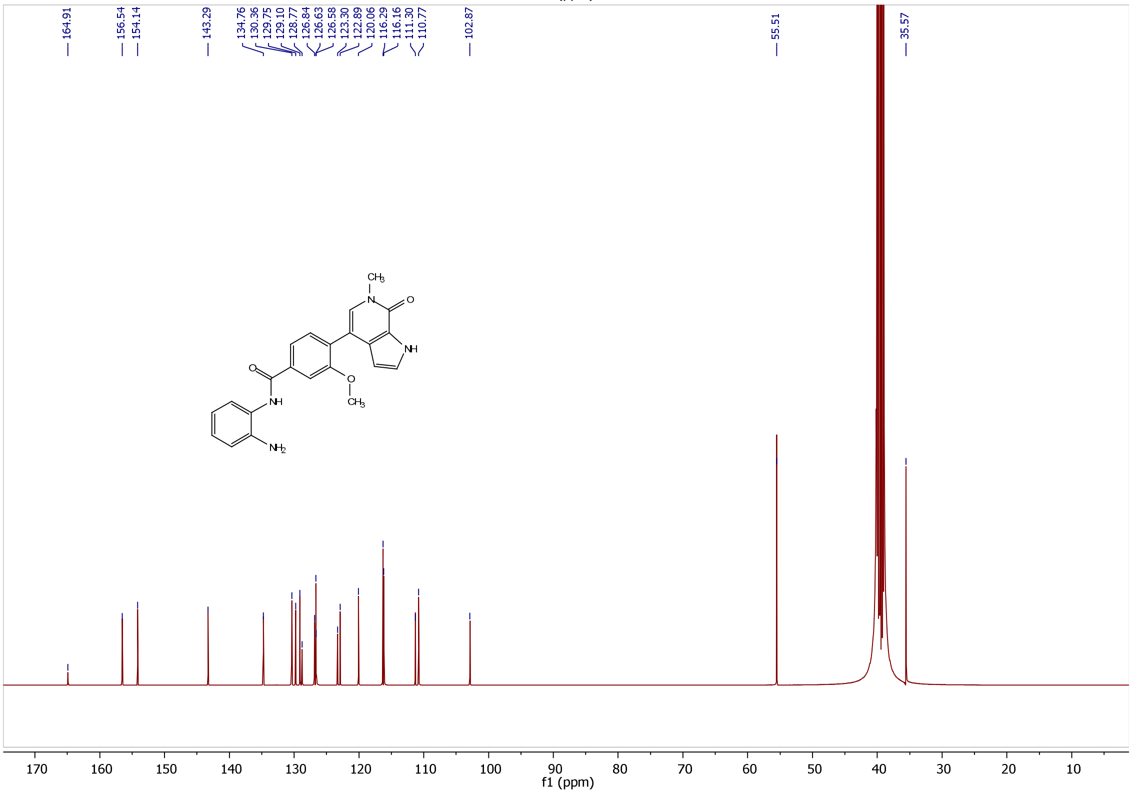

[1]H and [13]C Spectra of NB480 (**27**)

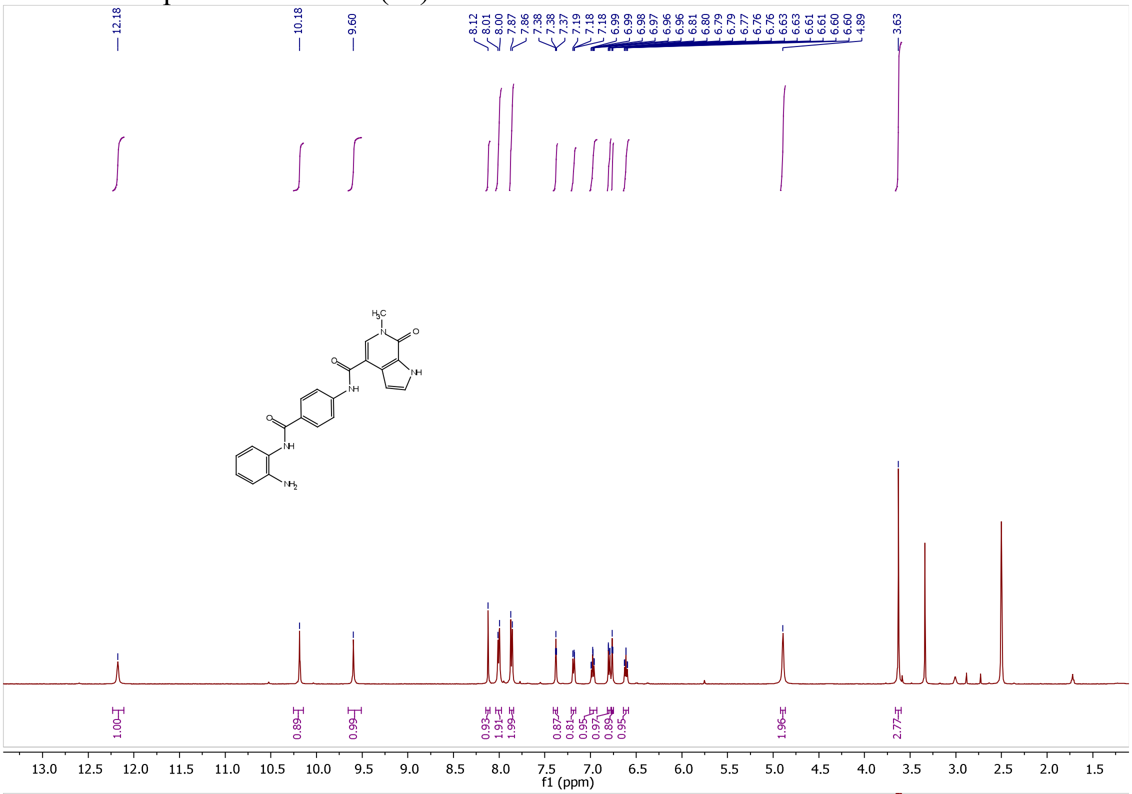

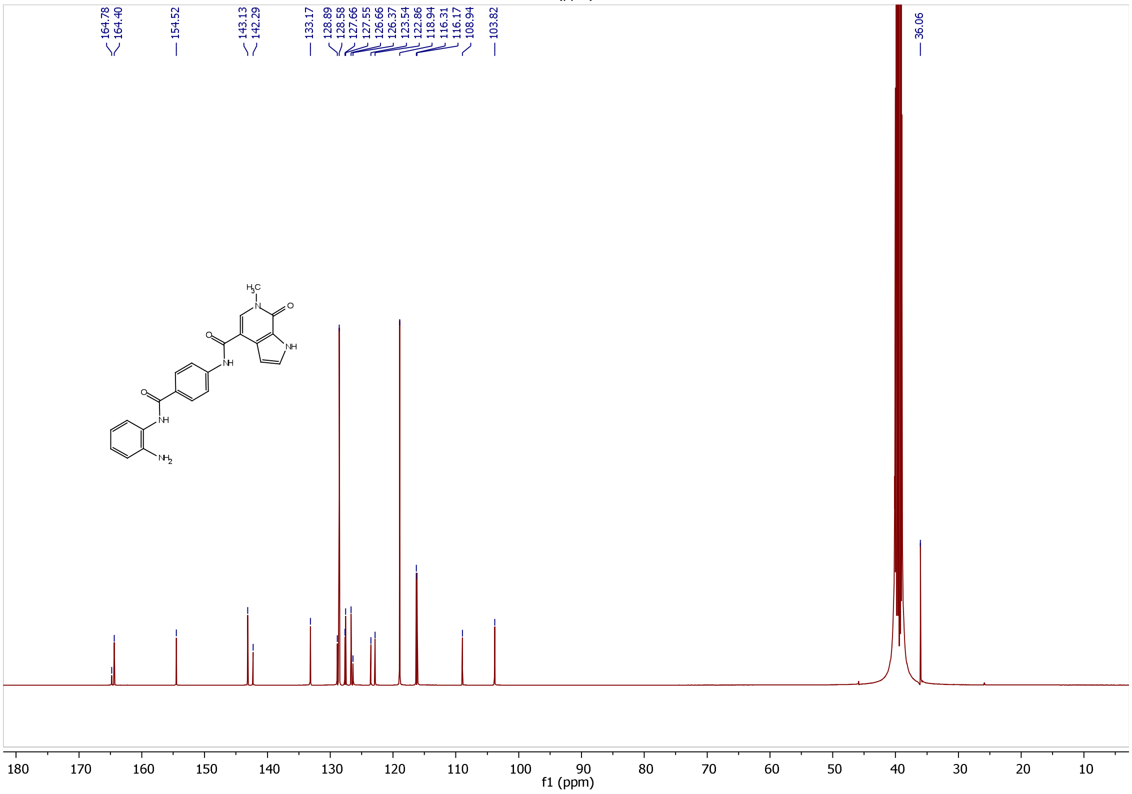

[1]H and [13]C Spectra of NB500 (**31d**)

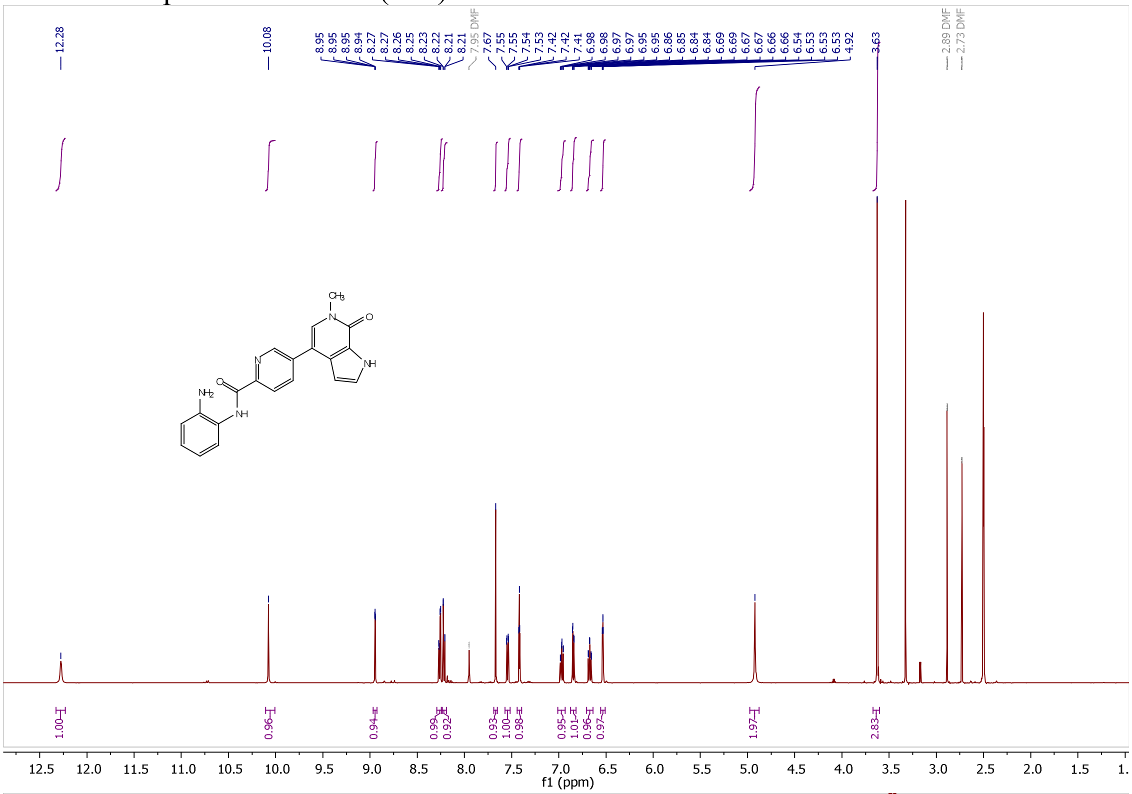


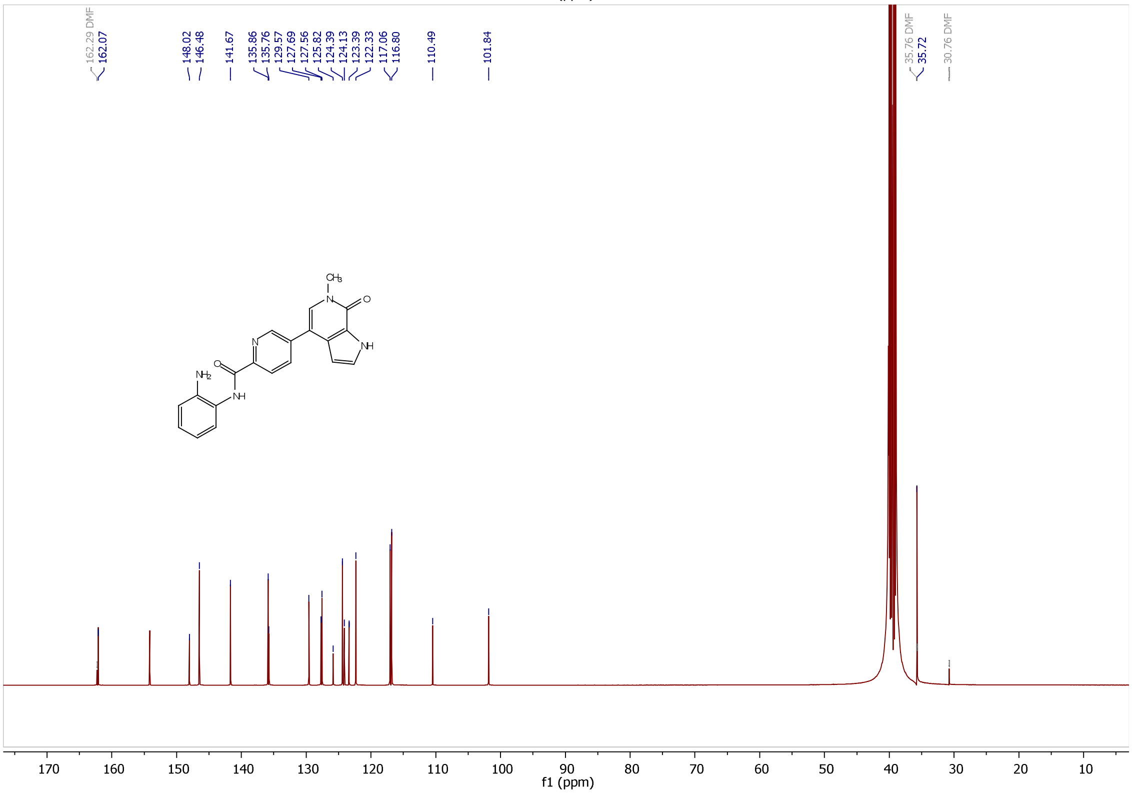

$^{1}H$ and $^{13}C$ Spectra of NB501 (**31e**)

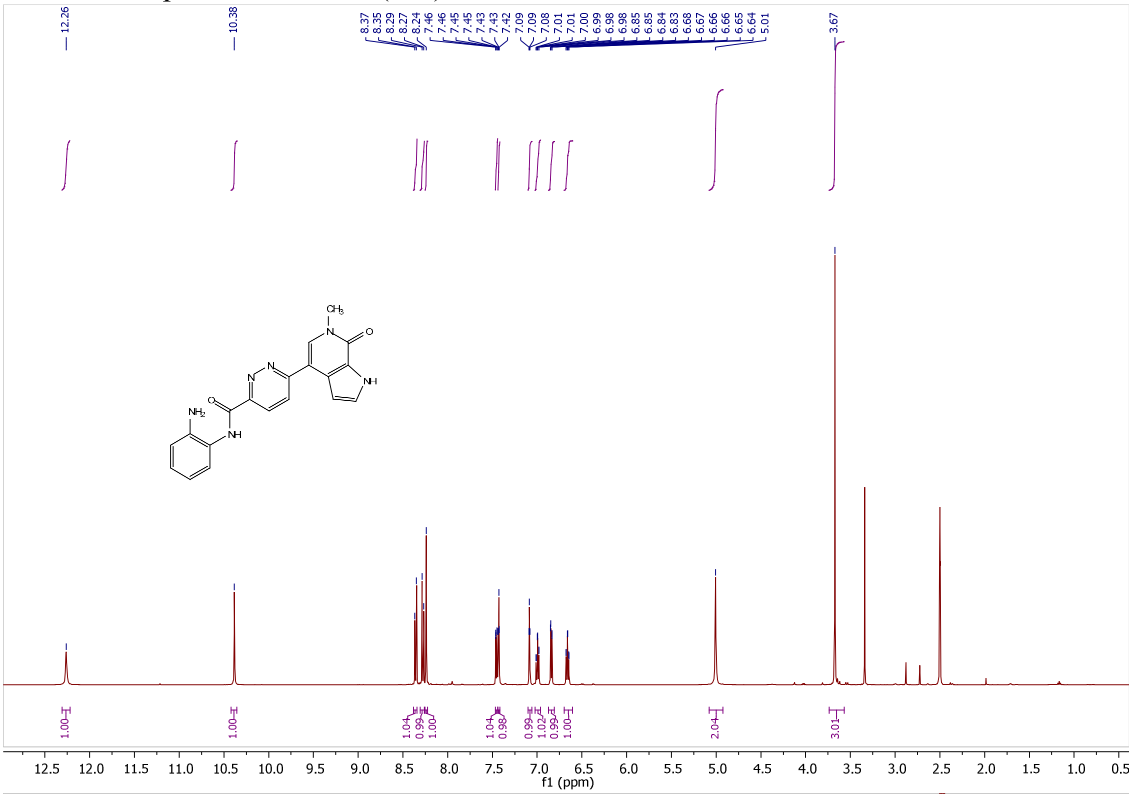


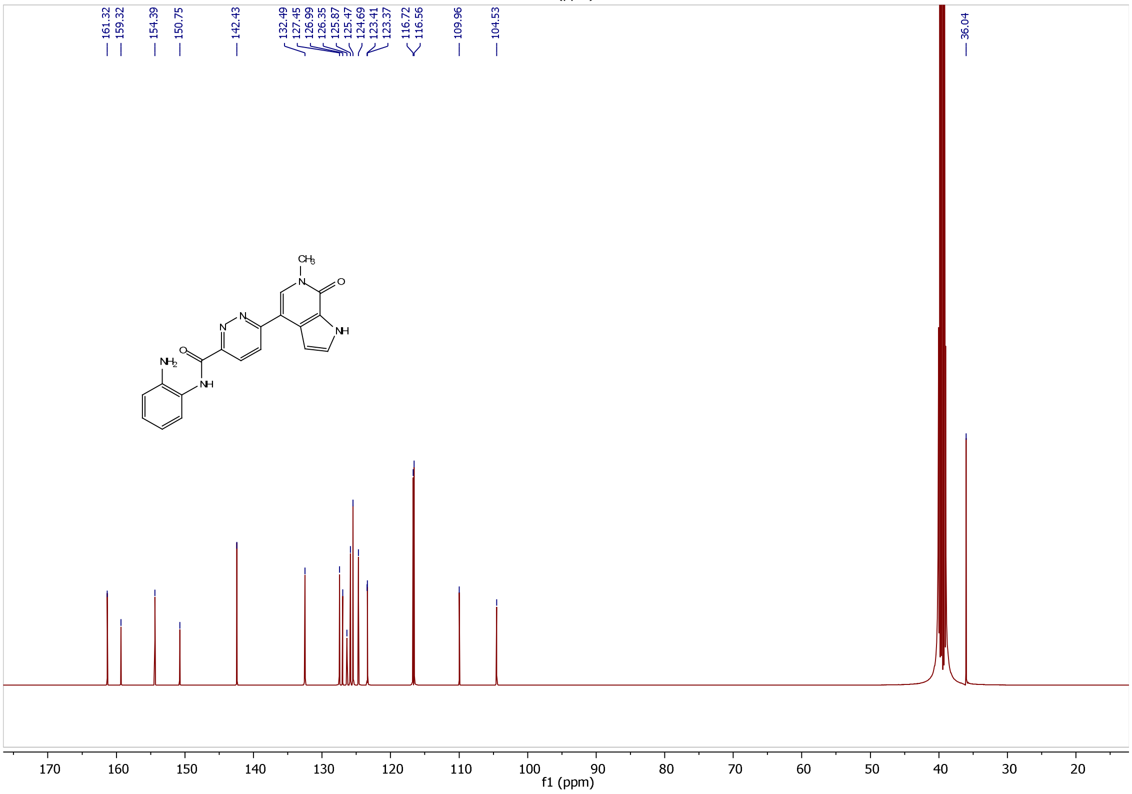

[1]H and [13]C Spectra of NB502 (**31f**)

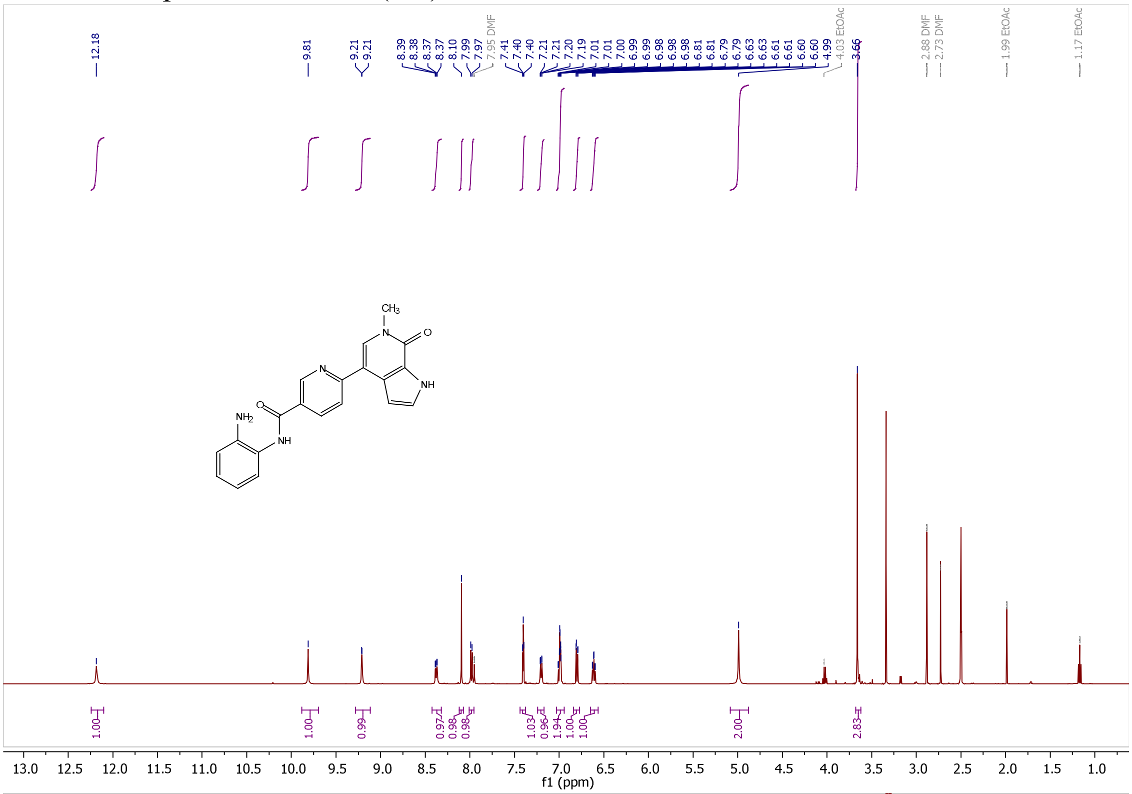

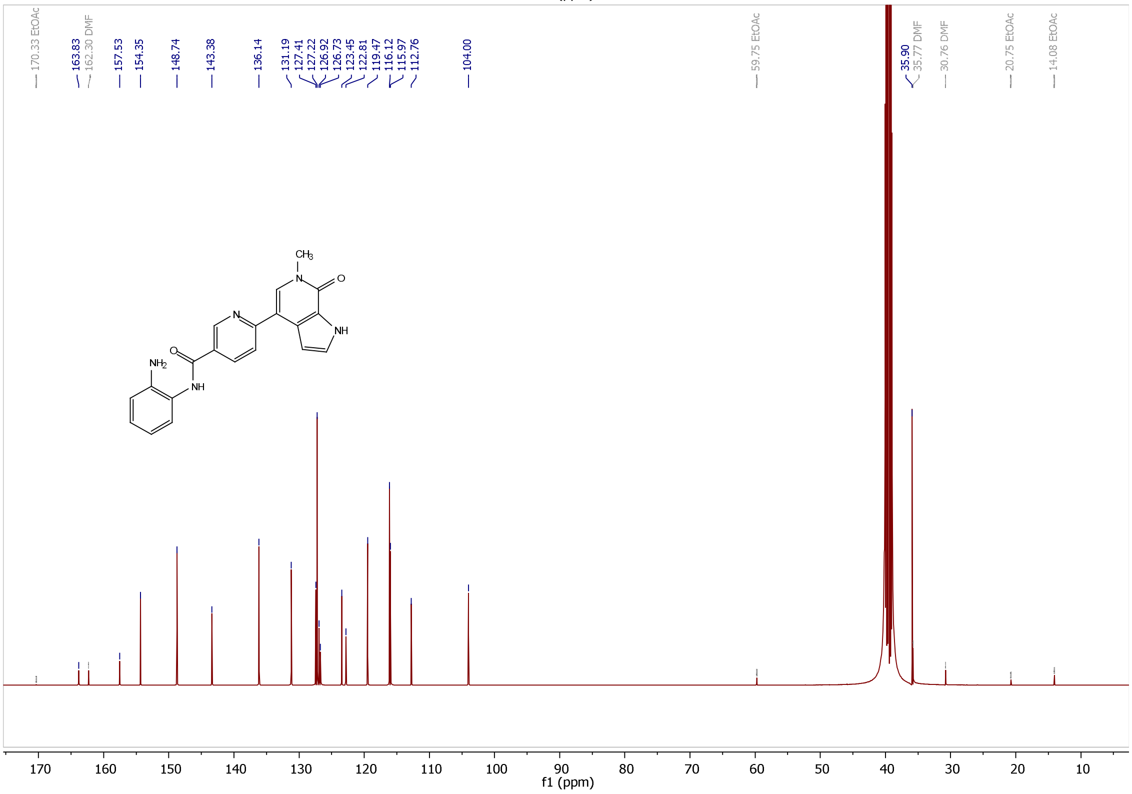

[1]H and [13]C Spectra of NB503 (**38**)

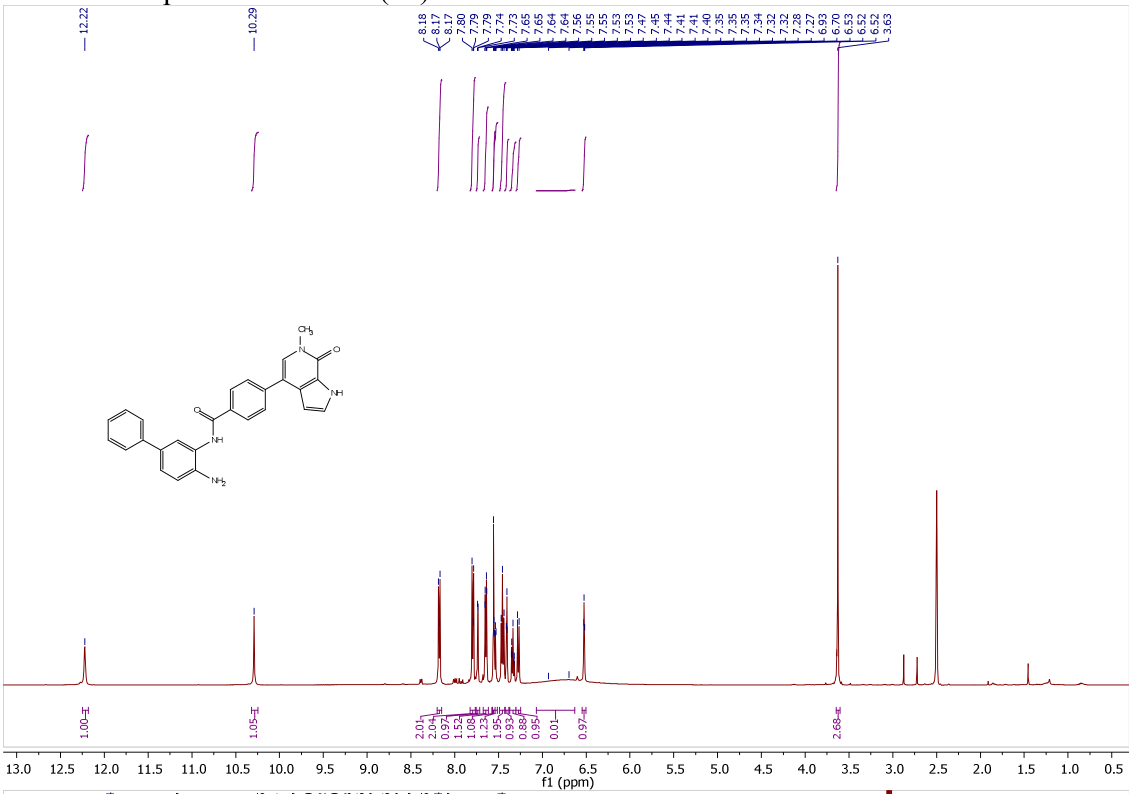


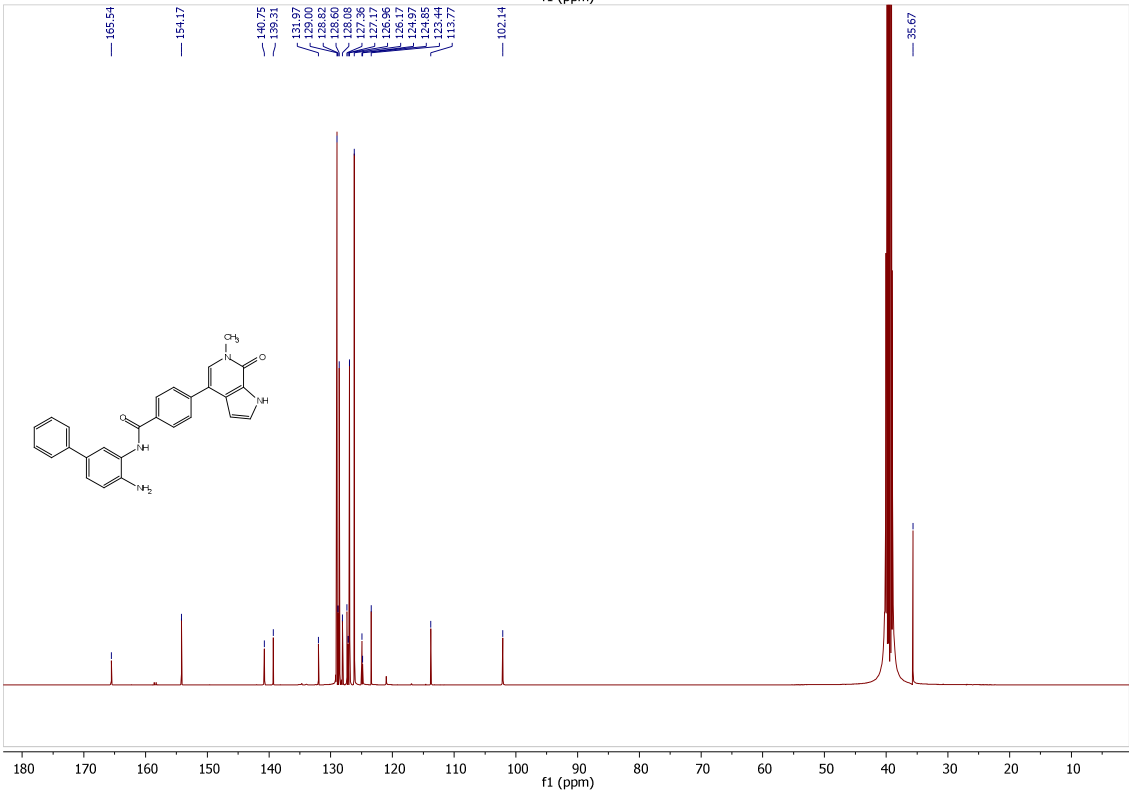

[1]H and [13]C Spectra of NB507 (**40**)

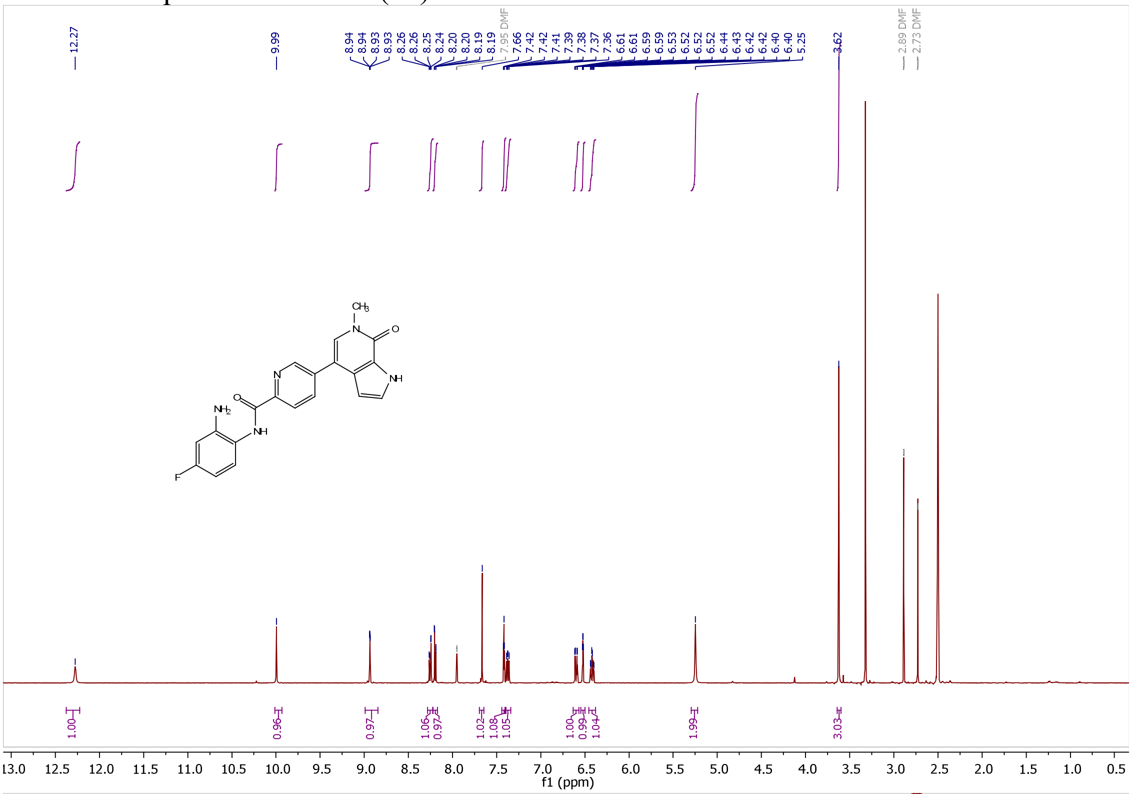


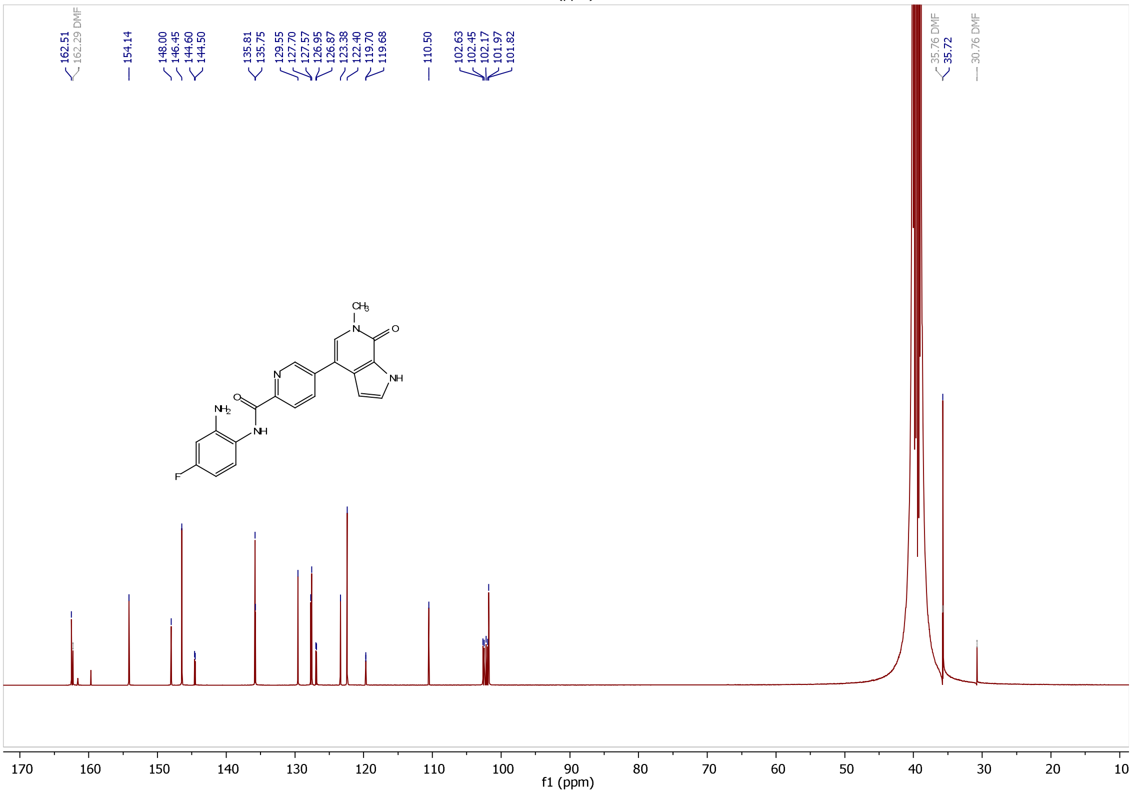

#### [1]H and [13]C Spectra of NB512 (**39a**)

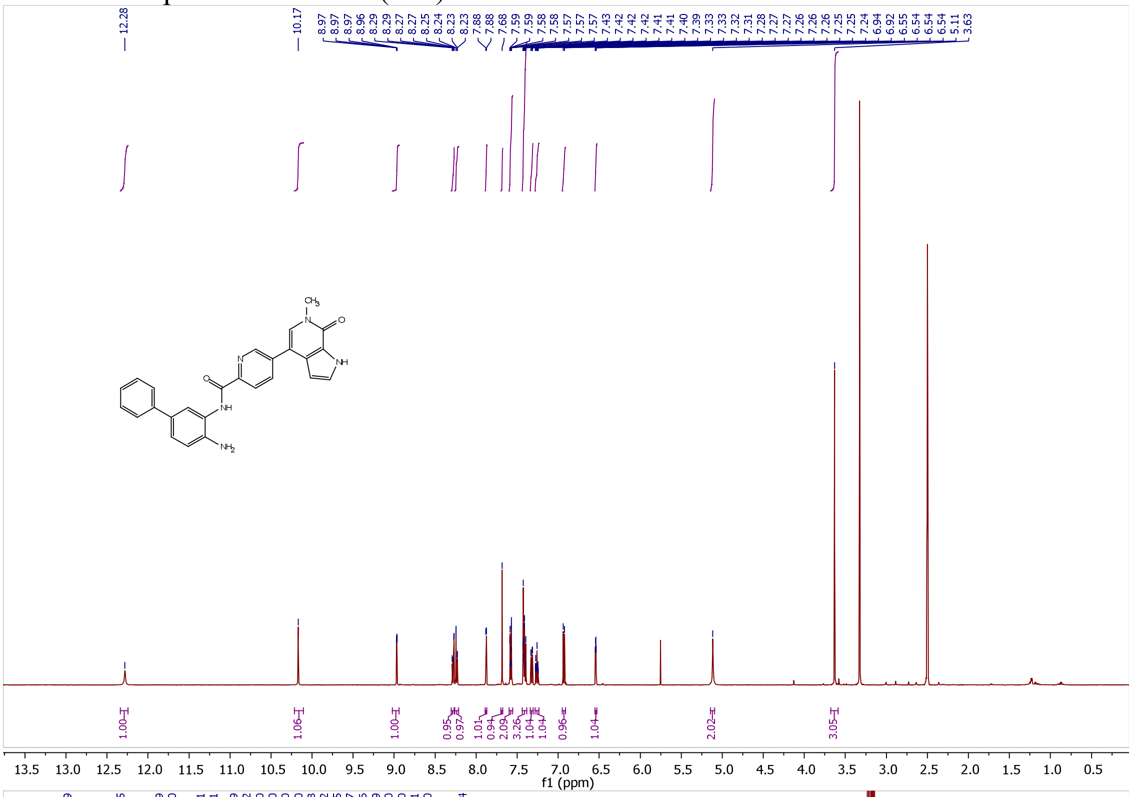


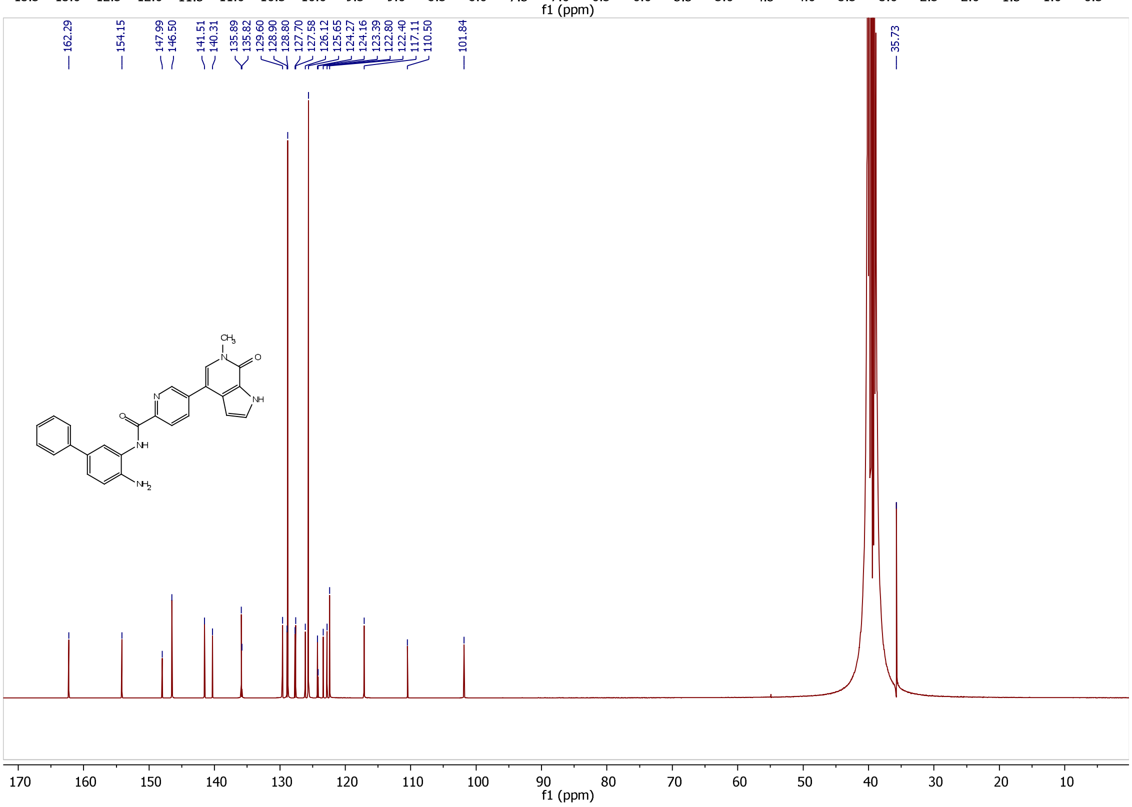

[1]H and [13]C Spectra of NB513 (**39b**)

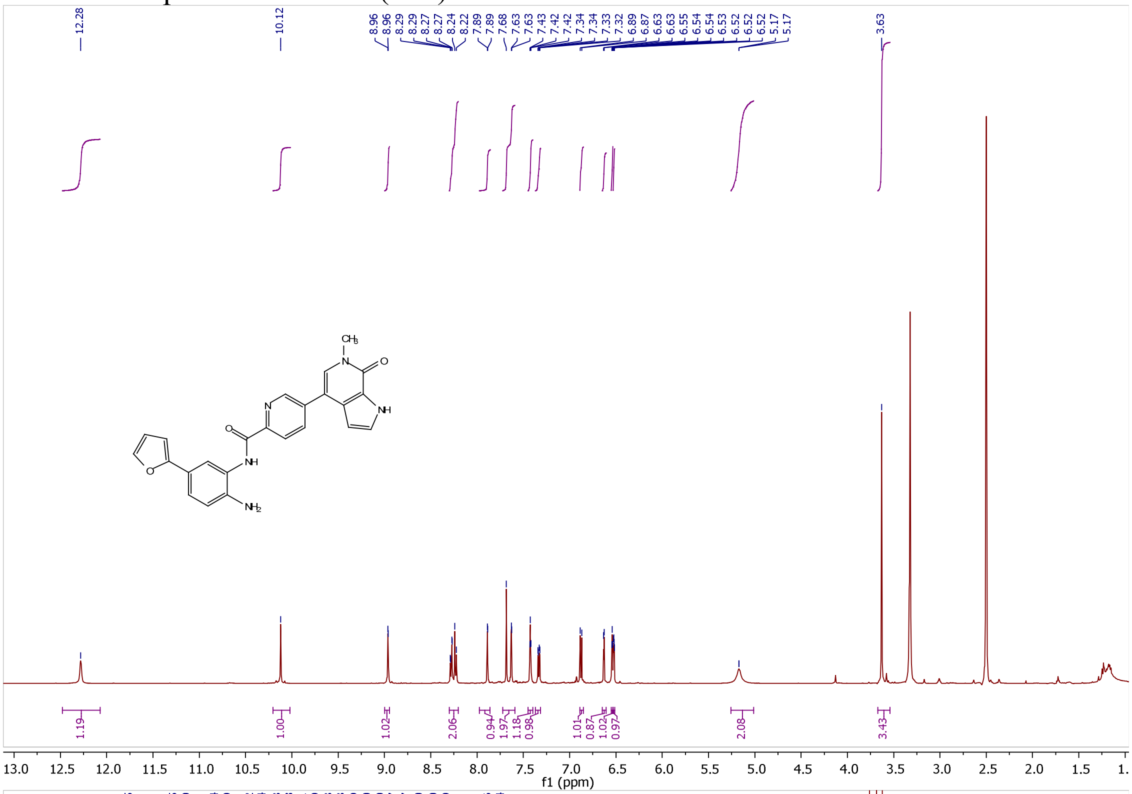

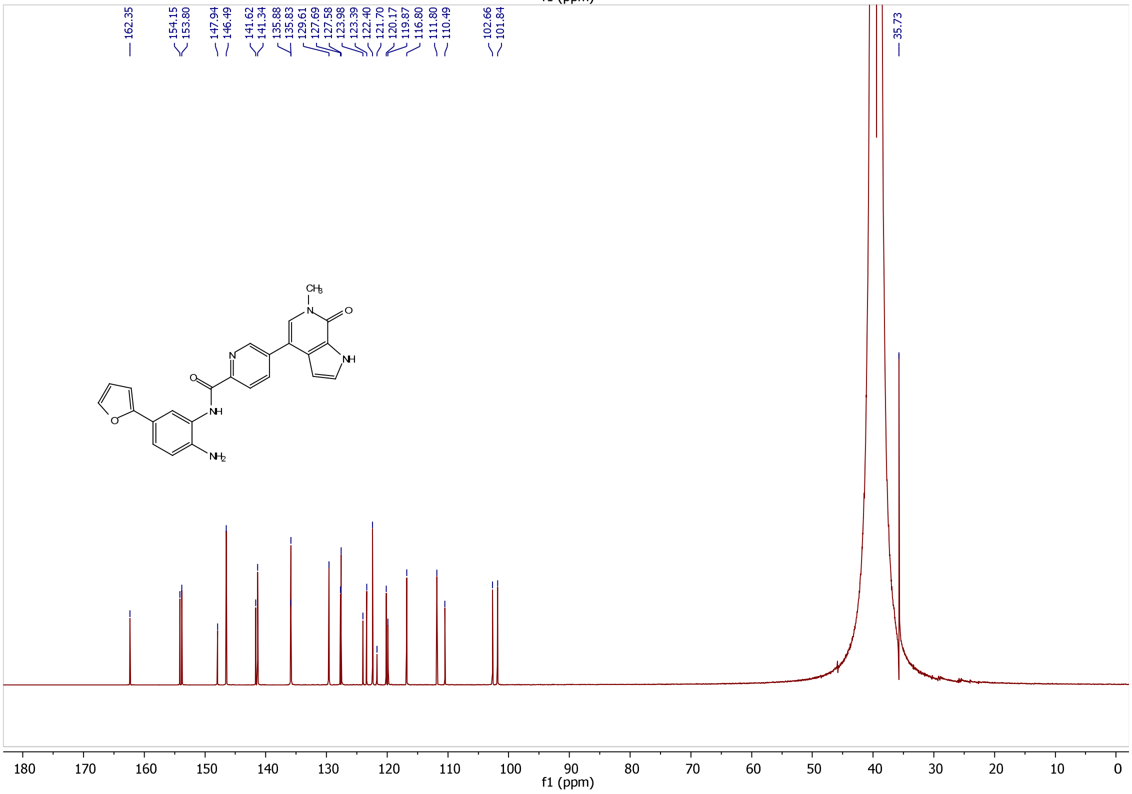

[1]H and [13]C Spectra of NB514 (**39c**)

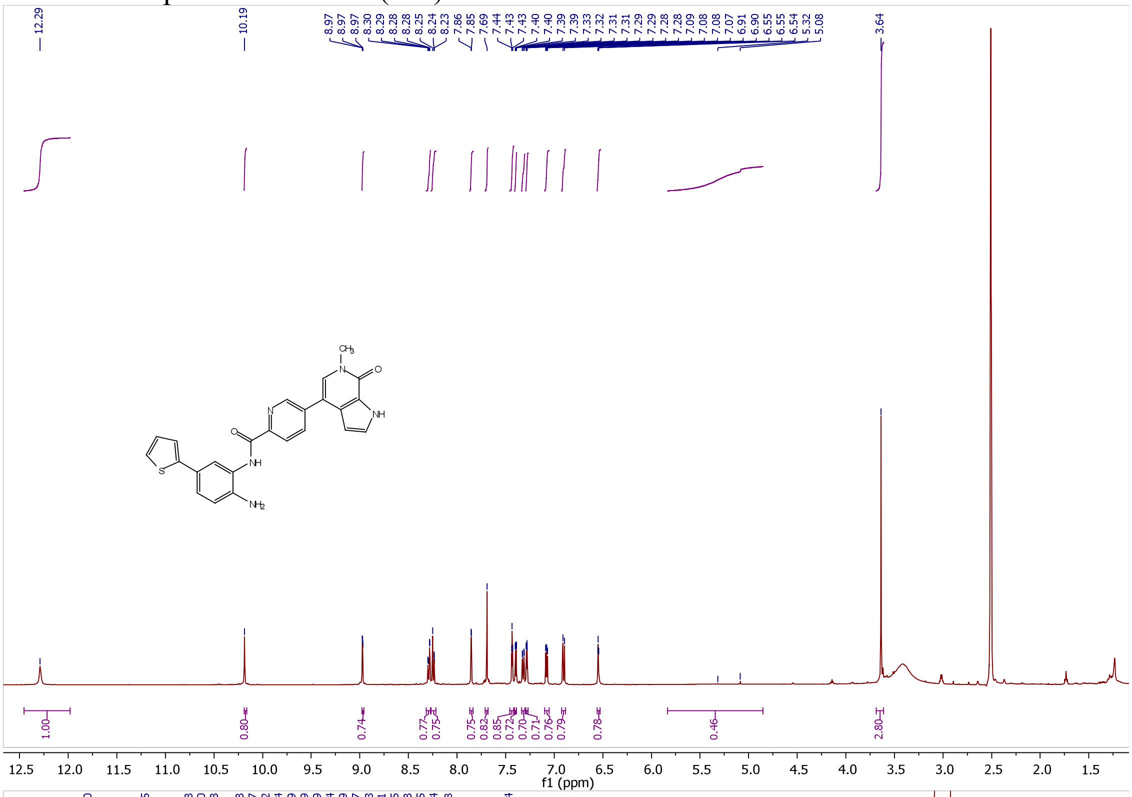

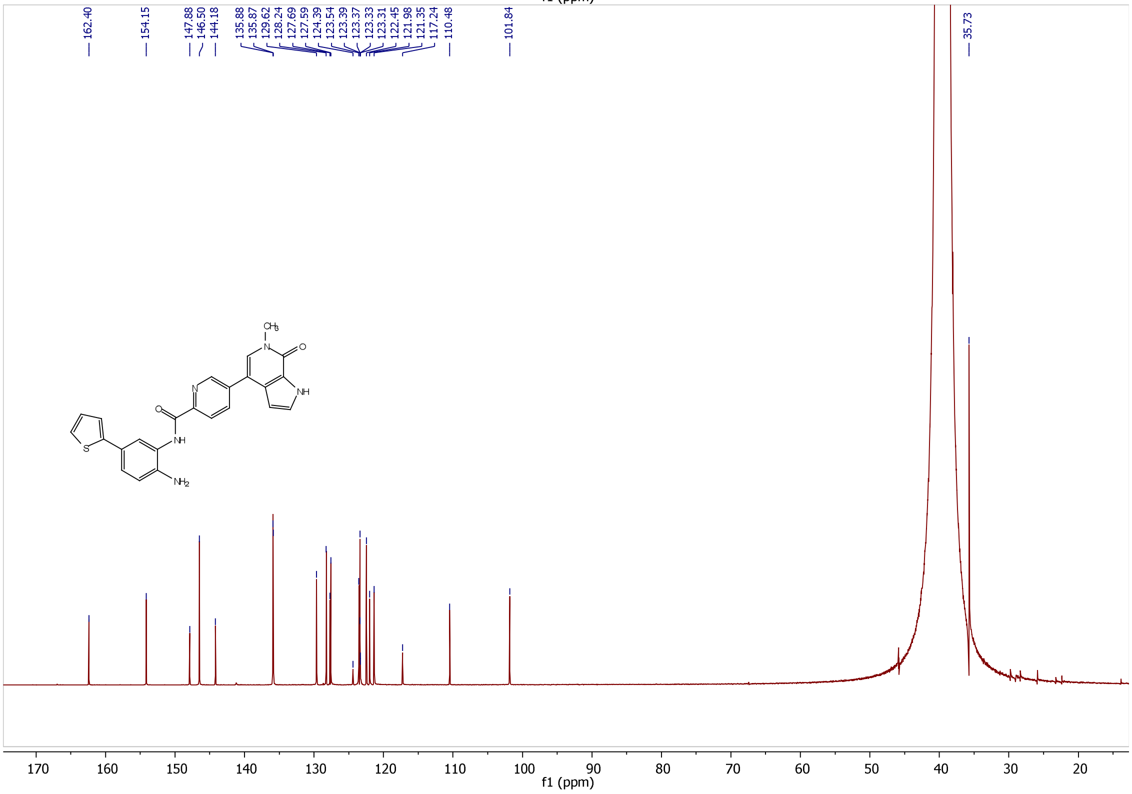
